# Improving The Performance Of The Amber Rna Force Field By Tuning The Hydrogen-Bonding Interactions

**DOI:** 10.1101/410993

**Authors:** Petra Kührová, Vojtěch Mlýnský, Marie Zgarbová, Miroslav Krepl, Giovanni Bussi, Robert B. Best, Michal Otyepka, Jiří Šponer, Pavel Banáš

## Abstract

Molecular dynamics (MD) simulations became a leading tool for investigation of structural dynamics of nucleic acids. Despite recent efforts to improve the empirical potentials (force fields, *ffs*), RNA *ffs* have persisting deficiencies, which hamper their utilization in quantitatively accurate simulations. Previous studies have shown that at least two salient problems contribute to difficulties in description of free-energy landscapes of small RNA motifs: (i) excessive stabilization of the unfolded single-stranded RNA ensemble by intramolecular base-phosphate and sugar-phosphate interactions, and (ii) destabilization of the native folded state by underestimation of stability of base pairing. Here, we introduce a general *ff* term (gHBfix) that can selectively fine-tune non-bonding interaction terms in RNA *ffs*, in particular the H-bonds. gHBfix potential affects the pair-wise interactions between all possible pairs of the specific atom types, while all other interactions remain intact, i.e., it is not a structure-based model. In order to probe the ability of the gHBfix potential to refine the *ff* non-bonded terms, we performed an extensive set of folding simulations of RNA tetranucleotides and tetraloops. Based on these data we propose particular gHBfix parameters to modify the AMBER RNA *ff*. The suggested parametrization significantly improves the agreement between experimental data and the simulation conformational ensembles, although our current *ff* version still remains far from being flawless. While attempts to tune the RNA *ffs* by conventional reparametrizations of dihedral potentials or non-bonded terms can lead to major undesired side effects as we demonstrate for some recently published *ffs*, gHBfix has a clear promising potential to improve the *ff* performance while avoiding introduction of major new imbalances.

## INTRODUCTION

Molecular dynamics (MD) simulations have become a very important tool for studies of biomolecular systems such as nucleic acids with routine access to micro- or even millisecond timescales.^1–7^ MD simulations are often instrumental for understanding and clarifying experimental results and for obtaining a more complete picture of their biological implications. Nevertheless, a sufficiently realistic description of biopolymers by the used empirical potentials (force fields, *ffs*) is essential for successful applications of MD.^4, 8^ The polyanionic nature and high structural variability of ribonucleic acid (RNA) makes the development of RNA *ffs* an especially daunting task.^4^ Despite huge efforts to fix problems that have emerged on ns-μs simulation timescales,^9–17^ RNA *ffs* still cause some behaviors in simulations, which are not consistent with experiments.

Limitations of the available RNA *ffs* have been reviewed in detail, with a suggestion that the currently available pair-additive RNA *ffs* are approaching the limits of their applicability.^4^ Examples of such problems for some recently suggested *ff* versions are documented in the present study. A radical solution of the RNA *ff* problem could be the use of polarizable *ff*s.^18–19^ However, their sophisticated parametrization not only results in extra computational cost but it yet remains to be seen if their accuracy will eventually surpass the best non-polarizable RNA *ff*s in extended simulations. Another option is to augment the existing *ff* forms by some additional simple *ff* terms that could be used to tune the *ff* performance while minimizing adverse side effects. One such *ff* term is introduced in this study.

Compact folded RNA molecules are typically well-described by modern *ff*s on a sub-μs timescale when starting simulations from established experimental structures.^4^ This has allowed many insightful studies on RNAs and protein-RNA complexes. However, the stability of folded RNAs on longer timescales is affected by the free-energy balance between folded, misfolded and unfolded states. Therefore, a common way to identify major problems in *ff*s is using enhanced-sampling techniques, where, generally, either the probability distribution of a limited number of selected degrees of freedom is changed (importance-sampling methods), or the total energy (equivalently, the temperature) of the system is modified (generalized-ensemble algorithms); see Refs. ^20–21^. Considering biomolecular simulations, replica exchange (RE) methods^22–23^ are among the most popular enhanced-sampling techniques. RE simulations profit from multiple loosely coupled simulations running in parallel over a range of different temperatures (T-REMD^22^ simulations) or Hamiltonians (e.g. REST2^23^ simulations). Exchanges between replicas are attempted at regular time intervals and accepted conforming to a Metropolis-style algorithm. RE methods do not require any prior chemical insights regarding the folding landscapes. Obviously, even enhanced-sampling simulations are not *a panacea* and their capability to accelerate sampling has some limits; see Refs. ^4^ and ^24^ summarizing recent applications of enhanced-sampling methods to RNA systems.

The structural dynamics of RNA tetranucleotides (TNs) represents one of the key benchmarks for testing RNA *ff*s.^14-15, 17, 25-32^ TNs are ideal testing systems due to their small size and straightforward comparison of their simulations with solution experiments. Obviously, any quantitative *ff* assessment is critically dependent on the convergence of structural populations because only well-converged simulations can provide unambiguous benchmark datasets. Nevertheless it appears that contemporary simulation methods and hardware already allow to obtain sufficiently converged simulation ensembles for TNs.^14-15, 17, 28-29, 31, 33-34^ TN simulations can specifically evaluate performance of *ff*s for several salient energy contributions, namely, (i) sugar-phosphate (SPh) and base-phosphate (BPh) interactions, (ii) base stacking interactions, (iii) backbone conformations and (iv) balance of these contributions with solvation. Experimental data revealed that TNs mostly populate A-form conformations^25–27^ but MD simulations tend to significantly sample also non-native intercalated structures (or some other non-native structures, Figure 1) that are considered to be a *ff* artifact.^14-15, 17, 27-31, 35^ Obviously, when using TNs as a benchmark for *ff* development, one has to be concerned about a possible over-fitting of the *ff* towards the canonical A-RNA conformation, which may have detrimental consequences for simulations of folded RNAs (see below).

**Figure 1.**
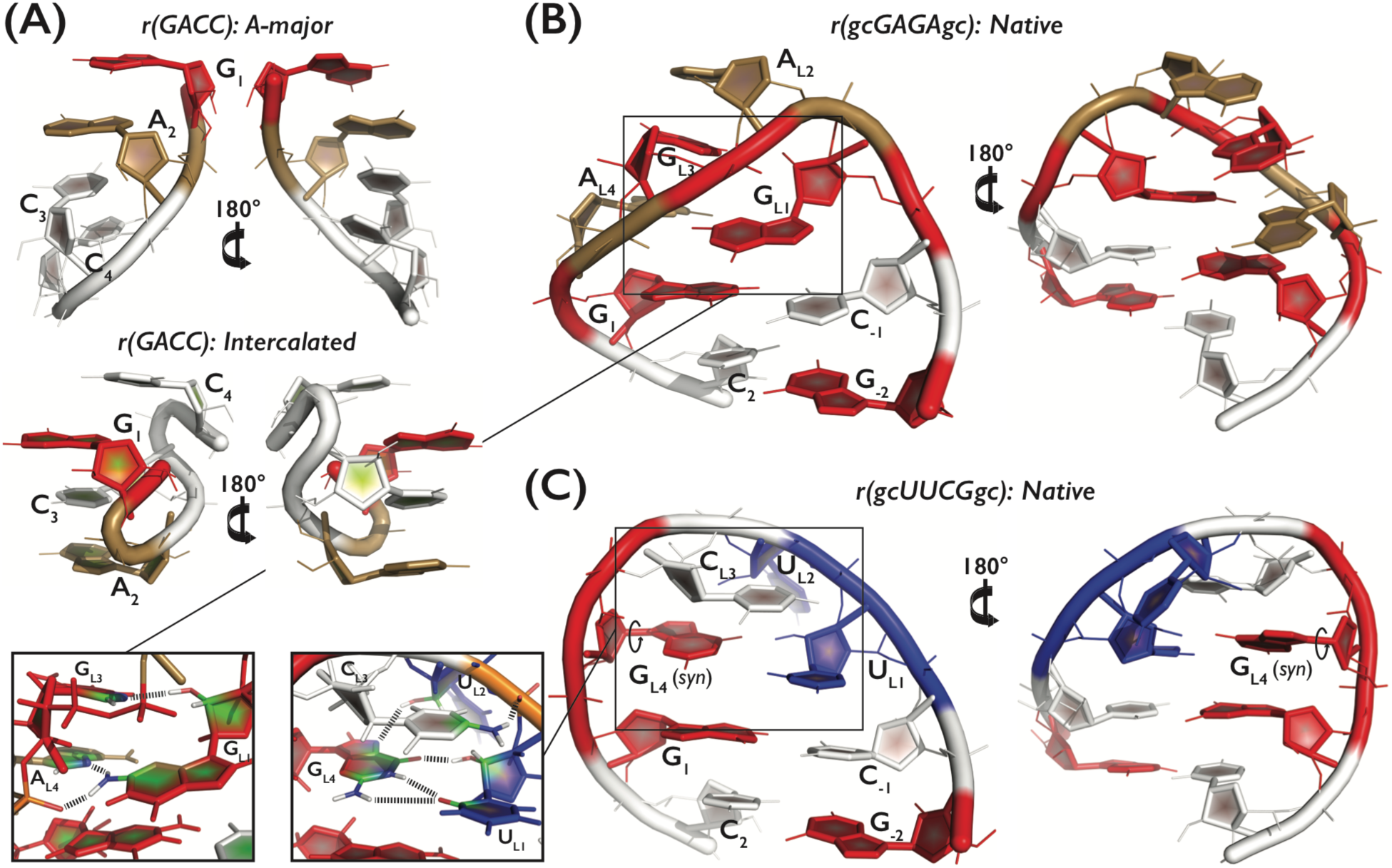
Tertiary structures and detail overview of the three systems: (A) r(GACC) TN in the dominant native A-major conformation (top) and the spurious intercalated structure (bottom), and (B) r(gcGAGAgc) and (C) r(gcUUCGgc) TLs in their native conformations. A, C, G and U nucleotides are colored in sand, white, red, and blue, respectively. The two insets (bottom left) highlight three and five signature H-bonds, i.e., GL1(N2H)…AL4(*pro*-RP), G_L1_(N2H)…A_L4_(N7), and G_L1_(2’-OH)…G_L3_(N7) for the GNRA TL (B) and U_L1_(2’-OH)…G_L4_(O6), U_L2_(2’-OH)…G_L4_(N7), C_L3_(N4H)…U_L2_(*pro*-RP), and bifurcated G_L4_(N1H/N2H)…U_L1_(O2) for the UNCG (C) TL, respectively. Note that all analyses involving occurrence of the native state within generated conformational ensembles considered just four signature H-bonds for the UNCG TL, i.e., we counted only the G_L4_(N1H)…U_L1_(O2) interaction from the bifurcated G_L4_(N1H/N2H)…U_L1_(O2) H-bond.

Another key energy contribution that needs to be described by a *ff* are base-pair interactions. A large part of the thermodynamic stability of folded RNAs is due to formation of canonical A-RNA helices stabilized by canonical Adenine-Uracil (AU) and Guanine-Cytosine (GC) Watson-Crick base pairs and their stacking interactions.^36^ Thus, correct description of canonical base pairs is fundamental for folding studies of RNA molecules. Besides this, RNA molecules contain an astonishing variability of other base-pair patterns.^4, 37^ Assessment of the capability of the *ffs* to describe these interactions requires analyses of more complicated systems like RNA tetraloops (TLs) and folded RNA motifs.^4^ The GNRA and UNCG TLs (N and R stand for any and purine nucleotide, respectively) are the most abundant hairpin loops in RNA molecules.^38–41^ These TLs contribute to various biological functions including tertiary folding, RNA-RNA and protein-RNA interactions, ligand binding, and thus play key roles in transcription, translation and gene regulation.^42–48^ Both above-noted TL types possess a clear dominant folded topology that is characterized by a set of signature molecular interactions that determine the consensus sequences (Figure 1).^39^ The structure of isolated small TL motifs in solution is in dynamic temperature-dependent equilibrium between folded and unfolded conformations. The two main parts of TL’s, i.e., their A-RNA stem and the structured loop itself, contribute differently to the folding free-energy landscape, and thus simulations of TLs allow us to simultaneously probe the capability of the *ffs* to describe A-RNA duplexes as well as some noncanonical interactions and backbone conformations.^49–51^ In contrast to dimerization of A-form duplexes, sampling of the hairpin free-energy landscape is simplified due to unimolecular nature of its folding. TL’s with longer stems formed by canonical base pairs are more stable and thus require higher temperature for melting (unfolding).^50–51^ Therefore, minimal 8-nucleotide long (8-mers) TL motifs with just two base pairs are the preferred targets for computational studies due to their small size and relative ‘metastability’, which is further affected by the nature and orientation of both the closing and terminal base pairs.^50^ Nevertheless, there is a clear experimental evidence that even these short oligomers should be dominantly in the folded state.^52^ The native TL conformations thus represent a genuine benchmark for the *ff* development, which might unambiguously identify underestimated stability of the folded RNA structures in the *ff*s. Such problems are manifested through a complete loss of the characteristic TL structures (Figure 1) in MD simulations; note that reversible fluctuations about the native TL conformations are not sufficient for *ff* verification since the available experimental data do not provide sufficiently unambiguous information about subtle plasticity of the folded TLs. The *ffs* need to be tested also on broad range of other RNA systems.^4^

Recent studies generated large conformational ensembles of TNs^14-15, 17, 28-29, 31, 33-34^ and TLs^14-15, 17, 28-29, 31, 35, 53-56^ in order to assess the performance of RNA *ff*s. They showed that the available RNA *ff*s have persisting deficiencies causing, e.g., (i) shifts of the backbone dihedral angles to non-native values, (ii) problems with the χ-dihedral angle distribution, and (iii) over-populated SPh and BPh hydrogen bond interactions.^15, 25, 55^ Some of those studies also suggested potential directions for *ff* improvement, which included modifications of backbone dihedral terms, van der Waals (vdW) radii and charges, RNA interaction with solvent/ions (adjustments of Lennard-Jones parameters typically accompanied by modifications of the Lennard-Jones combining rules via nonbonded fix, NBfix, to balance the RNA-solvent interaction) and enforced distributions from solution experiments.^14, 17, 29-31, 53, 57-60^ Computer folding of UNCG TLs appears to be much more challenging than folding of GNRA TLs and description of TNs,^15, 35, 53, 55, 61^ as for the latter two systems some partial successes have been reported. Note that there have been repeated past claims in the literature about successful folding of RNA TLs in simulations. However, the claims were not confirmed by independent research groups, as extensively reviewed in Ref. ^4^. The performance and possible shortcomings of another recently released *ff* ^60^ are discussed as part of this work.

Previously, we have shown that at least two different imbalances likely contribute to the (in)correct folded/unfolded free-energy balance of TNs and TLs. The first problem was excessive stabilization of the unfolded ssRNA structure by intramolecular BPh and SPh interactions.^62^ The excessive binding of 2’-hydroxyl groups (2’-OH) towards phosphate nonbridging oxygens (nbO) was reported earlier^27, 58^ and could be partially reduced by using alternative phosphate oxygen parameters developed by Case et al.^63^ in combination with the OPC^64^ explicit solvent water model.^14^ However, the OPC water model was shown to destabilize three-tetrad DNA quadruplex stems^65^ while the modified phosphate parameters were not successful in correcting base-phosphate H-bonding in simulations of Neomycin-sensing riboswitch,^66^ indicating that the modified phosphate parameters with OPC water model do not represent the ultimate solution for tuning the phosphate-group interactions. The second problem was destabilization of the native folded state by underestimation of the native H-bonds including the stem base pairing. As a general correction for under- or overestimation of base-base and other H-bond interactions remains challenging,^4^ we recently introduced a weak local structure-specific short-range biasing potential supporting the native H-bonds (HBfix). Its application led to a substantial improvement of GAGA TL folding^62^ and stabilization of U1A protein-RNA interface.^67^ The HBfix approach, however, does not provide a transferable *ff* that can be applied on systems, where the native structure is not known *a priori*.

In distant past, H-bond specific terms were used both in AMBER^68^ and CHARMM^69^ molecular mechanics (MM) packages for tuning H-bond distances and energies. They were introduced in the form of a modified Lennard-Jones 10-12 potential (AMBER *ff*)^70^ or more sophisticated Lennard-Jones 10-12 or 4-6 potential accompanied by cosine terms improving the directionality of the H-bond (CHARMM *ff*).^71^ Later, the breakthrough paper in *ff* development by Cornell *et al*.^72^ introduced the improved charge model, the restrained electrostatic potential (RESP) fit of charges. The authors claimed that a proper calibration of the point-charge model combined with standard vdW parameters for the AMBER *ff* is entirely sufficient to describe medium-strength H-bond interactions present in biomolecular systems.^73^ Similarly for CHARMM *ff*, Smith and Karplus argued that H-bond specific terms can be sufficiently represented by a balanced combination of electrostatic and vdW interactions.^74^ Thus, in the last two decades, separate terms for H-bonding interactions were deemed unnecessary. However, those conclusions were based on analyses of rather short simulations (sub ns-long time scales), which provided only limited and crude insight into the stability of the H-bond interactions. On the other hand, nowadays, powerful enhanced-sampling techniques and routine accessibility of μs-long time scales allow us to probe the quantitative details of the H-bond interactions. Recently, we have shown that the fine energy balances of the H-bonds significantly affect the conformational preferences between folded and unfolded states of RNA molecules.^53, 62^ These new observations fully justify our attempt to re-introduce some H-bond specific terms into the AMBER nucleic acids *ff* in order to fine-tune its performance. Further, it has been noted that the presently used basic form of the pair-additive *ff* for nucleic acids simulations may be, after the recent parameter refinements, close to its accuracy limits.^4^ Then its further tuning becomes impractical, since improving one simulation aspect deteriorates others. To lift this deadlock, addition of new *ff* terms orthogonal to the common *ff* terms appears as a viable strategy. Finally, also high-quality QM calculations suggest that relying only on the common Lennard-Jones sphere and point-charge model in description of H-bonding has inevitable accuracy limitations.^75–76^

In this work, we introduce a generalized formulation of the HBfix potential, in order to tune all interactions of the same kind, henceforth labelled as gHBfix. Most importantly, the gHBfix correction can be applied without previous knowledge of the native structure. gHBfix can be used for tunable modification of selected nonbonded terms, namely specified types of H-bond interactions, in order to improve the behavior of the current state-of-the-art RNA *ff*.^11, 13, 73, 77^ In the preceding applications, HBfix was used as a native-structure-centered *ff* correction to support known native interactions. In contrast, gHBfix is an interaction-specific *ff* correction, whose application depends only on the atom types. It is not biased in favor of any specific fold or RNA sequence. It can be easily applied and does not require any modification of standard simulation codes.^78–79^ We used enhanced-sampling methods and obtained a large amount of data (total simulations timescale more than 4 ms) during the testing phase with several variants of the gHBfix parameters. We mainly focused on structural dynamics and folding of TN and TL systems with the aim to generate ensembles with population of major conformers close to those reported by experimental datasets.^50–52^ The specific variant of the gHBfix potential suggested based on the TN and TL simulations was subsequently tested on various important RNA structural motifs using unbiased MD simulations. Although the suggested *ff* version certainly does not eliminate all the *ff* problems, as discussed in detail, it brings valuable improvements so far without visible side effects. We further propose that the gHBfix methodology has a substantial potential for further tuning.

## METHODS

### Starting structures and simulation setup

The starting structures of r(gcGAGAgc) and r(gcUUCGgc) (in unfolded states) and r(GACC) as well as the other four TNs were prepared using Nucleic Acid Builder of AmberTools14^80^ as one strand of an A-form duplex. The topology and coordinates were prepared using the tLEaP module of AMBER 16 program package.^78^ Single strands were solvated using a rectangular box of OPC^64^ water molecules with a minimum distance between box walls and solute of 10 Å, yielding ~2000 water molecules added and ~40×40×40 Å^3^ box size for TN and ~7000 water molecules added and ~65×65×65 Å^3^ box size for both TLs, respectively. Simulations were run at ~1 M KCl salt excess using the Joung-Cheatham ion parameters^81^ (K^+^: *R* = 1.590 Å, *ε* = 0.2795 kcal/mol, Cl^−^: *R* = 2.760 Å, *ε* = 0.0117 kcal/mol). We used the *ff*99bsc0χ_OL3_^11, 13, 73, 77^ basic RNA *ff* version with the vdW modification of phosphate oxygen developed by Case et al.^63^. These phosphate parameters for phosphorylated aminoacids were shown to improve the performance of RNA *ff*.^14, 62^ All the affected dihedrals were adjusted as described elsewhere.^58^ AMBER library file of this *ff* version can be found in Supporting Information of Ref. ^62^. Notice that although the OPC water model was used in this study, the derived gHBfix parameters should be applicable also with other water models, though one can expect that for some RNA systems the simulation outcomes may moderately depend on the used water models. We think that it could be possible in future to couple optimization of gHBfix parameters with specific water models. This would, however, be justified when being capable to test a finer “grid” of the gHBfix parameters, e.g., with the help of reweighting parametrization procedures. Based on the available literature data we note that although a choice of the water model may have some influence on RNA simulations of specific systems, it is not a decisive factor.^82–84^ In addition, we do not think it would be appropriate to suggest one water model as being universally best for RNA simulations. In summary, the RNA *ff* version primarily used here is composed of: *ff*99bsc0χ_OL3_ basic RNA *ff*11, 13, 73, 77 combined with (i) the phosphate oxygens by Case et al.,^63^ (ii) the ion parameters by Joung&Cheatham,^81^ and (iii) the OPC water model.^64^ For the sake of simplicity, this RNA *ff* is abbreviated as χ_OL3CP_ through the text and is further tuned by our external gHBfix potential.

### REST2 Settings

The replica exchange solute tempering (REST2)^23^ simulations of r(GACC) TN, r(gcGAGAgc), and r(gcUUCGgc) TLs were performed at 298 K with 8 (TN) and 12 (TLs) replicas. Details about settings can be found elsewhere.^62^ The scaling factor (λ) values ranged from 1 to 0.6017 (TN) or to 0.59984 (TLs) and were chosen to maintain an exchange rate above 20%. The effective solute temperature ranged from 298 to ~500 K. The hydrogen mass repartitioning^85^ with a 4-fs integration time step was used. The standard length of REST2 simulations was 10 μs per replica (specific tests were terminated earlier; see Table S1 in Supporting Information for summary of all enhanced-sampling simulations); the cumulative time of all REST2 simulations was ~4.2 ms.

### gHBfix for support/weakening of specific interactions

The main aim of this work is extension of the previously introduced structure-specific HBfix potential to the generalized interaction-specific gHBfix potential (Figure 2), creating essentially new generally applicable RNA *ff* versions. The potential function is defined as:

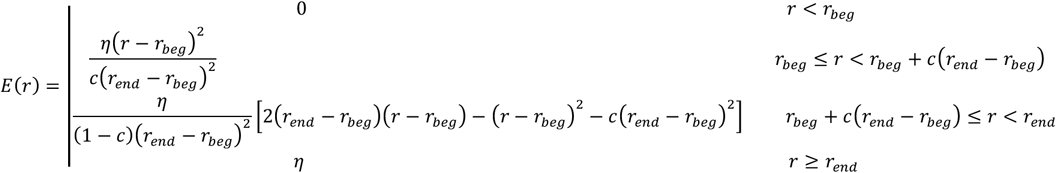

**Figure 2.**
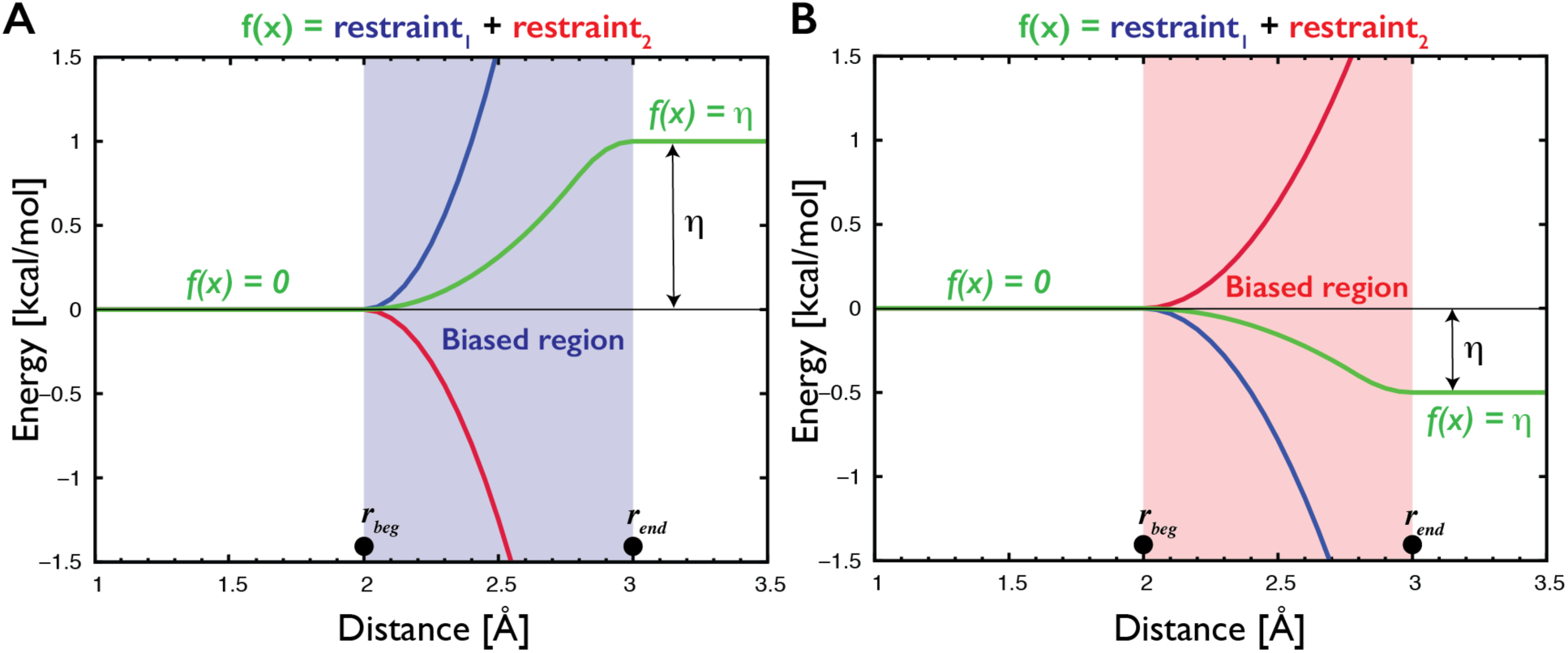
Description of the gHBfix potential (green curves) used for either support (A) or weakening (B) of H-bond interactions. In the present study, gHBfix potential is in all cases applied to the distance between hydrogen of the proton donor and proton acceptor heavy atom. The potential is constant (i.e., it provides zero forces) at all distances except for the narrow region between *r_beg_* and *r_end_* corresponding roughly to the expected location of the free energy barrier between bound and unbound states. For the used distance between hydrogen and proton acceptor heavy atom of the hydrogen bond we applied here 2 Å and 3 Å for *r_beg_* and *r_end_*, respectively. The potential is formally composed of a combination of two flat-well restraints with opposite sign of curvature and linear extensions that cancel each other at distances above *r_end_* (red and blue curves). The η parameter defines total energy support (A) or penalty (B) for each H-bond interaction. Although the gHBfix potential is technically constructed using two restraining functions, it is not a restraint; we have used this technical solution solely in order to be able to efficiently use common functions that are available in the simulation codes without any necessity to modify them.

where *r_beg_* and *r_end_* delimit the region, in which the potential is not constant, *c* defines the position of the point of inflection, so that its position is *rbeg + c(r_end_ - r_beg_)*, and the *η* is a controllable parameter, which defines total energy bias (in kcal/mol) for the H-bonds. The potential was originally applied to heavy atom distances of H-bonds, i.e., between proton donor and proton acceptor heavy atoms.^62^ Here we applied it between hydrogen of the proton donor and proton acceptor heavy atom.

Originally, the HBfix potential was used to support native H-bonds in a structure-specific manner that was sufficient to achieve folding of the r(gcGAGAgc) TL.^62^ For the sake of completeness, we report equivalent structure-specific HBfix T-REMD folding simulation of r(gcUUCGgc) TL as part of this work. However, the main focus of this study was the development of the gHBfix potential, where all possible interactions between specific groups are affected. We have tested a number of variants and combinations of the gHBfix potential. The list of all RNA groups and atoms involved in the gHBfix is summarized in Tables 1 and 2 (see Table S1 for summary of performed REST2 simulations with particular gHBfix settings). For the clarity, each tested setting of the gHBfix potential is marked as 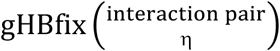, where the upper label in the parentheses defines the specific interactions between RNA groups (Table 2) and the lower parameter η defines the total energy bias for each H-bond interaction of this kind (in kcal/mol, Figure 2).

**Table 1.** The list of groups and atoms from RNA nucleotides whose interactions were modified by the gHBfix.

| Donors | Nucleotides |  |  |  |
| --- | --- | --- | --- | --- |
|  | A | C | G | U |
| NH (base) | N6(H61, H62) | N4(H41, H42) | N1(H1), N2(H21, H22) | N3(H3) |
| 2-OH (sugar) | O2'(HO2') | O2'(HO2') | O2'(HO2') | O2'(HO2') |
| Acceptors |  |  |  |  |
| N (base) | N1, N3, N7 | N3 | N3, N7 | - |
| O (base) | - | O2 | O6 | O2, O4 |
| O (sugar) | O4', O2' | O4', O2' | O4', O2' | O4', O2' |
| bO (phosphate) | O3', O5' | O3', O5' | O3', O5' | O3', O5' |
| nbO (phosphate) | *pro*-R<sub>P</sub>, *pro*-S<sub>P</sub> | *pro*-R<sub>P</sub>, *pro*-S<sub>P</sub> | *pro*-R<sub>P</sub>, *pro*-S<sub>P</sub> | *pro*-R<sub>P</sub>, *pro*-S<sub>P</sub> |

**Table 2.** All possible combinations of groups used for various gHBfix settings and their abbreviations. See Table 1 for atoms involved within each group.

| gHBfix name | Donors | Acceptors |
| --- | --- | --- |
| 2-OH...nbO | 2'-OH (sugar) | nbO (phosphate) |
| 2-OH...bO | 2'-OH (sugar) | bO (phosphate) |
| 2-OH...O2 | 2'-OH (sugar) | O2' (sugar) |
| 2-OH...O4 | 2'-OH (sugar) | O4' (sugar) |
| 2-OH...N | 2'-OH (sugar) | N (base) |
| 2-OH...O | 2'-OH (sugar) | O (base) |
| NH...N | NH (base) | N (base) |
| NH...O | NH (base) | O (base) |
| NH...O2 | NH (base) | O2' (sugar) |
| NH...O4<sup>a</sup> | NH (base) | O4' (sugar) |
| NH...bO<sup>a</sup> | NH (base) | bO (phosphate) |
| NH...nbO<sup>a</sup> | NH (base) | nbO (phosphate) |

<sup>a</sup> Not tested in this work.

### Conformational analysis

Native states of r(gcGAGAgc) and r(gcUUCGgc) TLs were determined based on the presence of all native H-bonds, i.e., those participating on the base pairing in the stem as well as signature interactions of the loop (presence of a H-bond was inferred from a hydrogen-acceptor distance within cutoff 2.5 Å), in combination with the ***ε***RMSD metric.^86^ The dominant conformations sampled in REST2 simulations were identified using a cluster analysis based on an algorithm introduced by Rodriguez and Laio^87^ in combination with the ***ε***RMSD.^86^ The algorithm was introduced in Ref. ^62^ and uses the idea that cluster centers are characterized by a higher local density of points than their neighbors and by a relatively large distance from any other points with higher local density. In order to apply the clustering algorithm with the *ε*RMSD metric,^86^ cluster selection was customized. The point having highest product of the local density and the distance to the nearest point of higher density was assigned as a next cluster center. Subsequently, the other points were assigned to this cluster center based on two rules: their nearest neighbor of higher density is (i) also member of the same cluster and is (ii) closer than 0.7 of *ε*RMSD (see Supporting Information of the Ref. ^62^ for details about implementation of the algorithm). This process was repeated until all points were processed and clusters with population above 1% were further analyzed. The cluster analysis was performed for unscaled replicas (λ=1, T = 298 K).

The estimations of error bars were calculated by bootstrapping^88^ using the implementation introduced in our recent study.^62^ The estimations of errors were obtained by resampling of the time blocks of the whole set of replicas and subsequent resampling of the coordinate-following replicas for the purpose of their demultiplexing to follow Hamiltonians in order to obtain a final resampled population at the reference unbiased Hamiltonian state (see Figure S1 in Supporting Information for more details). In time domain, we used a moving block bootstrap where optimal size of the blocks was selected to be close to *n*^1/3^, where *n* is the number of snapshots.^89^ Subsequently, the continuous replicas following coordinates were resampled (with replacement) and this resampled set of continuous replicas was subsequently demultiplexed, i.e., resorted to follow the Hamiltonians. The conformational ensemble was analyzed at each level of Hamiltonian bias (including the reference unbiased Hamiltonian) from this set of replicas. For each of the 1000 resample trials, we calculated the population of folded and misfolded states at each level of Hamiltonian. Finally, the confidence interval of these populations with a 5% level of significance was calculated from all 1000 resample trials. We would like to note that the bootstrapping is typically implemented in the literature as resampling in the time-domain only. Such simple resampling of time blocks might be used to estimate uncertainties originating from the limited sampling in the time domain. However, the more sophisticated bootstrap implementation used in this study allows us in addition to clearly distinguish between cases where the native state is formed only in one or very few replicas versus the native state being repeatedly formed in most or all replicas. Although in both cases one would observe both folded and unfolded structures in the reference replica, the latter scenario corresponds to populations that can be expected to be significantly closer to convergence, since multiple replicas contribute to the population of the native state. As shown in our previous study,^62^ the above-described resampling of replicas takes this effect into account and thus provides a considerably more conservative estimate of the convergence than the commonly used plain bootstrapping (see Figure S1 in the Supporting Information). The example of data analysis with error bars calculated only by bootstrap resampling in time-domain is shown on Figure S2 in the Supporting Information.

### Comment on convergence analysis of the TL REST2 simulations

RE simulations are converged once all replicas sample the same ensemble.^15^ Such ultimate convergence occurs typically on a much longer time-scale than the time-scale needed to reach an apparently steady-state population on the reference replica. Thus, seemingly converged steady-state population of the folded state as revealed by reference replica might still contain some statistical inaccuracy if the folding events were observed only in some replicas (see Supporting Information for details). Note that this is the case of the present REST2 simulations of TLs, where the full convergence is still beyond what is achievable for current computers. On the other hand, the bootstrapping implementation used in this study with resampling both in time- and replica-domain provides a fair estimate of uncertainties stemming from such still rather limited convergence. We further note that folding simulations performed in different *ff*s might require different simulation time to achieve the same level of convergence. In conclusion, we caution that with presently available computer resources and enhanced-sampling algorithms, quantitatively converged simulations of RNA TLs are not achievable; this unavoidable uncertainty obviously complicates fine-tuning of the RNA *ff*s.

### Data analysis

All trajectories were analyzed with the PTRAJ module of the AMBER package^78^ and the simulations were visualized using a molecular visualization program VMD^90^ and PyMOL^91^.

### Additional simulations

Besides the REST2 simulations of different variants of our gHBfix potential, we report also a number of other simulations for various RNA systems, to check the performance of the modified *ff*. The cumulative time of all unbiased simulations was ~100 μs and for space reasons, methodological details of these simulations are given in Supporting Information.

### Simulations with other RNA *ffs*

As part of our study, we tested the recently published *ff* by D. E. Shaw and co-workers.^60^ We applied this *ff* for folding simulations of RNA TLs with short A-RNA stems as well as for standard simulations of selected RNA motifs. We observed major instabilities caused by this *ff* version in folded RNAs. In addition, we tested the recently published vdW/NBfix RNA *ff* modification by Pak and co-workers.^31^ We also spotted major problems of this *ff* in description of noncanonical interactions within the Kink-turn RNA motif during standard MD simulations. Methodological details of both tested *ff*s are given in the Supporting Information of the article.

## RESULTS AND DISCUSSION

In the present study, we have attempted to parameterize the gHBfix correction for the χOL3CP^13, 73, 77^ AMBER RNA *ff*. Our primary training systems were r(GACC) TN and two TL hairpins, namely r(gcGAGAgc) and r(gcUUCGgc) (see Figure 1 for structures). Our goal was to improve performance of the simulations for these systems while avoiding undesired side effects for other RNA systems. We have used extended REST2 simulations to generate sufficiently converged conformational ensembles of the above small RNA systems while testing various combinations of the gHBfix potentials acting on specific types of H-bonds (see Methods and Table S1 in Supporting Information for overview of REST2 simulations). The obtained results were compared with the available experimental data.^25-27, 50-52^ We have found a promising gHBfix correction that decisively improves behavior of the r(GACC) TN and r(gcGAGAgc) TL. We did not detect any side effects so far in standard simulations of a wide range of other RNA structures. This indicates that although the suggested modification is not robust enough to fold the r(gcUUCGgc) TL and is less convincing for some other TNs, it provides a significant improvement of RNA simulations when added to the widely used χ_OL3CP_ RNA *ff*. We reiterate that the primary goal of the paper was introduction of the basic principles of the gHBfix potential and demonstration of its capability for improving RNA simulations.

### Weakening of the over-stabilized SPh interactions

According to the NMR data,^25–27^ RNA TNs adopt mostly two different conformations, A-major and A-minor (Figure S6 in Supporting Information). However, recently published standard MD and replica exchange simulations^14-15, 17, 27, 29, 31, 33-34^ using the χ_OL3CP_^11, 13, 73, 77^ RNA *ff* reported unsatisfactory populations of canonical A-form conformations and, instead, significant population of artificial intercalated structures (Figures 1, 3, and Figure S6 in Supporting Information) that are stabilized by SPh and BPh interactions.^27^ Overstabilization of the BPh and especially SPh interactions was repeatedly suggested in our recent studies.^58, 62^ Among all TNs for which benchmark experimental data is available,^25–27^ i.e., r(GACC), r(CAAU), r(CCCC), r(UUUU), r(AAAA), the r(GACC) sequence was most frequently used in the preceding *ff* testing, therefore, we primarily focused on this system, while the other sequences were tested afterwards.

In an attempt to eliminate the unsatisfactory MD behavior, we first designed a negative (destabilizing) gHBfix potential to probe the effect of destabilization of SPh interactions, i.e., H-bonds between all 2’-OH groups and phosphate nbOs (*pro*-RP, *pro*-SP), bOs (O3’, O5’) and sugar O4’ oxygens (Figure 3 and Tables 1 and 2). Several variants of the negative gHBfix were tested using a set of REST2 simulations of r(GACC) TN used as the training system. The results from clustering analysis show that all introduced negative gHBfix potentials acting on SPh interactions increased significantly the native RNA A-major population and essentially eliminated the presence of the intercalated structures (observed populations of artificial intercalated structures were marginal, typically ~1.0 %, Table 3). However, as a side effect, the canonical RNA A-minor conformation (characterized by the change of the α backbone dihedral at the 3’-end, allowing formation of the SPh contact, Figure S6 in Supporting Information) appeared to be destabilized and could even be eliminated when the repulsive gHBfix correction was too strong. For further tests, we took the 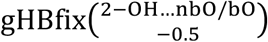 combination, which provides close agreement with experiments,^25^ i.e., χ^2^ value with respect to NMR observables of ~0.15 and populations of A-major/A-minor/Intercalated structures around 65%/18%/0%, respectively (Table 3). The chosen 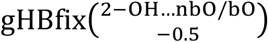 setting consists of the negative gHBfix applied to all possible interactions between 2’-OH groups as proton donors and nbO and bO oxygens as acceptors with the destabilization constant Ƞ of –0.5 kcal/mol. It is notable that comparable improvement over the control χ_OL3CP_ simulation was achieved by other tested potentials, especially those involving interactions between 2’-OH groups and nbO oxygens (Table 3), suggesting the possibility of further tuning of the gHBfix parameters for SPh interactions.

**Table 3.** Analysis of the most populated structural clusters (in %) obtained from r(GACC) REST2 simulations at the reference replica (T = 298 K) with five variants of the gHBfix parameters.^a^

| Cluster | Exp.<sup>b</sup> | Ref.<sup>c</sup> | Population (%) |  |  |  |  |
| --- | --- | --- | --- | --- | --- | --- | --- |
| | | | $\text{gHBfix}^{(2\text{-OH}\dots\text{nbO})}_{-1.0}$ | $\text{gHBfix}^{(2\text{-OH}\dots\text{nbO/bO})}_{-1.0}$ | $\text{gHBfix}^{(2\text{-OH}\dots\text{nbO/bO/O4})}_{-1.0}$ | $\text{gHBfix}^{(2\text{-OH}\dots\text{nbO})}_{-0.5}$ | $\text{gHBfix}^{(2\text{-OH}\dots\text{nbO/bO})}_{-0.5}$ |
| A-major | $\sim 65$ | $36.9 \pm 2.8$ | $69.2 \pm 0.5$ | $76.5 \pm 3.2$ | $61.3 \pm 4.3$ | $65.1 \pm 2.5$ | $58.7 \pm 2.1$ |
| A-minor | $\sim 18$ | $14.4 \pm 1.5$ | $4.4 \pm 0.5$ | $3.7 \pm 0.4$ | $2.8 \pm 0.5$ | $7.7 \pm 0.9$ | $7.4 \pm 1.3$ |
| Intercalated<sup>d</sup> | 0 | $\sim 9$ | $\sim 0$ | $\sim 0$ | $\sim 0$ | $\sim 1$ | $\sim 1$ |
| $\chi^2$<sup>e</sup> | - | 0.75 | 0.14 | 0.18 | 1.27 | 0.13 | 0.15 |

<sup>a</sup> Clustering was performed from the last 7 $\mu\text{s}$ of the 10- $\mu\text{s}$ -long trajectory.
<sup>b</sup> Experimental populations obtained by NMR refinement simulations<sup>25</sup> and by reweighting extensive MD simulations based on these NMR data.<sup>33</sup>
<sup>c</sup> Present control $\chi_{OL3CP}$ simulation without any external potential (see Methods for the detailed setup).
<sup>d</sup> See Figures 1, 2 and Figure S6 in Supporting Information for examples of intercalated structures.
<sup>e</sup> $\chi^2$ values were obtained as described elsewhere<sup>33</sup> by comparing calculated and experimental backbone $^3J$ scalar couplings, sugar $^3J$ scalar couplings, nuclear Overhauser effect intensities (NOEs), and the absence of specific peaks in NOE spectroscopy (uNOEs). NOEs and uNOEs were calculated as averages over the $N$ samples, i.e.,
$(u)NOE_{CALC} = \left( \frac{\sum_i^N r_i^{-6}}{N} \right)^{-1/6}$ . The calculated NOEs were directly compared against experimental values. Each uNOE signal was considered as violated, when the calculated value from the simulation was shorter than the assigned one from the experiment. $^3J$ scalar couplings were obtained from the Karplus relationships (see Supporting Information of the Ref.<sup>33</sup> for the details and experimental datasets).

**Figure 3.**
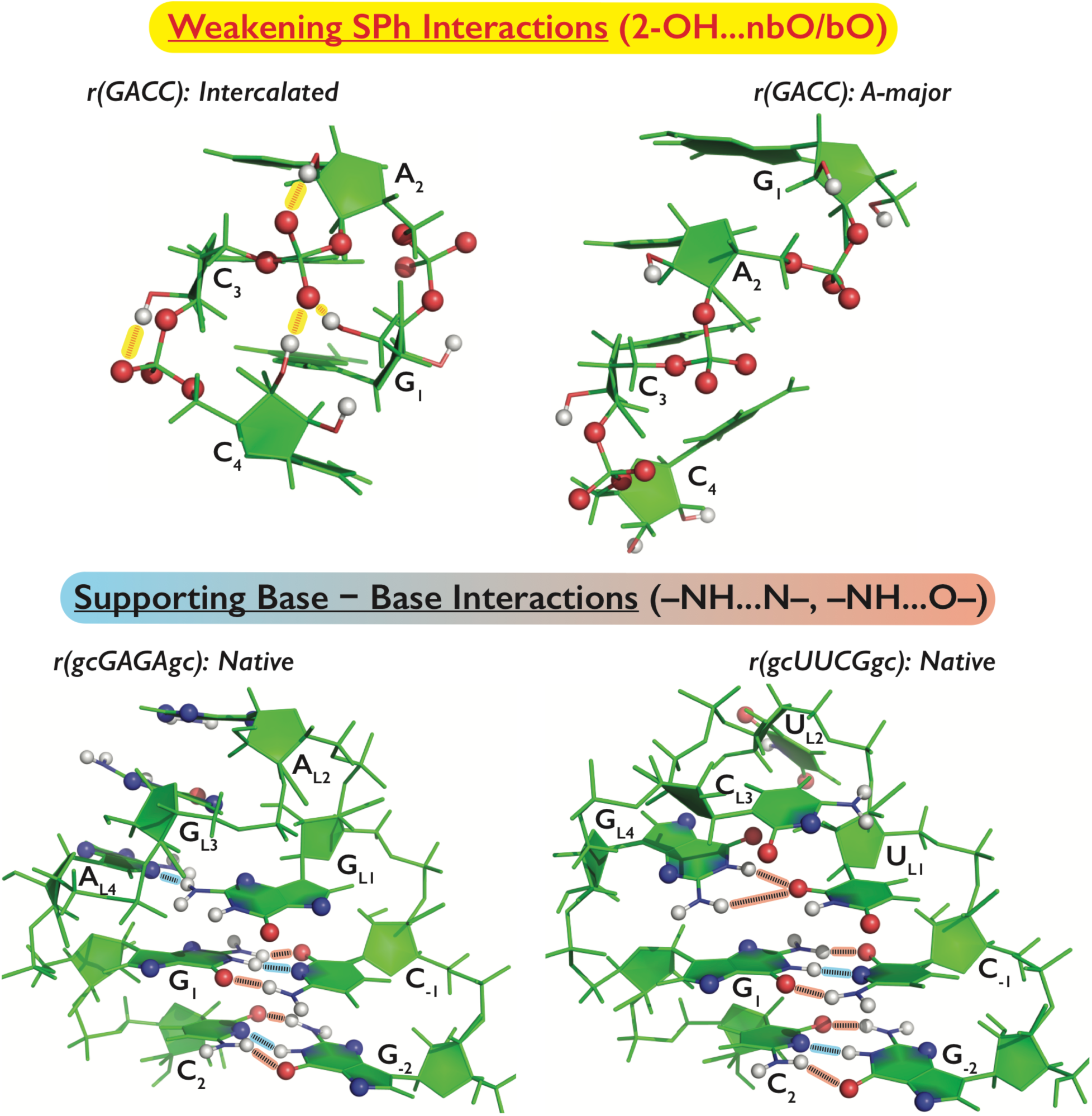
Application of the gHBfix potential and its impact on tertiary structures of the three main test systems, i.e., r(GACC) TN in the spurious intercalated structure (top left) and the dominant native A-major conformation (top right), and r(gcGAGAgc) (bottom left) and r(gcUUCGgc) (bottom right) TLs in their native conformations. Groups included in the gHBfix potential are highlighted by spheres (H, N, and O atoms in white, blue, and red, respectively). The particular version of the gHBfix depicted in the Figure is tuning H-bond interactions in order to: (i) destabilize sugar – phosphate interactions (red dashed lines highlighted by yellow background on the top left panel) and, simultaneously, (ii) support base – base interactions (lower panels; black dashed lines highlighted by blue and red background for –NH…N– and – NH…O– interactions, respectively). Note that the gHBfix potential is affecting all interactions of the same type, i.e., not only those present in the depicted conformations.

### Stabilization of the base-base H-bonding interactions

Short TL motifs are ideal model systems for assessing the folding capability of *ff*s due to their small size, clearly defined native conformation and the possible variety of other competing conformations. We initially applied the 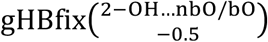 potential in r(gcGAGAgc) REST2 folding simulation but we did not obtain native states, i.e., neither the stem nor the loop sampled native conformations for the entire 10 λs of the REST2 simulation (Table 4). Nonetheless, the 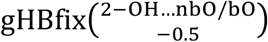 was able to (at least partially) suppress condensed (‘random-coil’) states with excess of BPh and SPh interactions in favor of A-form single stranded structures (see Figure S7 in Supporting Information for geometries and populations of major clusters during the r(gcGAGAgc) REST2 simulations). The persisting inability to sample the native state was not surprising because the problem with TL folding in χ_OL3CP_ RNA *ff* was deemed to be connected with two separate problems, namely, spurious stabilization of unfolded states by excessive BPh and SPh interactions and underestimation of the native H-bonds within the stem (canonical base-pairing interactions) and the loop (signature BPh and other interactions involving the 2’-OH groups, Figure 3).^55, 62^ Recently, we have shown that addition of structure-specific HBfix potential supporting native H-bonds results in satisfactory r(gcGAGAgc) TL folding.^62^ This suggests that the base-pairing interactions should be additionally stabilized, which is also consistent with several recent studies on DNA and RNA guanine quadruplexes and their folding intermediates stabilized by GG Hoogsteen base pairs.^92–94^ In addition, the usefulness of stabilization of base – base H-bonds is also indicated by potentially excessive fraying of base-paired segments occasionally seen in MD simulations of folded RNAs.^4^ In an attempt to fix the base-base interactions in a general fashion, i.e., to support both canonical and noncanonical base pairing, we added additional terms to the gHBfix potential, which support H-bonds between any proton donor from nucleobases (–NH groups) and nucleobase proton acceptors (N/O– atoms, see Figures 2, 3 and Methods for details). We would like to reiterate that these gHBfix potentials are biased neither towards any specific type of base pairing^95^ nor towards any specific pairs of nucleobases in the sequence, so they work irrespective to the base pairing in the native state. They are as general as all other *ff* terms and can be in principle transferred to other systems.

**Table 4.** Tuning of the gHBfix potential for base pairing and its effects on the folding of r(gcGAGAgc) TL. Populations (in %) of two major conformations, i.e., the native state (with properly folded stem and loop) and all states with folded A-form stem (independent of the loop conformation), are displayed for each gHBfix combination at the reference REST2 replica (T = 298 K). See Figure S7 for the examples of most populated clusters from REST2

| (-NH...N-)<br>interaction bias (kcal/mol) | (-NH...O-) interaction bias (kcal/mol) |  |  |
| --- | --- | --- | --- |
|  | (NH...O)<br>+0.0 | (NH...O)<br>+0.5 | (NH...O)<br>+1.0 |
| (NH...N)<br>+1.0 | 20.0/33.2<sup>b</sup> | 40.3/72.1 | 0.0/5.6 |
| (NH...N)<br>+0.5 | 5.1/6.9 | 9.1/26.4 | 22.1/53.6 |
| (NH...N)<br>+0.0 | 0.0/0.0<sup>c</sup> | 0.0/3.9 | 0.0/29.1 |

<sup>a</sup> Cluster analysis performed for the last 7 μs from the 10- μs-long trajectory at 298 K; the gHBfix $\left(\begin{smallmatrix} 2-\text{OH}\dots\text{nbO}/\text{bO} \\ -0.5 \end{smallmatrix}\right)$ potential applied in all simulations.
<sup>b</sup> The numbers before and after “/” report population of native state with all signature interactions and all states with folded A-form stem, respectively. The first population is thus inherently included in the second.
<sup>c</sup> REST2 simulation with the gHBfix $\left(\begin{smallmatrix} 2-\text{OH}\dots\text{nbO}/\text{bO} \\ -0.5 \end{smallmatrix}\right)$ potential affecting only Sph interactions.

We have tested several variants of the base–base gHBfix function, by using a combination of separate terms for (–NH…N–) and (–NH…O–) H-bonds, each with three different values of the gHBfix bias constant Ƞ, namely 0.0, +0.5, and +1.0 kcal/mol. Thus, we performed nine r(gcGAGAgc) TL REST2 simulations with the 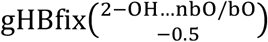 term introduced above for SPh interactions and various combinations of the 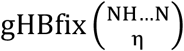 and 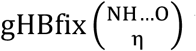 L base-base terms (Table 4). Subsequently, we used clustering analysis to estimate the population of the native state (see Methods). It turned out that additional support of (–NH…N–) H-bond interactions (with Ƞ = +1.0 kcal/mol) is crucial in order to promote significant population of the folded stem and to stabilize the native arrangement of the TL (Figure 4 and Table 4). This is in agreement with the known *ff* imbalance, where N atoms tend to have too large vdW radii, forcing the (–NH…N–) H-bonds to fluctuate around larger distances with respect to Quantum Mechanical (QM) calculations and experimental datasets.^50-52, 96^ We observed that a sufficiently strong support of (–NH…O–) H-bonds (with Ƞ equal to +1.0 kcal/mol) could also stabilize folded stem and/or native stem/loop even in combination with weaker (–NH…N–) support. This likely compensates for the effect of weaker (–NH…N–) interactions; see the r(gcGAGAgc) REST2 simulation with the 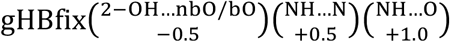 combination (Figure 4 and Table 4). Nonetheless, in such case the population of the native loop conformation is suboptimal and this rather heavy gHBfix combination leads already to some side effects in simulations of TNs (e.g., the r(GACC) TN tends to form non-native loop-like structures with too strong stabilization of base-base interactions, see Supporting Information for details). It also leads to a significant population of the left-handed Z-form helix conformation^97^ (stem guanines in *syn* orientation) instead of the dominant A-form in the r(gcGAGAgc) system (Figure 4). In general, it is advisable to keep the gHBfix *ff* corrections as mild as possible. Despite the fact that r(gcGAGAgc) REST2 simulations are not fully converged (Figure 4 and Figure S5 in Supporting Information), the data strongly suggests applying the positive biases (strengthening) on the (– NH…N–) H-bonds rather than on the (–NH…O–) H-bonds. This observation was also confirmed by nine REST2 simulations of r(GACC) TN differing in base-base gHBfix terms (using the same parameter combinations as tested for the r(gcGAGAgc) TL folding), all with 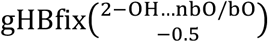 weakening SPh interactions (Table 5). Hence, REST2 simulations with both r(GACC) TN and r(gcGAGAgc) TL show that the 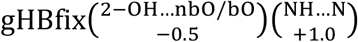 potential represents currently the best compromise (among those variants tested) for tuning H-bonding interactions with a correct balance between different conformers according to the experiments.^25,27, 50^

**Table 5.** The effect of gHBfix potential for base pairing on structural dynamics of r(GACC) TN. Populations (in %) of two major conformations, i.e., RNA A-major (the first number) and A-minor (the second number), are displayed for each gHBfix combination at the reference replica (T = 298 K). See Figure S6 in Supporting Information for examples of the other populated conformations from the REST2 simulations. The number in parentheses displays the χ^2^, which further validates the simulations against the data from experiments. ^a,b^

| (-NH...N-)<br>interaction bias (kcal/mol) | (-NH...O-) interaction bias (kcal/mol) |  |  |
| --- | --- | --- | --- |
|  | (NH...O)<br>+0.0 | (NH...O)<br>+0.5 | (NH...O)<br>+1.0 |
| (NH...N)<br>+1.0 | 46.2/9.7 (0.32<sup>c</sup>) | 35.1/6.0 (0.78) | 13.5/2.4 (1.98) |
| (NH...N)<br>+0.5 | 56.8/5.6 (0.11) | 55.9/9.0 (0.23) | 33.2/5.7 (0.95) |
| (NH...N)<br>+0.0 | 58.7/7.4 (0.15)<sup>d</sup> | 57.6/9.8 (0.15) | 54.1/9.3 (0.42) |

<sup>a</sup> gHBfix $\left(\begin{smallmatrix} 2-\text{OH}\dots\text{nbO}/\text{bO} \\ -0.5 \end{smallmatrix}\right)$ weakening Sph interactions is applied in all simulations.
<sup>b</sup> Cluster analysis performed for the last 7 out of 10 μs-long trajectory at 298 K.
<sup>c</sup> $\chi^2$ values were obtained by calculating and comparing backbone $^3J$ scalar couplings, sugar $^3J$ scalar couplings, NOEs and uNOEs (see Table 3 for details).
<sup>d</sup> REST2 simulation with the $\text{gHBfix}^{(2-\text{OH}\dots\text{nbO}/\text{bO})}_{-0.5}$ potential only (Table 3, last column).

**Figure 4.**
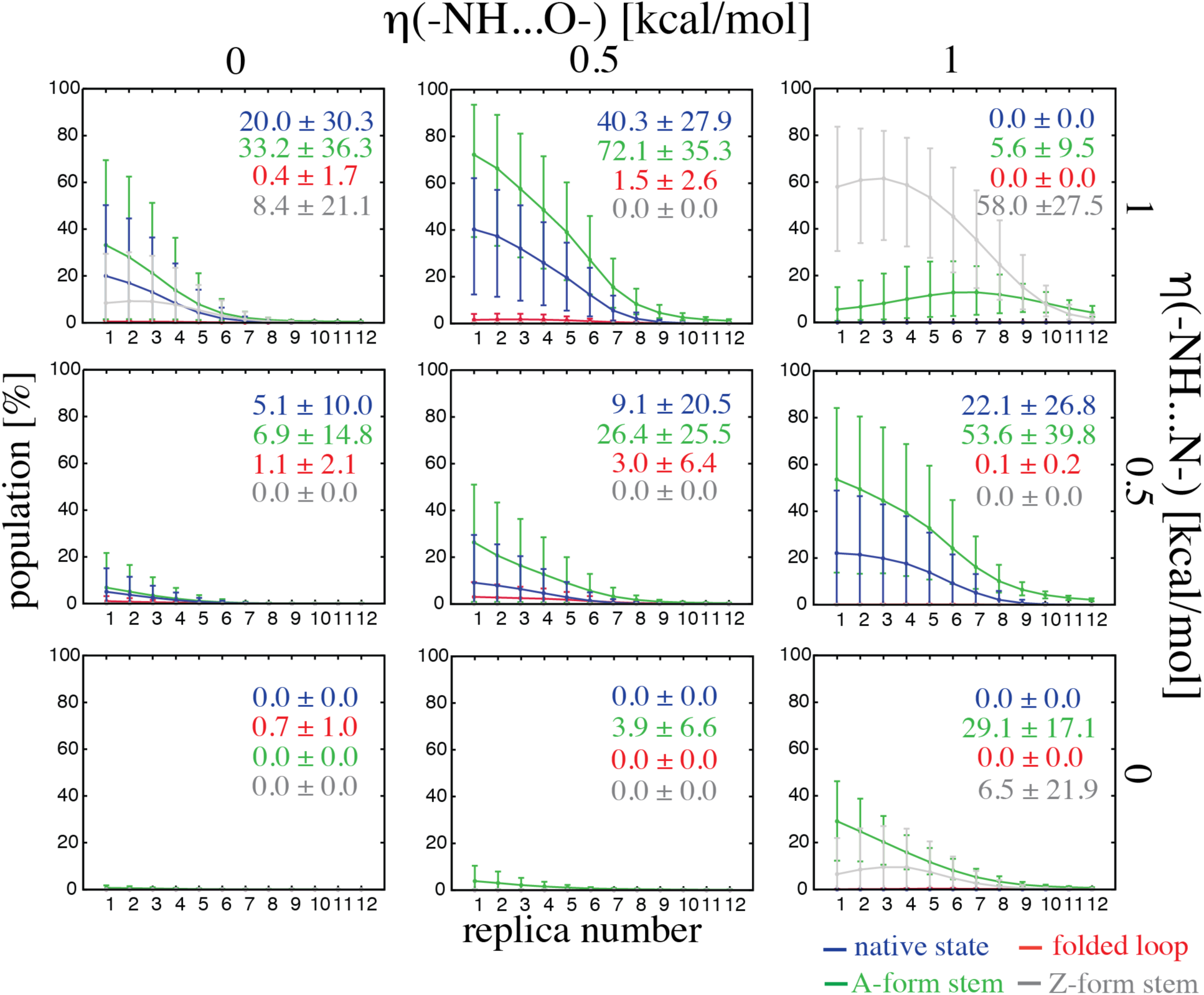
Populations (%) of the most important types of structures in r(gcGAGAgc) REST2 simulations with various gHBfix potentials for all twelve ladder replicas with errors estimated using bootstrapping with resampling both over time- and replica-domains (see Methods for details). Population of the native state is shown in blue. The remaining populations represent correctly folded A-form stem with any loop conformation (in green), correctly folded apical loop with stem not folded (red), and structures with left-handed Z-form stem with any loop conformation, i.e., including structures with properly structured apical loop accompanied with Z-form stem (gray). Displayed numbers highlight the final population in the unbiased replica 1 (T=298 K). The 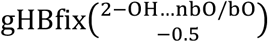 potential was applied in all simulations.

**Figure 5.**
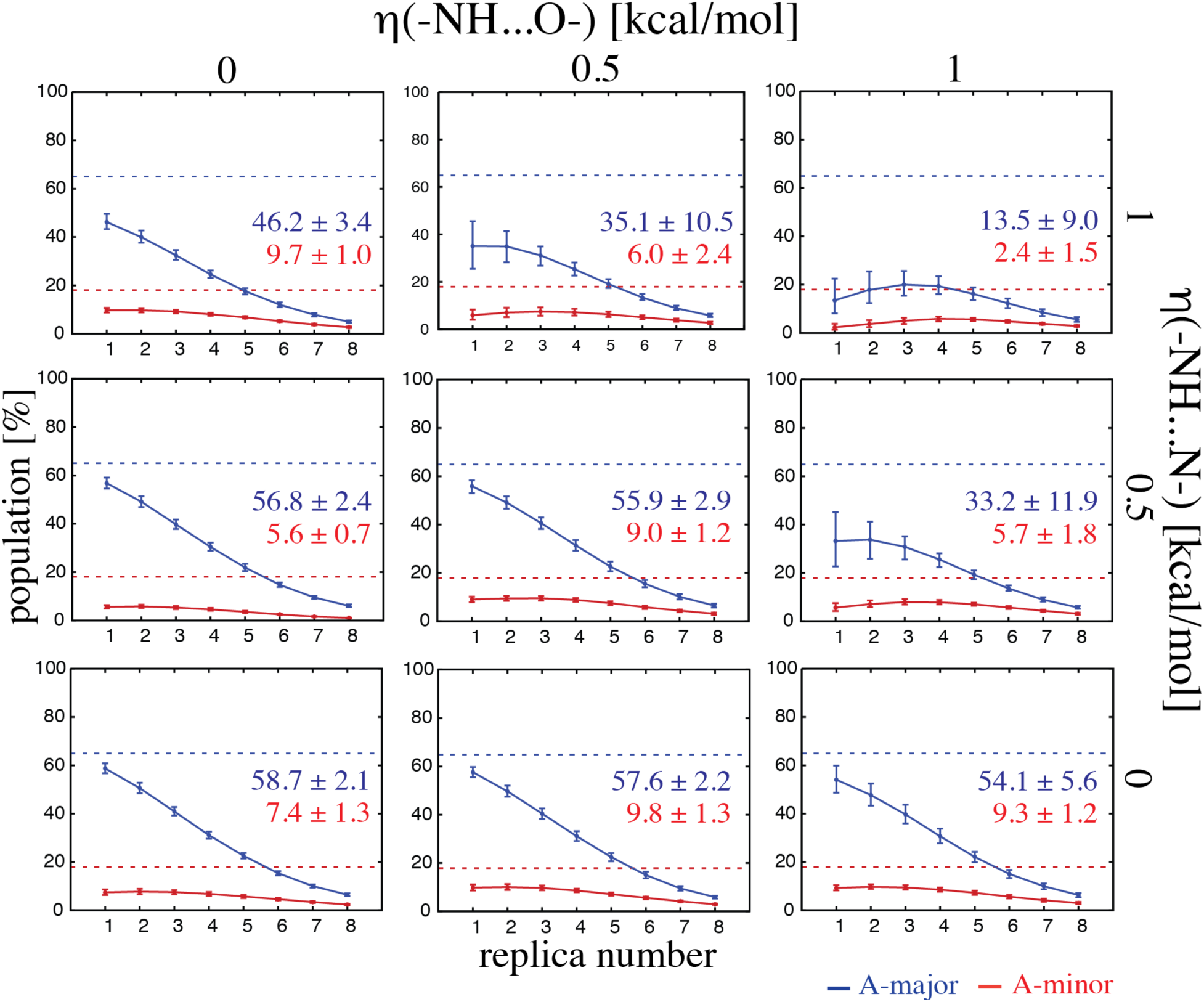
Population analysis of r(GACC) REST2 simulation with various gHBfix potentials. Occurrences (in %) of major conformers, i.e., RNA A-major (in blue) and A-minor (in red) is shown for each eight replicas in the ladder. Displayed numbers indicate the final population in the unbiased replica 1 (T=298 K). Dashed horizontal lines indicate populations suggested by experiments.^25^ Note that the 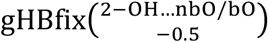 potential was applied in all simulations.

### Present combination of gHBfix parameters is not sufficient to entirely eliminate the intercalated state in other TN sequences

Besides the r(GACC) TN sequence, four other TNs, i.e., r(CAAU), r(CCCC), r(AAAA), and r(UUUU), are commonly used for *ff* validation^14-15, 17, 26-27, 29, 31, 33^ due to availability of the corresponding benchmark NMR data. We thus performed additional REST2 simulations of those TN sequences in order to explore the effects of the 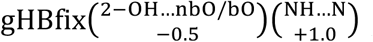 potential. The simulations were initiated with the same setup as for the r(GACC) TN (see Methods). Application of the gHBfix *ff* to these sequences resulted in a visible but, in contrast to r(GACC), not fully satisfactory improvement over the χ_OL3CP_. Namely, the population of native A-form was increased in all four sequences while the population of artificial intercalated structure was significantly reduced only in two of them, r(CCCC) and r(AAAA) (see Table 6). Population of intercalated structures still remained unsatisfactory for r(CAAU), r(CCCC), and r(AAAA) sequences, and notably, the r(CAAU) sequence revealed intercalated structure to be still more populated than the native A-form (Table 6). This suggests that the 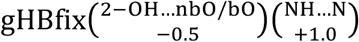 variant is not yet robust enough to entirely eliminate all artificial intercalated states in TNs and further *ff* modifications would be vital. It is worth to note that with the 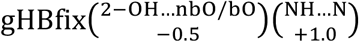 correction, all sequences having still unsatisfactory population of the intercalated structures involve C or A as 5’-terminal nucleotide, i.e., nucleotide, which can form 7BPh interaction^98^ in the intercalated state. This indicates that weakening of BPh interactions that were previously reported to be overpopulated in unfolded states^62^ might be used to eliminate the artificial intercalated structure. Work is in progress in our laboratories to find appropriate gHBfix potentials and parameters to refine simulations of all TNs simultaneously. However, although we already have promising results, finding the best solution is beyond the scope of the present study. It requires testing of a large number of parameter combinations, including subsequent verification of absence of side effects for a broad set of other RNA systems.

**Table 6.** Population (%) of native RNA A-form (including both A-major and A-minor states) and artificial intercalated structures as obtained in standard χ_OL3CP_ and with the suggested 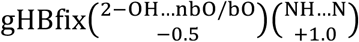 modification.

| sequence | A-form |  | Intercalated |  |
| --- | --- | --- | --- | --- |
| | $\chi_{OL3CP}$ | gHBfix | $\chi_{OL3CP}$ | gHBfix |
| r(CAAU) | <~2%<sup>a</sup> | ~21%<sup>d</sup> | ~50%<sup>a,b</sup> | ~51%<sup>d</sup> |
| r(CCCC) | ~20%<sup>b,c</sup> | ~54%<sup>d</sup> | ~56%<sup>b,c</sup> | ~39%<sup>d</sup> |
| r(AAAA) | ~21%<sup>a,b</sup> | ~25%<sup>d</sup> | ~23%<sup>a,b</sup> | ~11%<sup>d</sup> |
| r(UUUU) | ~4%<sup>a,b</sup> | ~5%<sup>d</sup> | ~18%<sup>b</sup> | ~3%<sup>d</sup> |

<sup>a</sup> Ref.<sup>27</sup>
<sup>b</sup> Ref.<sup>33</sup>
<sup>c</sup> Ref.<sup>14</sup>
<sup>d</sup> this study. Note that we used the implementation of modified vdW parameters of phosphates by Case et al.<sup>63</sup> including correction for the affected dihedrals (see Methods).

### Comment on BPh interactions

When tuning H-bond interactions we need to take into account that under or over-estimation of a given H-bond type is likely not uniform across diverse RNA structures. The apparent *ff* deficiency for a given interaction reflected by its unrealistic population can be substantially context-dependent, as it is also affected by contributions from many other terms such as base stacking, backbone conformations, etc. Thus, one has to be always concerned with potential of over-corrections, which could improve some structures but deteriorate others. This is the reason why we so far did not apply gHBfix for the BPh interactions. Tuning of BPh interactions may have conflicting consequences in simulations of TNs, where they should be avoided and TLs, where they do form significant signature interactions. As BPh interactions are widespread in folded RNAs,^98^ their uniform weakening or strengthening in the *ff* could lead to ambiguous results. We nevertheless work on finding suitable gHBfix modifications of the BPh interactions that would improve RNA simulations without introducing side effects.

### UNCG TL as a challenging system

As noted in the Introduction, earlier studies indicated that the UNCG TL is a considerably more difficult system than the TNs and GNRA TLs. In order to prove that the native state^99^ population of the r(gcUUCGgc) TL might be improved by modification of non-bonded terms, we initially applied the structure-specific HBfix potential to all ten native interactions^61, 99^ of the r(gcUUCGgc) TL motif, i.e., to the six H-bonds of the two canonical GC base pairs of the stem and the four signature H-bonds within the loop (Figure 1). Each native H-bond was biased by +1.0 kcal/mol in favor of the bound state (see Methods for details). Similar structure-based HBfix successfully folds the r(gcGAGAgc) TL in T-REMD simulations.^62^ We obtained converged results, with the population of r(gcUUCGgc) TL native state (27.1 ± 9.6% at 298 K in T-REMD) in a reasonable agreement with experiments^50–52^ (see Supporting Information for details). With aid of the structure-based HBfix, the r(gcUUCGgc) TL was readily able to significantly populate the native fold including *syn*-orientation of the G_L4_ nucleotide (Figure 1), without any specific tuning of the guanosine dihedral potential. However, we note that the HBfix corrections are structure-specific and cannot thus be used for general RNA sequences.

Next, we tried gHBfix potentials that revealed promising results for structural description and folding of both r(GACC) TN and r(gcGAGAgc) TL. Namely, we performed r(gcUUCGgc) REST2 folding simulation with 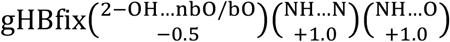 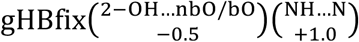 potentials (Table S1 in Supporting Information). Unfortunately, we did not detect any presence of the native state. The application of 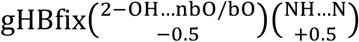 showed rapid stem folding but the loop fluctuated between several non-native conformers (Figure S8 in Supporting Information). In contrast, the 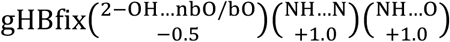 and 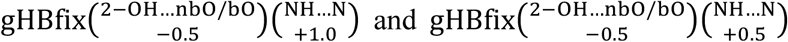 potentials led to either zero or marginal stem folding while misfolded states dominated the simulations (Figure S8 in Supporting Information). The fact that the 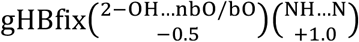 potential did not lead to any folding of the native state in r(gcUUCGgc) TL, while it was sufficient to reveal significant folding in r(gcGAGAgc) TL confirms that the inherent stability of the UNCG loop region is underestimated by the *ff* significantly more than in the case of GNRA TL.^53^ This conclusion is further supported by the observation that only misfolded loop states are populated when the stem is forced to fold by 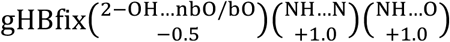 potential, which already overstabilizes the base-base interactions (see above).

The analysis of misfolded conformers obtained by the r(gcUUCGgc) REST2 simulations with the presently available gHBfix potentials showed several noncanonical interactions in the loop region dominating over the native ones. Based on this analysis, we decided to probe effect of either stabilization or destabilization of few other kinds of H-bonds. In particular, we tested the effect of the stabilization of all possible interactions between 2’-OH groups and H-bond acceptors of nucleobases, and the destabilization of interactions between all nucleobase proton donors (–NH) and all O2’ oxygens, and between 2’-OH groups and O2’/O4’ oxygens. We observed that the population of structures with correctly folded A-form stem slightly increased (up to ~70% in some cases, see Supporting Information for more details) but the folded native state appeared only in simulation using the most complex 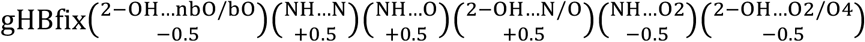 potential, with a rather marginal population of ~6 %. It was accompanied with ~14% population of misfolded states involving properly folded A-form stem but non-native loop conformations (Figure S8 and other details in Supporting Information). In summary, additional biases to SPh and base-base interactions were not sufficient to correct the free-energy imbalance between native folded and misfolded states of the r(gcUUCGgc) TL in χ_OL3CP_. Despite some possible convergence uncertainties of REST2 TL folding simulations, the data suggests that correct description of the r(gcUUCGgc) folding might be precluded also by some other *ff* inaccuracies whose elimination goes beyond fine-tuning of the non-bonded terms. In contrast to TNs which should be relatively easily correctable (see above), we presently have no clues how to decisively improve folding of the UNCG TLs.

### Application of the suggested set of gHBfix parameters to a diverse set of folded RNAs shows no side effects

Structural dynamics and folding simulations of both TN and TL motifs revealed that the combination of the 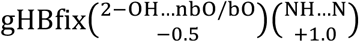 potential with χ_OL3CP_ significantly improves their structural behavior. Although further work is required, especially considering folding of the r(gcUUCGgc) TL, we decided to test the performance of the χ_OL3CP_/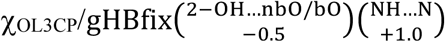 combination for other important (and much larger) RNA motifs and systems. Namely, we performed a set of standard MD simulations of Sarcin-Ricin loop, ribosomal L1-stalk, Kissing-loop complex, Hairpin ribozyme, preQ and Neomycin-sensing riboswitches, Kink-turns, RNA duplex and G-quadruplex (see Table S2 for overview of unbiased MD simulations). Importantly, the gHBfix potential with the presently suggested parameters did not cause any side effects and unexpected rearrangements in comparison to simulations with the unmodified widely used χ_OL3CP_ RNA *ff* (see Supporting Information for details). Therefore, the suggested set of gHBfix parameters appears to significantly improve simulation performance for at least some difficult RNA systems while not causing any undesired side effects in standard simulations of folded RNAs.

### The relationship between gHBfix and NBfix approaches

As discussed above, the gHBfix term is designed for versatile and fully controllable fine-tuning of specific pairwise interactions. It should be noted that at least part of the effect of gHBfix might be also achieved by the standard *ff* terms, in particular by modification of the pairwise vdW parameters via breakage of the combination (mixing) rules, i.e., by usage of the so-called NBfix term; for a recent review see Ref. ^100^. The NBfix approach has been in the past rather commonly applied in connection with proteins,^100–105^ carbohydrates,^106^ DNA molecular assemblies^57, 100^ and for cation interactions.^107–108^ Occasional attempts were made also for RNA molecules.^31, 54^

Nonetheless, the NBfix term has several drawbacks and limitations both from the perspective of *ff* development as well as *ff* performance in practical applications. Further, the earlier attempts to use NBfix term for RNA simulations used an implementation, where the solvent-solute interactions were retained after modifications of the solute atomic vdW parameters.^31, 54^ In this approach all the intra-RNA interactions involving the modified atoms are scaled. We suggest that such NBfix approaches are not convenient for nucleic acids because they are seriously prone to undesired side effects. For instance, below we show that the recently introduced vdW/NBfix modification by Pak *et al*.^31^ designed to improve the behavior of TNs causes swift disruption of the native conformation of the K-turn RNA motif (see the next section called Performance of other *ff*s for details) and is thus not suitable for common simulations of folded RNAs. This underlines that any significant *ff* modifications have to be extensively tested prior publication on a broad range of RNA systems to detect uncontrollable side effects. In contrast to the above-mentioned implementations of NBfix to RNA molecules, we would recommend applying the NBfix terms rather on specific pairwise interactions in a similar manner as suggested here for the gHBfix approach. Such pairwise-specific NBfix parametrizations have been used for proteins and ions in the past.^100, 104-105^

However, even performance of the pairwise-specific implementation of NBfix term may be sometimes limited as the modification of vdW parameters affects the interaction energy indirectly via electrostatics. Thus, for example, smaller vdW radii allow closer contact of the atom-centered partial charges, which in turn affects the interaction energy. In other words, although the NBfix approach formally modifies the vdW parameters, its primary effect comes from the electrostatic term and is dominantly localized in the region of short interatomic distances (at the interatomic contact region). As a consequence, the effect of the NBfix term might be ambiguous and substantially dependent on partial charges of the interacting atoms. For instance, although U(N3-H3)…A(N1) and G(N1-H1)…C(N3) interactions would share the same NBfix parameters they would be affected unequally due to the different partial charges (see Figure S9 and other details in the Supporting Information). In addition, only limited range of energy (de)stabilization might be achieved by the NBfix term in order to avoid extensive geometrical distortions of the molecular contacts, e.g., to avoid potentially excessively clashed or elongated geometries caused by modification of pairwise vdW radius in the NBfix term. Finally, some polar hydrogen atoms, like those from the 2’-OH groups, have zero vdW radii in the parent *ff*, which causes obstacles for fitting extra stabilization typically done by decreasing vdW radii. The way to proceed perhaps would be to combine very small hydrogen-atom radii with very high value of the epsilon parameter (depth of the Lennard-Jones potential). Note that when using the gHBfix modification we ultimately found that it is very useful to implement it via the polar hydrogens. In contrast to NBfix, the gHBfix modification straightforwardly tunes the stability of a given H-bond interaction and its effect is distributed over a broader range of interatomic distances beyond the interatomic contact region (Figure S10 in the Supporting Information). The suggested gHBfix term modifies the pairwise potential at intermolecular distances, where typically polarization is assumed to contribute, which is also one of the reasons, which to our opinion justifies the use of the gHBfix term. In our work we have used a default 1 Å interval for application of the gHBfix bias, however, this parameter (as well as the onset of the biased region), could be further tunable, which may provide an additional flexibility for the refinements. In order to illustrate the relation between the gHBfix and NBfix approaches, we aimed to parameterize particular pairwise-specific NBfix terms to match as closely as possible the effect of our suggested gHBfix terms. In other words, we attempted to find NBfix parameters that would be equivalent to our so far best 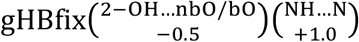 parameters. In order to parameterize the NBfix term, we calculated interaction energies as a function of the separation distance in a small testing system containing only H-bond donor, H-bond acceptor and hydrogen atom. We adjusted the NBfix parameters to reproduce the interaction energy curves obtained with the gHBfix term (see Supporting Information for details, including the NBfix parameters). Thus, we aimed to find (i) NBfix parameters that would provide ~+1.0 kcal/mol stabilization of all possible –NH…N– H-bonds (for stabilization of base-base interactions) and, simultaneously, (ii)another NBfix term that would provide ~–0.5 kcal/mol destabilization of all 2’-OH…bO/nbO H-bonds (for destabilization of SPh interactions, Table S4 in the Supporting Information). Note that the effect of NBfix might differ due to different partial charges of interacting atoms as mentioned in the previous paragraph. Such uncertainties of the NBfix correction derived from gHBfix became apparent, when we validated the fitted NBfix terms for base-base interactions by calculating the interaction energies for simple interacting fragments, i.e., AT and GC Watson-Crick base pairs (see Figure S10 in the Supporting Information), as a function of the base-pair stretch. As expected, the gHBfix approach was able to reproduce exactly the requested energy stabilization of 1.0 kcal/mol without affecting the position of stretch minima. However, the NBfix fit revealed non-uniform effect on the stabilization, as the observed change in the interaction energy in one-dimensional stretch scans was 1.2 kcal/mol and 0.6 kcal/mol for AT and GC base pairs, respectively (see Supporting Information for details). Note that although these optimized NBfix parameters lead to the stabilization effects distributed around the target value of ~1 kcal/mol, the distribution is rather broad and cannot be narrowed by further tuning of the NH…N NBfix parameters due to reasons described above. Subsequently, we tested our best attempt of the NBfix correction derived from the gHBfix for both base-base and SPh interactions (Table S4 in the Supporting Information) in a REST2 folding simulation of GACC TN. The results were rather comparable with the gHBfix folding simulations; the population of A-major, A-minor, and Intercalated states was 37.7%, 16.2%, and 4.6%, respectively, albeit the χ^2^ value with respect to NMR observables was increased to 0.72.

We would like to reiterate that, for reasons explained above, introduction of NBfix terms is equally unphysical as that of gHBfix terms. The gHBfix implementation is more straightforward for post-processing trajectory analysis (e.g., reweighting). In contrary, NBfix is, probably, easier to handle for users in current implementation of the MM packages. Most importantly, a controllable *ff* modification by the NBfix is much more difficult to parameterize than the gHBfix (see above). Taken together, we suggest that the gHBfix term may be more flexible for fine-tuning of the non-bonded interactions, avoids some side-effects and provides some advantages with respect to the NBfix approach in pair-specific treatment of H-bonds and interatomic contacts. Obviously, further work will be necessary to fully characterize the similarities and differences between the gHBfix and NBfix approaches. However, this issue is already beyond the scope of the present work and will be in detail addressed in our future studies. It is also possible that both approaches can be combined in future.

### Comment on other RNA *ff*s

As noted in the Introduction, despite occasional optimistic claims in the literature, none of the other *ff*s available in the literature as of the end of 2017 was proven to satisfactorily simulate the RNA TLs and TNs without undesired side effects, as reviewed in Ref. ^4^. Another RNA *ff* modification (abbreviated as DESRES here) has been published recently,^60^ reparametrizing the non-bonded as well as dihedral terms of the AMBER RNA *ff* and complementing the resulting parametrization with a specific water model.^109^ We have tested this *ff* (see Supporting Information for full details) with the following results.

The DESRES parameters, as implemented by us (see Supporting Information for the parameters), lead to serious side effects for some important folded RNA molecules. For example, structures of RNA Kink-turns and ribosomal L1-stalk RNA segment were entirely and reproducibly disrupted (Figure 6 and Figures S19 and S20 in Supporting Information), suggesting that the DESRES *ff* may be excessively biased in favor of the A-form RNA while also some noncanonical base pairs may be destabilized by the introduced modifications of the non-bonded terms (Figure 6 and Figures S15-S18 in Supporting Information). Similarly to the DESRES potential, the recent vdW/NBfix tuning by Pak and coworkers^31^ also revealed poor description of Kink-turn noncanonical interactions during classical MD simulations (Figure S20 and Supporting Information for details; further testing of this *ff* on additional systems was not performed). Such unstable behavior of two recently published *ff*s has never been observed in common AMBER *ff*s (including the version used and tested here modified by the gHBfix potential), which work well for these systems. Note that the tested Kink-turn 7 is type of Kink-turn that folds in absence of proteins. In presence of monovalent ions, experiments show that the Kink-turn 7 folds at around 30 mM Na^+^,^110–111^ which is well below the monovalent ion concentration used in our simulations. Thus, with a correct *ff*, Kink-turn 7 should be dominantly folded at our simulation conditions. Swift loss of the Kink-turn 7 fold on a sub-μs time scale is thus indication of some major *ff* imbalances.

**Figure 6.**
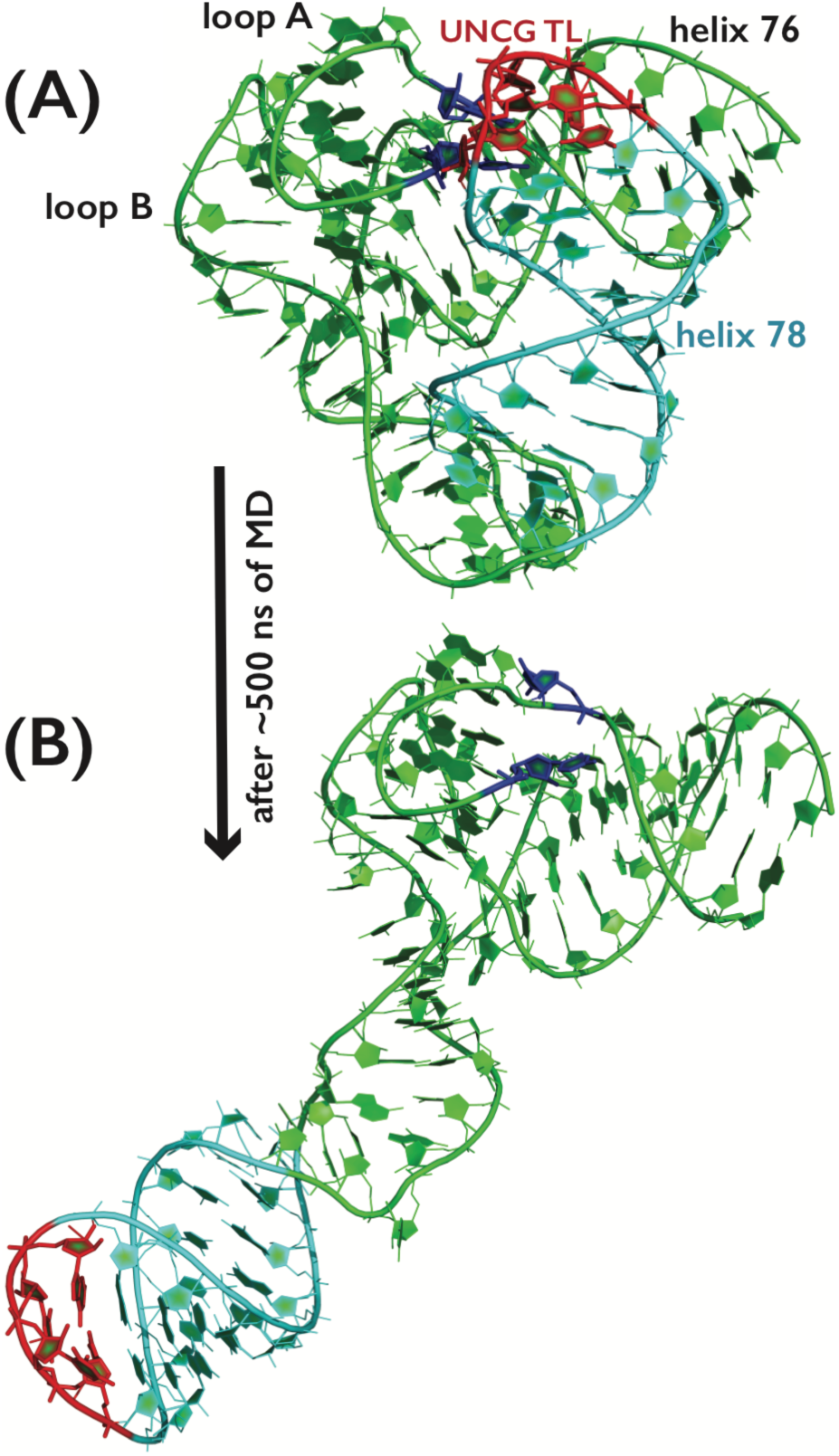
The example of poor description of the folded RNA molecule containing several noncanonical interactions during classical MD simulations with the recent DESRES^60^ RNA *ff*. (A) The starting snapshot from the MD simulation of the ribosomal L1-stalk RNA segment, where the compact fold is characterized by presence of two Kink-turns and by numerous tertiary interactions between three RNA loops, including the UNCG TL (highlighted in red; bases forming base-pair triplets with UNCG TL nucleotides are highlighted in blue). Water molecules, counter ions and hydrogen atoms are not shown for clarity. (B) Last snapshot from the simulation of the same system, where loss of several tertiary interactions, coupled with poor description of Kink-turns, was followed by large-scale degradation of the RNA fold. The degradation began immediately and fully progressed till the end of MD. Basically the same behavior was observed in a number of independent simulations (see Supporting Information for details).

Further, using extensive REST2 simulations with the DESRES potential, we were not capable to fold the r(gcGAGAgc) and r(gcUUCGgc) TLs, despite the fact that even these short constructs should have a non-negligible folded population based on experimental data.^52^ In other words, the DESRES potential^60^ with our protocol/implementation did not bring any observable benefit over standard χOL3CP^26-27, 50-52^ for the description of small 8-mer TLs. Note that in the original paper, the DESRES *ff* was reported to fold TLs with very long stems. Thus, the folding events may result from the increased propensity to form A-form double helix. Therefore, as another test, we have carried out series of unbiased DESRES simulations of the UNCG TL with a long stem starting from the folded states and we have observed loss of UNCG signature interactions on a time scale around 5-10 µs in all of them (Figure S13 in the Supporting Information). We thus suggest that the DESRES *ff* would require more tests before being used to simulate general RNA motifs.

## CONCLUDING REMARKS

Several recent studies suggested that the nucleic acid *ff*s suffer from imbalanced description of the non-bonded interactions.^3, 14-15, 17, 31, 53, 57, 60, 62^ Here we introduce a general pair potential, a generalized HBfix (gHBfix), that can selectively fine-tune specific non-bonded terms, in particular specific types of hydrogen-bonding interactions. The gHBfix potential can be easily combined with any existing *ff*s to probe the effect of the modification of their non-bonded terms. It is easily applicable for common MM engines such as AMBER and GROMACS packages without any requirement for the source code modifications (see Supporting Information for the attached toolkit, which generates files for MM packages). Most importantly, unlike in case of vdW or point-charge parameter modifications, it affects only the selected type of the interactions, while the others, including, e.g., interactions with the solvent and ions, remain intact. Thus, it reduces the likelihood of introducing undesirable side effects that are frequently associated with other ways of modifying non-bonded terms. The gHBfix term is an entirely legitimate way of modifying the pair-additive NA *ff* since, in fact, it is as empirical as the other terms. The common MM parameters such as atomic charges and radii are not QM observables and thus do not correspond to any real physical properties. So it seems genuine to straightforwardly target interaction energies of selected molecular contacts instead of trying to refine their properties indirectly via modified parameters of the MM atoms. In other words, the gHBfix term is as general as all the other *ff* terms and can be to certain extent physically justified, as it may compensate, e.g., for the lack of polarization terms in H-bonding.^4, 62^ As extensively discussed, a similar effect can be achieved also by modification of vdW parameters via NBfix,^31, 54, 57, 100^ although to our opinion in significantly less controllable manner with higher risk of causing some major simulation artifacts for other systems. It is possible that in future the NBfix and gHBfix corrections could be applied simultaneously, in a synergy.

We tested the gHBfix potential for fine-tuning of the solute-solute H-bond interactions in folding of small TL and TN RNAs; in particular by strengthening of the base-base interactions and weakening of the SPh interactions. Incorrect description of both interaction types was suggested to, e.g., interfere with balanced description of the TLs folding. We found out that the negative gHBfix applied to all possible interactions between 2’-OH groups as proton donors and nbO and bO oxygens as acceptors (i.e., weakening of the SPh interactions) eliminates (or reduces) some *ff* artifacts, namely, the presence of intercalated structures of TNs. Furthermore, the additional support of (–NH…N–) H-bonds (i.e., gHBfix strengthening the base pairing interactions) significantly promotes population of the TL folded stem and of the native arrangement of the GNRA TL. The necessity of additional support of H-bonds between groups with N atoms is probably related to their oversized vdW radii in *ff*s as suggested in some recent studies.^50-52, 96^ In summary, we propose a 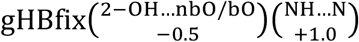 potential as an extension of the AMBER χ_OL3CP_ *ff*. The conformational ensembles generated by this *ff* reveal better agreement with available experimental data than the corresponding ensembles generated by the χ_OL3CP_ alone. In particular a significant improvement was observed in population of both canonical A-major and A-minor single stranded structures in r(GACC) TN folding simulations, together with elimination of the intercalated structures. It also achieved a significant population of native state in the simulation of small 8-mer GNRA TL. Most importantly, we have tested the suggested gHBfix tuning on a wide portfolio of other RNA structures without observing any undesired side effects in standard simulations. We suggest that especially the improved stabilization of base pairs may be profitable for many other systems in long simulations. In particular, the gHBfix potential also reduces the A-RNA end fraying (see Supporting Information for details), which was shown in the past to cause problems for example in simulations of some protein-RNA complexes.^112^ Although not directly tested, the gHBfix *ff* should be readily applicable to protein-RNA complexes, as the modification does not have any direct impact on the interactions at the protein-RNA interface.

Despite that the suggested specific gHBfix parameters provide undisputable improvement of the standard AMBER RNA *ff*, the modification is not yet sufficient to fully eliminate intercalated structures for r(CAAU), r(CCCC), r(AAAA) TNs and to correct the free-energy imbalance between folded and misfolded states observed for the challenging r(gcUUCGgc) TL. Thus, further tuning of the gHBfix potential is required and it seems that UNCG TL might require additional corrections that go beyond simple fine-tuning of H-bond interactions. On the other hand, correction of the remaining TNs should be rather straightforward thought, as discussed above, one has to be careful to avoid overstabilization of the A-form RNA, a major problem of several recently introduced *ff*s. Considering the variability and complexity of RNA structures, and the simplicity of the functional form of current *ff*s, it is probably naive to suppose that one would be able to introduce a perfect RNA *ff* with the ability to provide correct behavior of all RNA systems by tuning just some non-bonded terms within the framework of pair-additive *ff*s.^4^ Nevertheless, the introduced gHBfix potential increases the flexibility of the *ff* in a controlled way by adding small number of independent parameters directly modifying stabilities (populations) of the target interactions, indicating that performance of the pair-additive *ff*s could still be significantly improved. In general, *ff* improvements may be aided by using new terms that go beyond the basic conventional *ff* form and that further increase the flexibility of the *ff*. gHBfix is an example of such modifications.

The computational overhead of the gHBfix is rather insignificant for small systems (~2% for GPU accelerated simulations using fast GTX1080Ti GPU cards and even smaller for slower GPU cards or CPU version of the MM engine code). Although the computational overhead increases with the system size, it might be almost entirely eliminated by the direct implementation of gHBfix potential into the MM engine code in future, which would allow using the Non-Bonded pair list for calculations of the gHBfix terms. The specific version of RNA AMBER *ff* presented and tested in this work does provide some improvements while not showing any undesired side effect for the broad set of important folded RNA systems. It can thus be quite safely used in RNA simulations. Work is in progress to balance some additional H-bond interactions using the gHBfix potentials, to optimize the biasing parameters Ƞ via reweighting and to merge the gHBfix potentials with adjustments of some of the core *ff* terms. We suggest that future refinements of the pair-additive RNA *ff*s will likely require adding additional *ff* terms (such as the gHBfix) to the basic functional form to increase flexibility of the parametrization. This, together with testing on a broad set of RNA systems, will help to avoid over-fitting of the *ff* in favor of one type of target RNA structures, such as, e.g., the TNs or A-form RNA, a problem demonstrated in our study for several recently published *ff*s. Due to the huge dimensionality of the *ff* parameter space and their mutual inter-dependence, finding an optimal *ff* will be most likely a long-term process, which would profit from collaborative efforts within the RNA simulation community.

## ASSOCIATED CONTENT

### Supporting Information

The following files are available free of charge. Details about convergence of enhanced-sampling simulations, description of the r(gcUUCGgc) T-REMD folding simulation with structure specific HBfix, detailed description of additional REST2 folding simulations with various gHBfix potentials, further details about the relation between the gHBfix and NBfix approaches, structure preparation and other details about simulation protocols, detailed description of REST2 and standard MD simulations with recently published *ff*s, description of standard MD simulations with the gHBfix potential, Supporting Tables and Figures (PDF). C++ code to generate inputs for AMBER and GROMACS in order to perform simulations with the gHBfix (gHBfix.pdf). AMBER input files containing DESRES parameters (desres.zip).

### AUTHOR INFORMATION

Corresponding Authors

## ACKNOWLEDGMENT

We thank Petr Jurečka and Petr Stadlbauer for useful discussions. Sandro Bottaro is acknowledged for critically reading the manuscript and providing comments. Financial support from the Operational Programme Research, Development and Education – European Regional Development Fund project no. CZ.02.1.01/0.0/0.0/16_019/0000754 [to M.O., P.B., P.K.] of the Ministry of Education, Youth and Sports of the Czech Republic and Czech Science Foundation [18-25349S to P.B., P.K.] are acknowledged. J.S. acknowledges support by Praemium Academiae.

