## Supporting Information for "Improving The Performance Of The Amber Rna Force Field By Tuning The Hydrogen-Bonding Interactions"

**Additional comment on convergence of replica exchange simulations.**

The necessity to obtain converged enhanced sampling simulations with minimal statistical errors limits the evaluation of force fields (*ff*s) on larger systems. One can rigorously compare simulations with different *ff* variants only when all simulations are sufficiently converged, otherwise it is not possible to confidently separate the *ff* differences from sampling limitations.[1] The convergence of temperature replica-exchange molecular dynamics (T-REMD)[2] and replica exchange with solute tempering (REST2)[3] simulations was analyzed by rather complex bootstrapping protocol (see Methods in the main text and Figure S1). The fully converged T-REMD and REST2 simulation should generate equivalent sampling in all continuous replicas.[1, 4] We here used 10 μs-long runs per replica during simulations of both TN and TL RNA motifs. The analysis of r(GACC) REST2 simulations showed that TN motif requires few μs to provide uniform distribution of major conformations within the structural ensemble (Figures S3 and S4). Thus, we took the last 7 μs of each replica for the population analysis of r(GACC) REST2 simulations. As expected, the conformational space is much more complex for the folding simulations of TLs. Note that our TL REST2 folding simulations always started from the unfolded state. In these simulations the first folding events typically appeared at ~3 μs, though in some simulations they occurred in even later stages. As a compromise, we took the last 7 μs of each replica for the population analysis of major conformations in r(gcGAGAgc) and r(gcUUCGgc) REST2 folding simulations (Figure S5).

**Methodology of UNCG T-REMD simulation with structure-specific HBfix.**

The r(gcUUCGgc) folding T-REMD simulation was performed using the AMBER16 suite of programs[5] with *ff*99bsc0$\chi_{OL3}$[6-9] with the van der Waals (vdW) modification of phosphate oxygen developed by Case et al.[10] and OPC[11] water model. The starting single strand structures were generated using unbiased 640-ns-long MD simulations at 300 K starting from the A-form single strand structure generated by nucleic acid builder tool in AMBER. The 64 starting structures of r(gcUUCGgc) TL were taken in 10 ns long intervals. The protocol is in detail described elsewhere.[12] The temperatures of the 64 replicas spanned the range of 278-463 K and were chosen to maintain an exchange rate of ~25%. The T-REMD simulation was performed at constant volume (using the NVT ensemble in each replica), with long-range electrostatics calculated using PME with 1 Å grid spacing and 10 Å real-space cutoff. Langevin dynamics with a friction coefficient of 2 $ps^{-1}$ was used as a thermostat in all replicas, and the exchanges were attempted every 10 ps. Each replica was simulated for 10 μs and the total simulation time of the T-REMD simulation was thus 640 μs (Figures S11 and S12). Note that for this folding simulation we have used the more expensive T-REMD protocol rather than the REST2, to be entirely consistent with the preceding work.[12]

**Details about additional REST2 TN and TL folding simulations with various gHBfix potentials, complementing the corresponding parts of the main text.**

In order to analyze the effect of the positive base-base gHBfix biases for the TN motif, we performed a set of nine REST2 simulations of r(GACC) as in the case of r(gcGAGAgc) TL folding differing in base-base gHBfix terms (Table 5 and Figure 5 in the main text). The $\text{gHBfix}\binom{\text{2-OH}\ldots\text{nbO/bO}}{-0.5}$ weakening SPh interactions was included in all nine simulations. We identified some spurious side effects of certain combinations of base-base gHBfix, namely, in the case of a high +1.0 kcal/mol value of gHBfix bias constant η applied to (–NH…O–) interactions. In such cases, the population of the native RNA A-major conformation was significantly reduced (Figure 5 in the main text). For example,

$\text{gHBfix}\binom{\text{2–OH…nbO/bO}}{-0.5}\binom{\text{NH…N}}{+1.0}\binom{\text{NH…O}}{+1.0}$ potential reduced the population of RNA A-major conformation to just 15.5% which is even below what we obtained during the standard simulation without any gHBfix potential (Table 3 in the main text). Although we did not detect intercalated structures during any r(GACC) REST2 simulation with gHBfix potentials, an excessive support applied to (–NH…O–) interactions resulted in loop-like structures closed by spurious base-base interactions (Figure S6).

Various attempts and different gHBfix biases were designed in order to stabilize the native state of UNCG TL during the r(gcUUCGgc) REST2 folding simulations. Initially, we tested (besides destabilization of the SPh interactions and stabilization of base pairing) the effect of the (i) stabilization of all possible interactions between 2'-OH groups and H-bond acceptors of nucleobases with bias energy η equaling to either +0.5 kcal/mol or +1.0 kcal/mol (simulations denoted as $\text{gHBfix}\binom{\text{2–OH…nbO/bO}}{-0.5}\binom{\text{NH…N}}{+1.0}\binom{\text{NH…O}}{+1.0}\binom{\text{2–OH…N/O}}{\eta}$), and (ii) the destabilization of interactions between all nucleobase proton donors (–NH) and all O2' oxygens, and between 2'-OH groups and O2'/O4' oxygens, both with η equaling to –0.5 kcal/mol (simulation denoted as $\text{gHBfix}\binom{\text{2–OH…nbO/bO}}{-0.5}\binom{\text{NH…N}}{+1.0}\binom{\text{NH…O}}{+1.0}\binom{\text{NH…O2'}}{-0.5}\binom{\text{2–OH…O2'/O4'}}{-0.5}$, see Table S1 in for overview of the REST2 simulations). In both cases, we observed that the population of structures with correctly folded A-form stem slightly increased (up to ~70%) but the loop still sampled only non-native misfolded states similarly to those obtained with the $\text{gHBfix}\binom{\text{2–OH…nbO/bO}}{-0.5}\binom{\text{NH…N}}{+1.0}\binom{\text{NH…O}}{+1.0}$, see Figure S8. Additionally, we probed the simultaneous effect of two previously introduced gHBfix biases, i.e., additional stabilization of sugar-base H-bonds and destabilization of interactions involving O2' and O4' oxygens. In contrast to simulations discussed in the previous paragraph, we also modified gHBfix bias energies η for base-base interactions, so that the additional simulations probed the $\text{gHBfix}\binom{\text{2–OH…nbO/bO}}{-0.5}\binom{\text{NH…N}}{\eta}\binom{\text{NH…O}}{\eta}\binom{\text{2–OH…N/O}}{+0.5}\binom{\text{NH…O2'}}{-0.5}\binom{\text{2–OH…O2'/O4'}}{-0.5}$ function, in which η values equaled either to +0.5 or +1.0 kcal/mol (see Table S1). Despite all attempts, the folded native state appeared only in simulation using the $\text{gHBfix}\binom{\text{2–OH…nbO/bO}}{-0.5}\binom{\text{NH…N}}{+0.5}\binom{\text{NH…O}}{+0.5}\binom{\text{2–OH…N/O}}{+0.5}\binom{\text{NH…O2'}}{-0.5}\binom{\text{2–OH…O2'/O4'}}{-0.5}$ potential, with a rather marginal population of ~6 %. It was accompanied with ~14% population of misfolded states involving properly folded A-form stem but non-native loop conformations (Figure S8).

**Details about the relation between the gHBfix and NBfix approaches.**

We made an attempt to modify interaction curves for specific atom pairs by adjusting Lennard-Jones combining rules (NBfix) in a way to produce similar effects introduced by the $\text{gHBfix}\binom{\text{2–OH…nbO/bO}}{-0.5}\binom{\text{NH…N}}{+1.0}$ potential. We aimed to find NBfix parameters that would provide ~+1.0 kcal/mol stabilization of the –NH…N– base-base interactions, and simultaneously, ~–0.5 kcal/mol destabilization for SPh interactions (Table S4). Considering the base-base interactions, the NBfix effect varies for different atom pairs forming the same kind of –NH…N– interaction because it depends on partial charges of interacting atoms (see Figure S9). Those uncertainties of the NBfix approach are further translated into uncertainties of the total interaction energy bias for different base pairs. To illustrate this issue, we calculated interaction energies of GC and AT Watson-Crick base pairs as a function of the base-pair stretch (one-dimensional scan) and compared interaction energy curves affected by both gHBfix and NBfix approaches. Plots on Figure S10 show that the gHBfix was able to reproduce exactly the requested energy stabilization of 1.0 kcal/mol without affecting the position of stretch minima. In contrary, the NBfix fit revealed ambiguous effect on the stabilization. We identified changes in the interaction energy of 1.2 kcal/mol and 0.6 kcal/mol for AT and GC base pairs, respectively. The depth wells of potentials were shifted as well from those revealed by the standard *ff*99bsc0$\chi_{OL3}$[6, 8-9] AMBER *ff* with the vdW modification of phosphate oxygen developed by Case et al.[10] (Figure S10). Notable is also the fact that the NBfix effect is localized in the narrow range of stretch while the gHBfix approach distributes the stabilization effect more smoothly.

**Folding and conformational sampling studied by the recently suggested RNA *ff*.**

Very recent work by the D.E. Shaw laboratory attempted to improve RNA *ff* by modifying specific vdW parameters, charges, and dihedral parameters (denoted here as DESRES potential[13]). We have adapted the suggested parameters and thus the attached files (desres.zip) contain our implementation of DESRES parameters for AMBER. The DESRES *ff* version is to be used with the TIP4P-D water model (previously developed for simulations of disordered proteins[14]). The original study by Shaw *et al.*[13] provides very extended set of test simulations including TNs, TLs, single-stranded RNA, and RNA duplexes. On the other hand, test simulations for folded RNAs were limited to standard simulations of two riboswitches with just simple overall analysis of their behavior.

Here, we initially tested the performance of DESRES potential on similar systems as reported originally, namely r(GACC) TN (the same system as in the original paper[13]), and the r(gcGAGAgc) and r(gcUUCGgc) 8-mer TL's. Note that the original paper[13] used longer sequences, i.e., 10-mer and 14-mer for GNRA and UNCG TL's, respectively.

Consistently with the original study introducing the DESRES potential,[13] we observed that r(GACC) sampled well the native A-RNA conformations in good agreement with the experimental data (population of A-major and A-minor conformation during the 10 μs-long REST2 simulation was ~62 % and ~14 %, respectively, and deviation from the experimental NMR data $\chi^2$ equals to 0.15) with marginal population of spurious intercalated structures (below 1.0%). We note that populations reported in the original study,[13] i.e., A-major ~73 % and A-minor ~3 %, were obtained from unbiased 90 μs-long MD simulation using Anton.[15]

On the other hand, the DESRES potential in our simulations was not capable to fold 8-mer TLs. It is worth to reiterate that in the present study we are using rather short 8-mer TL sequences due to their relatively low melting temperature (~330 K) but still significant fraction of folded state at 298K,[16] which makes them ideal testing systems allowing to probe the free-energy balance between stabilization of the stem and loop region. In contrast, the DESRES potential was originally tested on longer sequences, which might support the hairpin formation by stabilization of the longer stem, since the DESRES *ff* appears to substantially overstabilize A-form duplex and likely also A-form ssRNA (see below). To be specific, we did not observe any folding events in both our DESRES REST2 simulations of r(gcGAGAgc) and r(gcUUCGgc) TLs starting from the unfolded state. The dominantly populated state remained unfolded A-form single strand structures (population of A-form single strand was almost 100% and ~81 % for r(gcGAGAgc) and r(gcUUCGgc) TL, respectively). To probe the convergence of these results, we performed another REST2 simulation of these 8-mer TLs, in which all replicas were initiated from the folded (native) states. These simulations revealed that all replicas rapidly lost their folded conformations (after ~1 μs and ~200 ns of the r(gcGAGAgc) and r(gcUUCGgc) REST2 simulation, respectively). In subsequent phases the simulations populated only the unfolded states as observed in the above-noted REST2 simulations initiated from the unfolded states. This clearly shows that the steady-state populations of the native states in both r(gcGAGAgc) and r(gcUUCGgc) 8-mer TL simulations are negligible with DESRES potential. Note that similar rapid 'unfolding' behavior was observed also in standard *ff*99bsc0$\chi_{OL3}$ simulations (without any HBfix or gHBfix potentials),[17-18] suggesting that while DESRES potential[13] was able to significantly improve conformational dynamics of TNs, in case of small 8-mer TLs it does not bring any benefit over standard *ff*99bsc0$\chi_{OL3}$ AMBER *ff*. The discrepancy in behavior of DESRES *ff* between longer 10-mer and 14-mer TLs reported in the original paper,[13] and short 8-mer TLs reported here might be explained by further stabilization coming from the longer stem. Namely, the estimated $\Delta G_{300K}$ of folding at 300 K using Turner parameters[19] equals to −0.9 and −3.5 kcal/mol for the r(gcGAGAgc) 8-mer and r(cgcGAGAgcg) 10-mer, respectively (see, e.g., Table 2 in Ref. [12]). Notice that results presented here and those from the original study[13] were obtained using different protocols, i.e., REST2 simulations and simulated tempering protocols.[20-21]

In order to obtain more information about the balance in description of canonical and non-canonical structures in DESRES potential,[13] we performed series of additional standard MD simulations of several different RNAs (see Table S2 for the overview of unbiased MD simulations). Firstly, we obviously performed an additional set of five separate standard MD simulations with 14-mer UNCG TL starting from the NMR structure (PDB ID 2KOC).[22] Surprisingly, all five simulations revealed irreversible loss of signature interactions of the TL, i.e., degradation of the TL architecture, on a few μs-long time scale (Figure S13). The typical behavior was that two signature $U_{L1}$(2'-OH)…$G_{L4}$(O6) and $U_{L2}$(2'-OH)…$G_{L4}$(N7) H-bonds were lost and $G_{L4}$ residue flipped out of the loop. Subsequently, the loop conformation was completely disrupted, which was followed by the loss of H-bonds of the closing GC and neighboring AU base pairs (i.e., the indication of stem unfolding within two simulations out of five). Further, an unbiased MD simulation of the smaller 8-mer UNCG TL revealed spectacular irreversible unfolding (after ~650 ns) towards A-form single strand, which was initiated by the base-pair breathing in the short stem (two canonical GC base pairs, Figure S14). We suggest that the declared capability of the DESRES *ff* to fold RNA TLs should be further investigated by other groups. Our attempt to implement the suggested parameters resulted in an RNA *ff* with unsatisfactory behavior.

In summary, while the DESRES *ff* strongly prefers the straight A-form single-strand structure (~76%, ~100%, and ~81% in the reference replica of r(GACC), r(gcGAGAgc), and r(gcUUCGgc) motifs, respectively) the standard *ff*99bsc0$\chi_{OL3}$ AMBER *ff* shows significant sampling also of other structures such as coil-like arrangements. This indicates a shift of the DESRES parameterization (as implemented by us) towards straight A-form RNA conformation at least for short sequences. Further (see below), the DESRES parametrization has produced major instabilities is standard simulations of several important folded RNAs.

**Structure preparation and other details about standard unbiased simulation protocols.**

The starting topology and coordinates of various RNA systems were prepared from particular experimental structures (Table S2) by using the tLEaP module of AMBER 16 program package.[5] RNA molecules for simulations with DESRES potential[13] were solvated by the TIP4P-D[14] water model. Simulations with *ff*99bsc0$\chi_{OL3}$[6-9] and *ff*99bsc0$\chi_{OL3}$ with the gHBfix potential (gHBfix *ff*) used the vdW modification of phosphate oxygen developed by Case et al.[10] and all the affected dihedrals were adjusted as described elsewhere.[23] For the simplicity, the applied RNA *ff* is abbreviated as $\chi_{OL3CP}$ through the remaining text. RNA molecules for simulations with $\chi_{OL3CP}$ gHBfix *ff* were solvated with the OPC[11] water model. The minimum distance between box walls and solute was 10 Å and all simulations were performed in ∼150 mM KCl salt using the Joung−Cheatham[24] ionic parameters. The position of RNA molecule remained constrained during minimization and optimization of waters and ions. Subsequently, all RNA atoms were frozen and the solvent molecules with counter-ions were allowed to move during a 500-ps long MD run under NpT conditions (p = 1 atm., T = 298.16 K) in order to relax the total density. After this, the RNA molecule was relaxed by several minimization runs, with decreasing force constant applied to the sugar-phosphate backbone atoms. After the relaxation, the system was heated in two steps: the first step involved heating under NVT conditions for 100 ps, whereas the second step involved density equilibration under NpT conditions for and additional 100 ps. The particle mesh Ewald (PME) method for treating electrostatic interactions was used. The standard unbiased MD simulations were performed under periodic boundary conditions in the NpT ensemble at 298.16 K using weak-coupling Berendsen thermostat[25] with coupling time of a 1 ps. The SHAKE algorithm, with a tolerance of $10^{-5}$ Å, was used to fix the positions of all hydrogen atoms, and a 10.0 Å cut-off was applied to non-bonding interactions to allow a 2-fs integration step.

Three nonstandard residues were used during $\chi_{OL3CP}$ *ff*, gHBfix *ff*, and DESRES unbiased MD simulations: N1-protonated Adenine, N3-protonated Cytosine and preQ$_0$ ligand. The parameters and charges were taken from our previous works.[26-27] Nonstandard residues for simulations using the DESRES potential[13] were modified as follows: charges were developed according to the Cornell et al. procedure,[6, 28] i.e., similar to those used for $\chi_{OL3CP}$ and gHBfix *ff* simulations, and the vdW and bonding parameters were set in accordance with the DESRES[13] potential. Note that the charges in DESRES potential were arbitrarily tuned during the process of the *ff* parametrization and it is thus not clear how to extend these charge modifications to the other nonstandard residues.

**Unbiased MD simulations with DESRES potential (our implementation).**

Although folding simulations of short RNAs such as TNs and TLs are of interest, the vast majority of RNA simulation studies are (and will be) done for various folded RNA molecules. Note that folded RNA rather than model systems such as TNs are the primary simulation targets for which the *ffs* should be optimized. Thus, any suggested RNA *ff* needs to be tested for these systems. We have carried out an additional set of unbiased DESRES MD simulations for a wide range of RNA systems (Table S2), essentially overlapping with the tests that we have done for our gHBfix *ff*. Note that the gHBfix *ff* was entirely successful in these tests (see below). In contrast, the DESRES variant fails in description of several biochemically relevant RNA systems that are quite well described by the standard $\chi_{OL3CP}$[6-9] casting considerable doubts about applicability of DESRES in general simulations of RNAs.

Firstly, we simulated the Sarcin-Ricin loop RNA motif (SRL, PDB ID 3DW4[29]), a widespread and highly conserved RNA motif which was originally found in helix 95 of domain VI of the large ribosomal subunit.[30] SRL is a rigid RNA motif containing an amazingly intricate network of non-canonical base pairs, BPh interactions and some other H-bonds, complemented by very complex backbone conformations.[31-32] It is being used as a standard benchmark for *ff* testing.[33] Due to its stiffness, SRL is typically globally stable in MD simulations, but none of the available *ffs* is capable to reproduce all details of this intricate system. The overall fold, the triple base-pair stack (GpU platform), and adjacent GNRA TL remained stable also during 1 μs-long unbiased MD simulation with DESRES potential (Figure S15). The G7-A17 BPh interaction within the GpU platform was stable, i.e., the G7(N2H)…A17(*pro*-R$_P$) H-bond revealed similar fluctuations as in the reference $\chi_{OL3CP}$ simulation. On contrary, the other two A6-C18 and U8-G16 BPh contacts were weakened in DESRES simulation (Figure S15). Previous MD studies showed that both signature A6(N6H)…C18(*pro*-R$_P$) and G16(N2H)…U8(*pro*-R$_P$) interactions fluctuate between direct H-bonds and indirect (water bridged) contacts.[33-34] However, the simulation with DESRES potential described both A6-C18 and U8-G16 BPh interactions predominantly as water mediated, i.e., populating indirect contacts for ~95% of the time, Figure S15). This is to be considered as sign of deterioration of the simulations by the DESRES potential.

We then simulated the Hairpin Ribozyme (HrRz, PDB ID 2OUE[35]), which is the member of small self-cleaving Ribozymes. We prepared the system with the canonical G8 and N1-protonated A38 ($A38H^+$, see section Structure preparation and other details about simulation protocols for parameterization) forms of catalytically important residues that were shown to be required for the compact arrangement of the active site during MD simulations.[27] We observed that the overall fold and the S-turn backbone conformation remained stable on the 1μs-long timescale (Figure S16) within three independent simulations using the DESRES potential. Initially, we identified rapid repuckering (C2'-endo/C3'-endo flip) of the A-1 ribose ring, simultaneous loss of G8(N1H)…A-1(O2') H-bond and formation of bifurcated G8(N1H,N2H)…G+1(*pro*-$R_P$/*pro*-$S_P$) H-bonds (Figure S16), which is consistent with the behavior of this systems during MD simulations using the post-*ff*99 RNA *ffs*.[23, 27] However, we also identified spurious loss of G8(N1H)…G+1(*pro*-$R_P$/*pro*-$S_P$) and G8(N2H)…G+1(*pro*-$R_P$/*pro*-$S_P$) H-bonds in two independent simulations. The G8 subsequently departed from the active site and either remained positioned more than ~7 Å away from the scissile phosphate or returned back before being repelled again (Figure S17 and S18). Such a loss of the catalytically important BPh interaction and subsequent distortion of the active site is not compatible with the simulated protonated state and resembles earlier-published simulations with the N1-deprotonated form of G8 ($G8^-$).[27] Such behavior was never observed with the canonical G8 at the active site using the post-*ff*99 AMBER RNA *ffs*.[23, 27, 36-38] Two out of three simulations also revealed weakening of the other catalytically important $A38H^+$(N1H)…G+1(O5') H-bond (Figure S17). Based on all our experience, we again consider this as a visible deterioration of the simulation behavior by the DESRES potential, as it is a visible corruption of the catalytic center.

Next, we simulated Kink-turn which is a recurrent internal loop RNA motif, widespread for example in ribosome, facilitating a sharp bend between two A-RNA double helices.[39-40] Kink-turns have a well-defined topology characterized by non-canonical *trans* Hoogsteen/Sugar Edge A-G base pairs (non-canonical stem) and by canonical *cis* Watson-Crick/Watson-Crick G-C base pairs (canonical stem). The longer strand contains a (typically three-nucleotide) bulge. There is an A-minor interaction between the stems, and a signature O2'/N1 H-bond between the first nucleotide of the bulge and the adenine from the AG base pair closest to the canonical stem.[40] In other words, a folded kink-turn must contain five signature interactions which define its 3D shape and its consensus sequence. Many kink-turns are involved in protein-assisted RNA folding and they were also suggested to facilitate large-scale dynamics movement of the ribosome.[41] For our simulations, we selected the *Haloarcula marismortui* kink-turn 7 (Kt-7) which is very close to the consensual sequence of the kink-turn motif and can thus serve as a useful reference for the motif's behavior in simulations. In all our DESRES simulations of Kt-7, we observed instability of the non-canonical *trans* Hoogsteen/Sugar Edge AG base pairs within few nanoseconds after the start (Figure S19). The instability of the AG base pairs was reversible on the simulation timescale. Then, in later parts of the simulations, we evidenced an irreversible loss of the A-minor interaction between the two stems. After this, the structure entirely lost its characteristic bent shape and completely straightened (Figure S19). This corresponds to a complete loss of the Kt-7 structure in all attempted DESRES simulations, something that has never occurred with any common AMBER RNA *ff*s. In addition, the time-scale of the complete disruption of the motif is very short, ~200 ns, suggesting a significant destabilization of the native fold by the used *ff*. Except of some details, standard AMBER *ff* is known to perform very well for kink-turns.[42-44] Considering the importance of the kink-turn motif it indicates a major disbalance of the DESRES *ff* (our implementation), tentatively suggesting it may have been over-fitted in favor of a straight A-RNA. Further, some of the attempted non-bonded term modifications may excessively destabilize key non-Watson-Crick base pairs. Note that Kt-7 folds in absence of proteins and thus should be stably folded under our simulation conditions (see the main text).

For our next test, we have utilized the structure of the ribosomal L1 stalk RNA which is a highly-conserved rRNA element located in the large ribosomal subunit. During the ribosomal elongation cycle, the L1 stalk interacts with the deacylated tRNA molecules as they exist the ribosome during the process of proteosynthesis.[45-46] The L1 stalk RNA is a useful target for *ff* testing due to multitude of non-canonical interactions and recurrent RNA motifs it contains. Specifically, there are two kink-turns, a UNCG TL, and an internal loop. Further, via tertiary interactions, these elements form a nucleotide platform composed of six layers of non-canonical base pairs and triplets stacked on top of each other.[44] The nucleotide platform is biologically significant as it forms the binding site for the deacylated tRNA molecule. It is well established that the L1 stalk rRNA segment is quite well described by $\chi_{OL3CP}$ RNA *ff*.[44, 47] In contrast, DESRES simulations of this RNA segment were entirely unsatisfactory. In DESRES simulations, we have observed instability of AG base pairs in both of the kink-turns, consistently with our simulations of the isolated Kt-7 (see above). However, since the kink-turns in L1 stalk are additionally stabilized by tertiary interactions with the surrounding elements, unlike in the Kt-7, we did not observe their straightening on the simulation timescale. There was, however, major instability observed among the non-canonical base pair interactions within the nucleotide platform in which the bottommost and uppermost base pair layers were irreversibly lost by the end of all the attempted DESRES simulations. However, the most problematic was the description of the characteristic tertiary interaction formed between the UNCG TL and the internal loop which was permanently lost on a scale of a few nanoseconds in all attempted DESRES simulations. The loss of this tertiary interaction always started with its direct H-bonds being replaced by water-mediated interactions and progressed into a permanent spectacular degradation of the L1 stalk structure and its partial unfolding (Figure S20). In conclusion, the simulation description of the L1 stalk structure by the DESRES potential (our implementation) was very unsatisfactory, with profound structural issues affecting all key segments of the structure.

**Unbiased MD simulations with gHBfix.**

We have carried out a set of unbiased MD simulations in order to investigate an overall performance of the $\chi_{OL3CP}$ RNA *ff* combined with the gHBfix$\binom{2-OH...nbO/bO}{-0.5}\binom{NH...N}{+1.0}$ potential (termed here as gHBfix *ff*) for a wide range of RNA systems (Table S2). Initially, we simulated the 14-mer UNCG TL starting from the NMR structure (PDB ID 2KOC).[22] We reiterate that the gHBfix *ff* was not able to correct the free-energy imbalance between folded and misfolded states of the 8-mer r(gcUUCGgc) TL (see the REST2 simulations in the main text). Consistently with this, in our 10 μs-long unbiased MD simulation we observed that the 14-mer TL maintained its overall fold with the fully native conformation (all signature interactions formed) of the loop till 4.5 μs, where we identified loss of two signature $U_{L1}$(2'-OH)…$G_{L4}$(O6) and $U_{L2}$(2'-OH)…$G_{L4}$(N7) H-bonds and $G_{L4}$ residue flipped out of the loop. After ~1.2 μs-long sampling of states with $G_{L4}$ in bulge-out conformation, $G_{L4}$ spontaneously flipped back and all signature interactions of the loop were reestablished. However, the newly formed canonical loop arrangement survived for just another ~200 ns, when we observed again the previously described disruption of the native contacts and misfolded states dominated till the end of 10 μs-long MD simulation. Such a behavior confirms that the UNCG TL remains a challenge for further *ff* modifications. Nevertheless, the possible recreation of the native conformation after the disruption, which has not been observed yet during classical MD simulations, indicates that gHBfix *ff* reduces the known energy imbalance between native and misfolded conformations of the UNCG TL in RNA *ffs*.[48] Thus, we suggest that gHBfix *ff* rather than the standard *ff* should be used as the starting point for further attempts to fold the UNCG TL.

We continued testing with two RNA duplexes, i.e., r(GCACCGUUGG)$_2$ decamer (excised from the PDB ID 1QC0[49] structure) and r(UUAUAUAUAUAUAA)$_2$ tetradecamer (PDB ID 1RNA[50]). In order to probe the effect of gHBfix term, we performed two standard MD simulations of each above-mentioned duplex, one with and the other without the gHBfix term. All four simulations revealed stable behavior with RMSD of heavy atoms from the starting X-ray structure fluctuating around 1.31 ± 1.35 Å (1.27 ± 1.31 Å with the gHBfix term) and 2.15 ± 2.20 Å (1.86 ± 1.90 Å with the gHBfix term) for the decamer and tetradecamer, respectively. These RMSD numbers are fully comparable with those seen in A-RNA simulations with standard $\chi_{OL3CP}$ RNA *ff* without gHBfix; for a detailed analysis of benchmark A-RNA simulations see Ref. [51]. Base-pair breathing (fraying) of GC pairs at the end of helices occurred only in few frames from the entire 1 μs-long MD simulation with the gHBfix *ff*. AU base-pair fraying was also significantly reduced in comparison with standard $\chi_{OL3CP}$ simulations with the OPC water model and the population of base-pair open state was ~1.5% for both terminal AU base pairs (Table S3). Note that accurate (unambiguous) experimental data for the quantification of frayed structures are not available for RNA sequences.[52] However, recent computational study of fraying in RNA and DNA sequences showed that base-pair openings are usually followed by formation of likely spurious long-lived noncanonical interactions.[52] Thus, the observed decrease in probability of terminal base-pair fraying by the gHBfix *ff* is expected to have positive effects to the behavior of RNA systems in MD simulations. The analysis of helical base-pair parameters, i.e., those measuring orientation and displacement of Watson–Crick base pairs, revealed that application of the gHBfix *ff* term did not affect the helical parameters of the simulated A-RNA duplexes. In particular, average values of base-pair roll and inclination, which serve as the main descriptors of A-RNA duplexes,[51, 53] were not affected by the gHBfix potential (Table S3). The measured roll and inclination values for both duplexes are slightly decreased in comparison with data reported in Ref. [51]. This is likely caused by the different water model[54] as the simulations in Ref. [51] were performed in TIP3P and SPC/E water models, while simulations present here used OPC water model. It is thus possible that the OPC water model may cause subtle underestimation of roll/inclination, see discussion in Ref. [51]. In summary, the gHBfix potential does not seem to affect the A-RNA structure, while the terminal base-pairs of A-RNA duplexes are thermodynamically stabilized. It is expected that gHBfix also stabilizes the A-RNA duplexes in general due to stabilization of the base pairs; this, however, was not explicitly tested. Thus, gHBfix should improve simulations of A-RNA segments compared to the $\chi_{OL3CP}$ *ff* alone.

We then applied the gHBfix *ff* for the SRL motif. In agreement with previous tests,[33-34] the overall fold of SRL remained stable. We observed comparable dynamics to the standard simulation with $\chi_{OL3CP}$ RNA *ff* (Figure S15) with a minor difference in the stability of one BPh contact. The U8G16 BPh interaction appeared to be weakened in the gHBfix simulation. Although the structural stability of both the whole GpU platform and the neighboring *trans* Hoogsteen/Sugar Edge A9G16 base pair were unaffected, the U8G16 BPh contact was identified as indirect, i.e., water bridged, for most of the time by the gHBfix *ff* (Figure S15). The difference is either due to some indirect effects or due to sampling, since the BPh interactions are not modified by the current version of the gHBfix. The results are suggesting that an adjustment of BPh interactions could be attempted in future (see the main text for details).

As the next systems, we simulated folded RNA motifs containing protonated nucleobases near either a cleavage or a ligand binding site. The HrRz system revealed stable dynamics, with all catalytically important contacts remaining to be firmly established. Results from the simulation with the gHBfix *ff* were thus very similar to the standard simulations with the $\chi_{OL3CP}$ *ff* (Figure S16). We followed with simulations of structural dynamics of the ligand-bound (Holo state, PDB ID 3GCA[55]) and ligand-free (Apo state, PDB ID 3Q51[56]) preQ class I riboswitch, which regulates biosynthesis of the hypermodified nucleoside queuosine by sensing its metabolic precursor 7-amino-methyl-7-deazaguanine (preQ).[55, 57] Both states contain a protonated cytosine nucleotide (C7H$^+$).[26] In general, the gHBfix *ff* describes both Apo and Holo states in agreement with the standard $\chi_{OL3CP}$ RNA *ff*, i.e., all canonical, noncanonical and triplet base pairings remained stable. Nucleotides forming the base-pair quadruplet and the ligand binding pocket in the Holo state were fluctuating around their starting positions. We observed that both base pairs of the short P2 stem remained stable, providing less dynamical motions in comparison with the original study using the $\chi_{OL3CP}$.[26] This is consistent with the stabilization of H-bonded base pairs by the gHBfix *ff*. On the other hand, the gHBfix *ff* did not reveal the spontaneous formation of the GpU platform during the simulation with Apo state reported in our earlier study as the G11(N2H)…U12(O4) H-bond was not established (see the original study[26] for details). It should be noted that the intrinsic dynamics reported in the original paper[26] has not yet been supported by experiment and thus the quick formation of the GpU platform in Ref. [26] could be a simulation artifact. Thus, we consider the description of preQ riboswitch by gHBfix *ff* as a satisfactory. The differences in the local dynamics between the simulations may reflect some sampling uncertainties and with presently available experimental data it is not possible to tell which *ff* variant is more realistic.

We then simulated the RNA kissing loop complex of the HIV-1 virus dimerization initiation site (PDB ID 1ZCI[58]), which is composed of two identical strands that interact by forming 6-base-pair long interstrand helix. The kissing loop is a medium sized system containing canonical base pairs with a limited number of noncanonical structural features and is often used as a standard benchmark for *ff* testing along with the SRL motif.[59] Experimental structures agree in the overall fold of the complex, but differ in the orientation of two unpaired adenines from each strand. Those are either located stacked inside helices according to NMR structures[60-61] or exposed to the solvent (bulged out of the helices) according to X-ray structures.[58] Previous MD simulations with the $\chi_{OL3CP}$ RNA *ff* classified several substates of unpaired adenines and concluded that without any external interaction, adenines preferably adopt bulged-in conformation.[59] However, the simulation results were far from being converged.[59] With gHBfix *ff* we also observed stable behavior of the complex, where all base pairs were maintained during 1 μs-long unbiased MD simulation. The unpaired residues sampled several conformations in agreement with the original study.[59] Within the limits of sampling it looks that gBHfix and $\chi_{OL3CP}$ have similar performance for the kissing complex. Note that the ambiguity of the experimental data (in detail discussed in Ref. [59]) precludes to use the dynamics of the bulge-out bases to benchmark *ffs*.

Next, we simulated a tetrameric structure composed of four GGG strands forming three stacked G-quartets (RNA G-quadruplex, based on PDB ID 3IBK[62]). A channel formed in the middle of G-stem binds two potassium cations. The simulated three-tetrad stem was prepared by removing flanking and loop nucleotides, whereas the channel cations were kept in place. We observed that the gHBfix *ff* did not affect the structural stability of the RNA G-quadruplex during the 1 μs-long standard MD simulation compared to $\chi_{OL3CP}$ simulations. All base pairs were preserved and both cations remained bound in the channel. However, the gHBfix *ff* did not eliminate the bifurcated hydrogen bonds (BHB) pattern identified in $\chi_{OL3CP}$ simulations,[63] i.e., the preference of bifurcated G(N2H)/G(N1H)…G(N7) H-bonds and the partial weakening of G(N2H)…G(O6) H-bonds, which results in slight deformation of tetrad geometries. The bifurcated H-bonding in G-tetrads is a known local problem that occurs with all versions of the AMBER *ff* in simulations of DNA and RNA quadruplexes and which does not preclude successful application of MD simulations to quadruplexes, as in detail discussed in Ref. [59].

The simulations of the kink-turn Kt-7, the L1 stalk rRNA and Neomycin-sensing riboswitch (NSR) utilizing the gHBfix potential have revealed identical behavior (within the sampling limitations) as reported in earlier simulation studies of these molecules using the $\chi_{OL3CP}$ RNA *ff*.[44, 64] Namely, all the signature interactions were fully maintained in gHBfix *ff* simulations of the kink-turn Kt-7. The individual segments of the L1 stalk were stable as well, including the two kink-turns and the non-canonical nucleotide platform. Lastly, the use of gHBfix modestly lowered the range of some of the NOE distance violations reported for simulations of the NSR riboswitch,[64] although it did not entirely eliminate them. Most significantly, it did not introduce any new distance violations. Thus, for the NSR riboswitch, the gHBfix *ff* may actually bring some improvement. Let us, however, point out that all three systems are well behaving in both the original $\chi_{OL3CP}$ and gHBfix versions of the RNA *ff*.

In summary, no adverse structural side-effects of the gHBfix potential were noticed in our set of standard (unbiased) simulations of a number of folded RNAs.

**Unbiased MD simulations with *ff* modification by Pak and coworkers.**

Popular approach towards *ff* improvement is adjusting Lennard-Jones parameters with simultaneous modifications of the Lennard-Jones combining rules via nonbonded fix (NBfix, for a recent review see Ref. [65]) to balance the RNA-solvent interaction. For a more detailed discussion see also the main text. An example of such approach is the recent *ff* modification published by Pak and coworkers (denoted here as Pak-NBfix),[66] which was developed with the attempt to improve the structural description of TNs during folding simulations. Although the presented results for TNs and one TL motif were encouraging,[66] the authors did not present any tests on more complicated RNA motifs.

Here, we performed a set of classical MD simulations of the Kink-turn Kt-7, where we applied the Pak-NBfix correction. The Kt-7 structure was very poorly described by simulations utilizing the Pak-NBfix reparameterization. We observed almost immediate and irreversible loss of the signature A-minor interaction in all attempted simulations. In majority of the simulations, we also at some point observed loss of the characteristic bent shape and complete straightening of the structure (Figure S19). Thus, we were unable to obtain any stable simulations of the Kt-7 structure when utilizing the Pak-NBfix *ff*. Similarly to the Kt-7 simulations utilizing the DESRES potential (see above), the attempted non-bonded term tuning appears to excessively destabilize the non-canonical base-base and sugar-base interactions such, as the A-minor interaction. Based on the entirely unsatisfactory Kt-7 results, no further testing of this *ff* has been attempted.

**Program to generate AMBER inputs for MD simulation with the gHBfix potential**

We are attaching the C++ program (gHBfix.pdf) that can be used to generate AMBER restraint input files for MD simulations with the external potential selectively influencing specific types of H-bonds (gHBfix). We note that the gHBfix potential (as a method) can be combined with any existing RNA *ff*. The specific parameters used in the present study were derived and tested in combination with the standard $\chi_{OL3CP}$ *ff*,[6-9] adjusted in addition by vdW modification of phosphate oxygen developed by Case et al.,[10] and OPC water model.[11] The gHBfix C++ code is designed to generate the input file for any RNA system and requires certain options to be specified, i.e., (i) the RNA sequence, (ii) scaling parameter $\lambda$ (required for REST2 simulation, otherwise it should be set to 1.0), and (iii) values in energy ($\eta$, kcal/mol) to support/penalize particular interactions. For each particular gHBfix term, the proton donor and proton acceptor types have to be specified. The current version of the program supports following proton donors: (i) –NH groups of nucleobases including both hydrogens of exocyclic amino groups and (ii) –OH hydroxyl groups including the 2'-OH groups as well as terminal 3'-OH and 5'-OH groups. Following proton acceptors are supported: (i) phosphate nbOs, (ii) phosphate bOs, i.e., O3' and O5' except for the terminal ones, (iii) oxygens of hydroxyl groups (acting as proton acceptors), (iv) ribose O4' oxygens, (v) carbonyl oxygens of nucleobases, and (vi) iminogroups of nucleobases capable to accept the H-bond (see the Table 1 in the main text). In summary, the gHBfix potential might be applied to twelve different types of H-bonding pair-wise interactions, i.e., all possible combinations of two types of H-bond donors with six types of H-bond acceptors (see the Table 2 in the main text). Note that in order to prevent affecting the dihedral terms of the *ff*, the gHBfix potential is not applied to any pair-wise contacts that are separated by three covalent bonds between the heavy atoms of a given type of H-bond. This includes H-bonding between 2'-OH group and neighboring O3' oxygen on the same ribose, and all –NH donor and N/O acceptor pairs within the same nucleobase. All these interactions are automatically excluded by the program. The types of interactions and the parameters can be readily selected by the users, i.e., they may differ from those tested in the present work.

Additionally, the gHBfix code contains options for printing input file for CPPTRAJ[67] (option -cpptraj, useful for trajectory analysis) and for PLUMED[68] (option -plumed), which allows the direct usage of the gHBfix potential in GROMACS[69] MM package.

**AMBER input files containing DESRES parameters**

We are attaching AMBER input files for MD simulation with our implementation of the DESRES potential[13] (see the attachment called desres.zip). All files with parameters, including those specifying the TIP4P-D water model,[14] are loaded into the tLEaP module of AMBER 16 program package[5] by sourcing the attached leaprc.RNA.Shaw file.

### Supporting Tables

**Table S1**: Overview of all performed enhanced sampling simulations of TN and TL motifs.

| Motif | RNA *ff* | Modification [a] | Replicas | Length [μs] |
|---|---|---|---|---|
| r(GACC) | $\chi_{\text{OL3CP}}$ | - | 8 | 10 |
| r(GACC) | $\chi_{\text{OL3CP}}$ | $\text{gHBfix}\binom{\text{2−OH…nbO}}{-1.0}$ | 8 | 10 |
| r(GACC) | $\chi_{\text{OL3CP}}$ | $\text{gHBfix}\binom{\text{2−OH…nbO/bO}}{-1.0}$ | 8 | 10 |
| r(GACC) | $\chi_{\text{OL3CP}}$ | $\text{gHBfix}\binom{\text{2−OH…nbO/bO/O4}}{-1.0}$ | 8 | 10 |
| r(GACC) | $\chi_{\text{OL3CP}}$ | $\text{gHBfix}\binom{\text{2−OH…nbO}}{-0.5}$ | 8 | 10 |
| r(GACC) | $\chi_{\text{OL3CP}}$ | $\text{gHBfix}\binom{\text{2−OH…nbO/bO}}{-0.5}$ | 8 | 10 |
| r(gcGAGAgc) | $\chi_{\text{OL3CP}}$ | $\text{gHBfix}\binom{\text{2−OH…nbO/bO}}{-0.5}$ | 12 | 10 |
| r(gcGAGAgc) | $\chi_{\text{OL3CP}}$ | $\text{gHBfix}\binom{\text{2−OH…nbO/bO}}{-0.5}\binom{\text{NH…N}}{+1.0}\binom{\text{NH…O}}{+1.0}$ | 12 | 10 |
| r(gcGAGAgc) | $\chi_{\text{OL3CP}}$ | $\text{gHBfix}\binom{\text{2−OH…nbO/bO}}{-0.5}\binom{\text{NH…N}}{+1.0}\binom{\text{NH…O}}{+0.5}$ | 12 | 10 |
| r(gcGAGAgc) | $\chi_{\text{OL3CP}}$ | $\text{gHBfix}\binom{\text{2−OH…nbO/bO}}{-0.5}\binom{\text{NH…N}}{+1.0}$ | 12 | 10 |
| r(gcGAGAgc) | $\chi_{\text{OL3CP}}$ | $\text{gHBfix}\binom{\text{2−OH…nbO/bO}}{-0.5}\binom{\text{NH…N}}{+0.5}$ | 12 | 10 |
| r(gcGAGAgc) | $\chi_{\text{OL3CP}}$ | $\text{gHBfix}\binom{\text{2−OH…nbO/bO}}{-0.5}\binom{\text{NH…N}}{+0.5}\binom{\text{NH…O}}{+1.0}$ | 12 | 10 |
| r(gcGAGAgc) | $\chi_{\text{OL3CP}}$ | $\text{gHBfix}\binom{\text{2−OH…nbO/bO}}{-0.5}\binom{\text{NH…N}}{+0.5}\binom{\text{NH…O}}{+0.5}$ | 12 | 10 |
| r(gcGAGAgc) | $\chi_{\text{OL3CP}}$ | $\text{gHBfix}\binom{\text{2−OH…nbO/bO}}{-0.5}\binom{\text{NH…O}}{+0.5}$ | 12 | 10 |
| r(gcGAGAgc) | $\chi_{\text{OL3CP}}$ | $\text{gHBfix}\binom{\text{2−OH…nbO/bO}}{-0.5}\binom{\text{NH…O}}{+1.0}$ | 12 | 10 |
| r(GACC) | $\chi_{\text{OL3CP}}$ | $\text{gHBfix}\binom{\text{2−OH…nbO/bO}}{-0.5}\binom{\text{NH…N}}{+1.0}\binom{\text{NH…O}}{+1.0}$ | 8 | 10 |
| r(GACC) | $\chi_{\text{OL3CP}}$ | $\text{gHBfix}\binom{\text{2−OH…nbO/bO}}{-0.5}\binom{\text{NH…N}}{+1.0}\binom{\text{NH…O}}{+0.5}$ | 8 | 10 |
| r(GACC) | $\chi_{\text{OL3CP}}$ | $\text{gHBfix}\binom{\text{2−OH…nbO/bO}}{-0.5}\binom{\text{NH…N}}{+1.0}$ | 8 | 10 |
| r(GACC) | $\chi_{\text{OL3CP}}$ | $\text{gHBfix}\binom{\text{2−OH…nbO/bO}}{-0.5}\binom{\text{NH…N}}{+0.5}$ | 8 | 10 |
| r(GACC) | $\chi_{\text{OL3CP}}$ | $\text{gHBfix}\binom{\text{2−OH…nbO/bO}}{-0.5}\binom{\text{NH…N}}{+0.5}\binom{\text{NH…O}}{+1.0}$ | 8 | 10 |
| r(GACC) | $\chi_{\text{OL3CP}}$ | $\text{gHBfix}\binom{\text{2−OH…nbO/bO}}{-0.5}\binom{\text{NH…N}}{+0.5}\binom{\text{NH…O}}{+0.5}$ | 8 | 10 |
| r(GACC) | $\chi_{\text{OL3CP}}$ | $\text{gHBfix}\binom{\text{2−OH…nbO/bO}}{-0.5}\binom{\text{NH…O}}{+1.0}$ | 8 | 10 |
| r(GACC) | $\chi_{\text{OL3CP}}$ | $\text{gHBfix}\binom{\text{2−OH…nbO/bO}}{-0.5}\binom{\text{NH…O}}{+0.5}$ | 8 | 10 |
| r(GACC) | $\chi_{\text{OL3CP}}$ | NBfix as $\text{gHBfix}\binom{\text{2−OH…nbO/bO}}{-0.5}\binom{\text{NH…O}}{+1.0}$ [b] | 8 | 10 |
| r(gcUUCGgc) | $\chi_{\text{OL3CP}}$ | HBfix(10 signature interactions) [c] | 64 | 10 |
| r(gcUUCGgc) | $\chi_{\text{OL3CP}}$ | $\text{gHBfix}\binom{\text{2−OH…nbO/bO}}{-0.5}\binom{\text{NH…N}}{+1.0}\binom{\text{NH…O}}{+1.0}$ | 12 | 10 |
| r(gcUUCGgc) | $\chi_{\text{OL3CP}}$ | $\text{gHBfix}\binom{\text{2−OH…nbO/bO}}{-0.5}\binom{\text{NH…N}}{+1.0}$ | 12 | 7 |
| r(gcUUCGgc) | $\chi_{\text{OL3CP}}$ | $\text{gHBfix}\binom{\text{2−OH…nbO/bO}}{-0.5}\binom{\text{NH…N}}{+0.5}$ | 12 | 5 |
| r(gcUUCGgc) | $\chi_{\text{OL3CP}}$ | $\text{gHBfix}\binom{\text{2−OH…nbO/bO}}{-0.5}\binom{\text{NH…N}}{+1.0}\binom{\text{NH…O}}{+1.0}\binom{\text{2−OH…N/O}}{+0.5}$ | 12 | 6.6 |
| r(gcUUCGgc) | $\chi_{\text{OL3CP}}$ | $\text{gHBfix}\binom{\text{2−OH…nbO/bO}}{-0.5}\binom{\text{NH…N}}{+1.0}\binom{\text{NH…O}}{+1.0}\binom{\text{2−OH…N/O}}{+1.0}$ | 12 | 7.2 |
| r(gcUUCGgc) | $\chi_{\text{OL3CP}}$ | $\text{gHBfix}\binom{\text{2−OH…nbO/bO}}{-0.5}\binom{\text{NH…N}}{+1.0}\binom{\text{NH…O}}{+1.0}\binom{\text{NH…O2}}{-0.5}\binom{\text{2−OH…O2/O4}}{-0.5}$ | 12 | 6 |
| r(gcUUCGgc) | $\chi_{\text{OL3CP}}$ | $\text{gHBfix}\binom{\text{2−OH…nbO/bO}}{-0.5}\binom{\text{NH…N}}{+1.0}\binom{\text{2−OH…N/O}}{+0.5}$ | 12 | 10 |
| r(gcUUCGgc) | $\chi_{\text{OL3CP}}$ | $\text{gHBfix}\binom{\text{2−OH…nbO/bO}}{-0.5}\binom{\text{NH…N}}{+1.0}\binom{\text{NH…O}}{+0.5}\binom{\text{2−OH…N/O}}{+0.5}\binom{\text{NH…O2}}{-0.5}\binom{\text{2−OH…O2/O4}}{-0.5}$ | 12 | 5 |
| r(gcUUCGgc) | $\chi_{\text{OL3CP}}$ | $\text{gHBfix}\binom{\text{2−OH…nbO/bO}}{-0.5}\binom{\text{NH…N}}{+0.5}\binom{\text{NH…O}}{+0.5}\binom{\text{2−OH…N/O}}{+0.5}\binom{\text{NH…O2}}{-0.5}\binom{\text{2−OH…O2/O4}}{-0.5}$ | 12 | 10 |
| r(CAAU) | $\chi_{\text{OL3CP}}$ | $\text{gHBfix}\binom{\text{2−OH…nbO/bO}}{-0.5}\binom{\text{NH…N}}{+1.0}$ | 8 | 10 |
| r(CCCC) | $\chi_{\text{OL3CP}}$ | $\text{gHBfix}\binom{\text{2−OH…nbO/bO}}{-0.5}\binom{\text{NH…N}}{+1.0}$ | 8 | 10 |
| r(AAAA) | $\chi_{\text{OL3CP}}$ | $\text{gHBfix}\binom{\text{2−OH…nbO/bO}}{-0.5}\binom{\text{NH…N}}{+1.0}$ | 8 | 10 |

| | | | | |
|---|---|---|---|---|
| r(UUUU) | $\chi_{\mathrm{OL3CP}}$ | gHBfix$\left(^{\mathrm{2-OH...nbO/bO}}_{-0.5}\right)\left(^{\mathrm{NH...N}}_{+1.0}\right)$ | 8 | 10 |
| r(GACC) | DESRES [d] | - | 8 | 10 |
| r(gcGAGAgc) | DESRES | - | 12 | 4 |
| r(gcGAGAgc) [e] | DESRES | - | 12 | 2 |
| r(gcUUCGgc) | DESRES | - | 12 | 2 |
| r(gcUUCGgc) [e] | DESRES | - | 12 | 2 |

[a] additional potential function tuning the stability of specific types of H-bonds (gHBfix, see Methods).
[b] NBfix *ff* reparameterization was prepared in a way to be comparable with the gHBfix$\left(^{\mathrm{2-OH...nbO/bO}}_{-0.5}\right)\left(^{\mathrm{NH...N}}_{+1.0}\right)$ potential (see the section Details about the relation between the gHBfix and NBfix approaches).
[c] T-REMD simulation using structure-specific version of HBfix (see Methods).
[d] implemented parameters as described in Ref. [13].
[e] all replicas started from the folded (native-like) state.

**Table S2:** Overview of performed unbiased MD simulations using various RNA systems with standard $\chi_{\mathrm{OL3CP}}$ RNA *ff*, $\chi_{\mathrm{OL3CP}}$ RNA *ff* with the gHBfix$\left(^{\mathrm{2-OH...nbO/bO}}_{-0.5}\right)\left(^{\mathrm{NH...N}}_{+1.0}\right)$ potential, and recent DESRES[13] *ff* and Pak-NBfix[66] modification.

| Motif | PDB ID | RNA *ff* | Modification [a] | Simulations [b] | Length [μs] |
|---|---|---|---|---|---|
| SRL | 3DW4 | $\chi_{\mathrm{OL3CP}}$ | - | 1 | 1 |
| HrRz [c] | 2OUE | $\chi_{\mathrm{OL3CP}}$ | - | 1 | 1 |
| RNA duplexes | 1QC0, 1RNA | $\chi_{\mathrm{OL3CP}}$ | - | 1, 1 | 1 |
| RNA G-quadruplex [d] | 3IBK | $\chi_{\mathrm{OL3CP}}$ | - | 1 | 1 |
| UNCG TL (14-mer) | 2KOC | $\chi_{\mathrm{OL3CP}}$ | gHBfix | 1 | 10 |
| SRL | 3DW4 | $\chi_{\mathrm{OL3CP}}$ | gHBfix | 1 | 1 |
| HrRz [c] | 2OUE | $\chi_{\mathrm{OL3CP}}$ | gHBfix | 1 | 1 |
| Kt-7 *H.m* [e] | 1S72 | $\chi_{\mathrm{OL3CP}}$ | gHBfix | 4 | 1 |
| L1-stalk | 3U4M | $\chi_{\mathrm{OL3CP}}$ | gHBfix | 4 | 1 |
| NSR | 2N0J | $\chi_{\mathrm{OL3CP}}$ | gHBfix | 4 | 1 |
| preQ [f] | 3GCA, 3Q51 | $\chi_{\mathrm{OL3CP}}$ | gHBfix | 1, 1 | 1 |
| Kissing loop | 1ZCI | $\chi_{\mathrm{OL3CP}}$ | gHBfix | 1 | 1 |
| RNA duplexes | 1QC0, 1RNA | $\chi_{\mathrm{OL3CP}}$ | gHBfix | 1, 1 | 1 |
| RNA G-quadruplex [d] | 3IBK | $\chi_{\mathrm{OL3CP}}$ | gHBfix | 1 | 1 |
| UNCG TL (8-mer) | 1F7Y | DESRES [g] | - | 1 | 1 |
| UNCG TL (14-mer) | 2KOC | DESRES | - | 5 | 10 |
| SRL | 3DW4 | DESRES | - | 1 | 1 |
| HrRz [c] | 2OUE | DESRES | - | 3 | 1 |
| Kt-7 *H.m* [e] | 1S72 | DESRES | - | 4 | 1 |
| L1-stalk | 3U4M | DESRES | - | 3 | 1, 1, 0.41 |
| Kt-7 *H.m* [e] | 1S72 | $\chi_{\mathrm{OL3CP}}$ | Pak-NBfix [h] | 6 | 1 |

[a] additional potential function selectively influencing specific types of H-bonds (gHBfix).
[b] number of independent simulations.
[c] simulation performed with the N1-protonated form of A38 (A38H$^+$) required for the stability of the active site in MD simulations.[27]
[d] the simulated RNA G-quadruplex three-tetrad stem was prepared by removing flanking and loop nucleotides, whereas the channel cations were kept in place.
[e] the simulated Kt-7 structure was obtained by excising the residues 76-83 and 91-101 from the structure of the large ribosomal subunit of *Haloarcula marismortui.*
[f] preQ riboswitch was simulated both with and without ligand in the binding pocket (Holo and Apo states).
[g] see attached files (desres.zip) for implemented parameters as described in Ref. [13]
[h] NBfix *ff* reparameterization as suggested by Pak and coworkers.[66]
Abbreviations: SRL (Sarcin-Ricin Loop RNA motif), HrRz (Hairpin Ribozyme), NSR (Neomycin-sensing riboswitch), Kt-7 (Kink-turn 7), preQ (preQ riboswitch), Kissing-loop (HIV-1 RNA DIS Kissing Complex).

**Table S3:** The analysis of helical base-pair parameters and base-pair fraying for two simulated duplexes, i.e., decamer r(GCACCGUUGG)$_2$ and tetradecamer r(UUAUAUAUAUAUAA)$_2$. Averaged values of base-pair roll and inclination from 1 μs-long $\chi_{OL3CP}$ and gHBfix *ff* MD simulations were obtained by CPPTRAJ[67] and calculated for the inner six and ten base-pair segments of the decamer and tetradecamer, respectively. The frequency of fraying was estimated by calculating Root Mean Square Deviation (RMSD, considering nucleobase atoms of each terminal base pair) with RMSD cutoff 1.35 Å.

| helical parameters | $\chi_{OL3CP}$ *ff* | | gHBfix *ff* | |
|---|---|---|---|---|
| | inclination [°] | roll [°] | inclination [°] | roll [°] |
| r(gcACCGUUgg)$_2$ | 14.1 | 7.6 | 14.2 | 7.7 |
| r(uuAUAUAUAUAUaa)$_2$ | 18.6 | 10.7 | 18.2 | 10.5 |

| fraying analysis | $\chi_{OL3CP}$ *ff* | | gHBfix *ff* | |
|---|---|---|---|---|
| | top [%] [a] | bottom [%] [b] | top [%] | bottom [%] |
| r(gcACCGUUgg)$_2$ | 0.2 | 0.5 | 0.0 | 0.0 |
| r(uuAUAUAUAUAUaa)$_2$ | 14.6 | 47.3 | 1.3 | 1.5 |

[a] population of frayed states for the top base pair, i.e., nucleotides 1-20 and 1-28 for the decamer and tetradecamer, respectively.

[b] population of frayed states for the bottom base pair, i.e., nucleotides 10-11 and 14-15 for the decamer and tetradecamer, respectively.

**Table S4:** The modified Lennard-Jones combining rules (NBfix) obtained as an attempt to mimic changes of interaction-energy curves introduced by the gHBfix potential. We aimed to find NBfix parameters for particular Lennard-Jones pairs that would provide approximately +1.0 kcal/mol stabilization for all possible –NH…N– interactions and approximately –0.5 kcal/mol destabilization of all possible SPh interactions.

| gHBfix [a] | η [b] (kcal/mol) | Atom pairs modified | *R* (Å) [c] | | *ε* (kcal/mol) [d] | |
|---|---|---|---|---|---|---|
| | | | $\chi_{OL3CP}$ [e] | NBfix as gHBfix [f] | $\chi_{OL3CP}$ [e] | NBfix as gHBfix [f] |
| NH…N | +1.0 | H…NC, H…NB | 2.4240 | 1.8740 | 0.0517 | 0.0517 |
| | | NA…NB, NA…NC, N2…NB, N2…NC | 3.6480 | 3.5980 | 0.1700 | 0.1700 |
| 2-OH…bO | –0.5 | HO…OR | 1.7718 [g] | 2.2578 | 0 | 0.1304 |
| 2-OH…nbO | –0.5 | HO…OP | 1.7493 [g] | 2.0693 | 0 | 0.1449 |

[a] interactions modified by the gHBfix potential

[b] total energetic support or penalty for each H-bond interaction by the gHBfix potential function (see Methods in the main text for details).

[c] distance, where the Lennard-Jones potential for the interaction of atoms *i* and *j*, $R_{i,j}$, is exactly zero. Parameters for particular Lennard-Jones pairs derived as: $R_{i,j} = R_i + R_j$

[d] depth of the potential well for the interaction of atoms *i* and *j*, $\varepsilon_{i,j}$. Parameters for particular Lennard-Jones pairs derived as: $\varepsilon_{i,j} = \sqrt{(\epsilon_i + \varepsilon_j)}$

[e] *ff*99bsc0$\chi_{OL3}$[6-9] RNA *ff* version with the vdW modification of phosphate oxygens developed by Case et al.[10] used during gHBfix simulations

[f] NBfix *ff* reparameterization was prepared in a way to be comparable with the gHBfix$\binom{2\text{-OH...nbO/bO}}{-0.5}\binom{\text{NH...N}}{+1.0}$ potential

[g] those values equal to radii of bO and nbO oxygens, respectively, as the radius of polar hydrogen is equal to zero, but they are essentially irrelevant due to zero epsilon value.

**Supporting Figures**

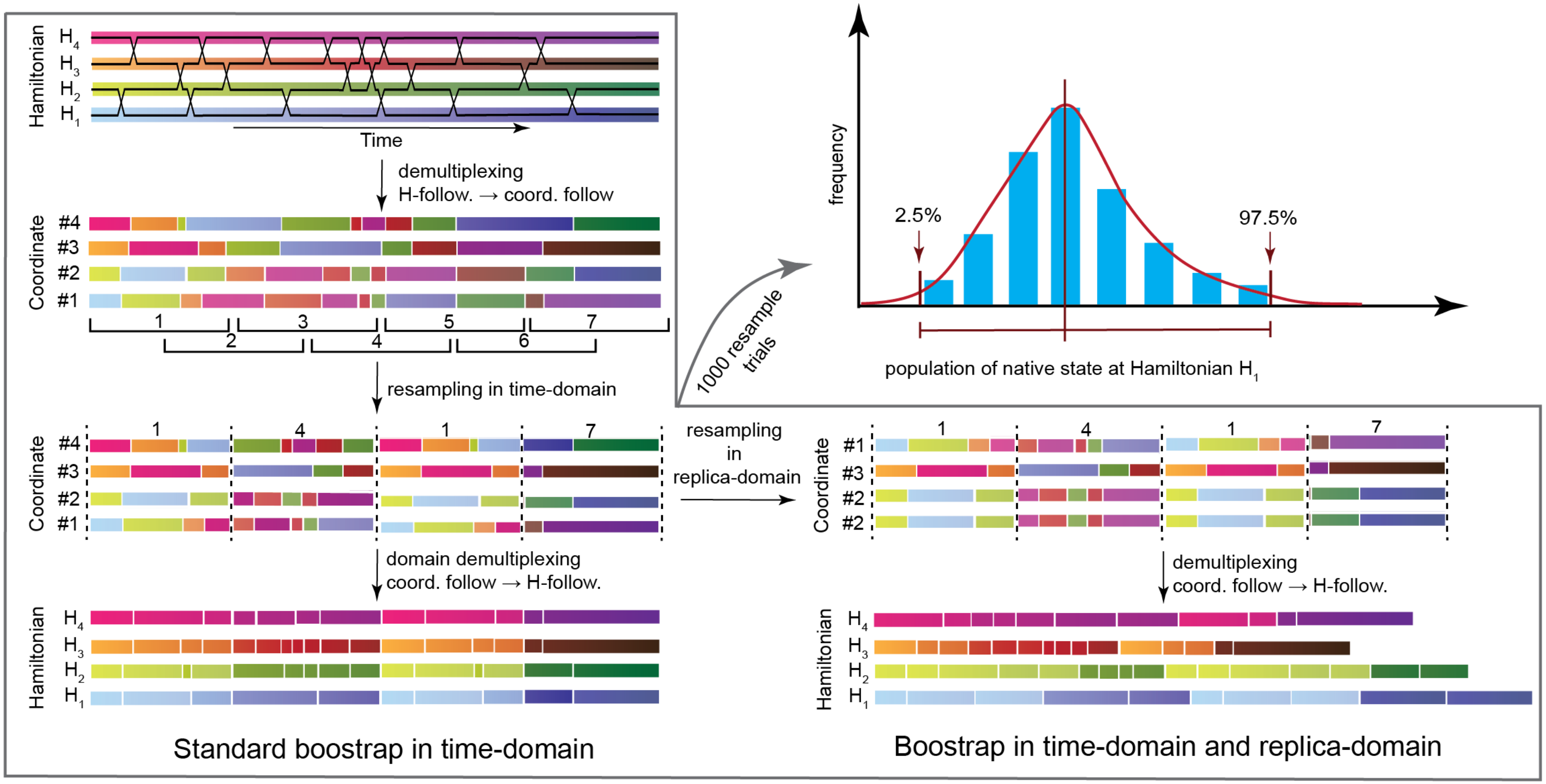

**Figure S1:** Illustrative scheme of the bootstrapping protocol, which we used for the analysis of conformational ensembles generated by REST2 and REMD simulations. The left panels illustrate the workflow of the standard bootstrap analysis in time domain only, where the time blocks are resampled. The resampled replicas are composed of randomly selected time blocks; i.e., depicted blocks on the Figure are numbered as 1, 4, 1, and 7. In our implementation of the bootstrap analysis (right-bottom panels), the replicas are also resampled following coordinates, i.e., the resampled set of trajectories is composed of randomly selected replicas (already resampled in time-domain; depicted replicas on the Figure are numbered as #2, #2, #3, and #1). Subsequently, the set of replicas (resampled both in time- and replica-domains) is resorted to follow the Hamiltonians and the population of a particular folded or misfolded states is calculated for each Hamiltonian. This resampling trials are repeated several times (in our case 1000-times), and error bars are calculated as a confidence interval covering 95% of the obtained data from the statistical ensemble (right-top panel). See the difference of obtained error bars calculated by the standard bootstrap resampling in time-domain only (Figure S2) and those by the resampling in both time- and replica-domains (Figure 4 in the main text).

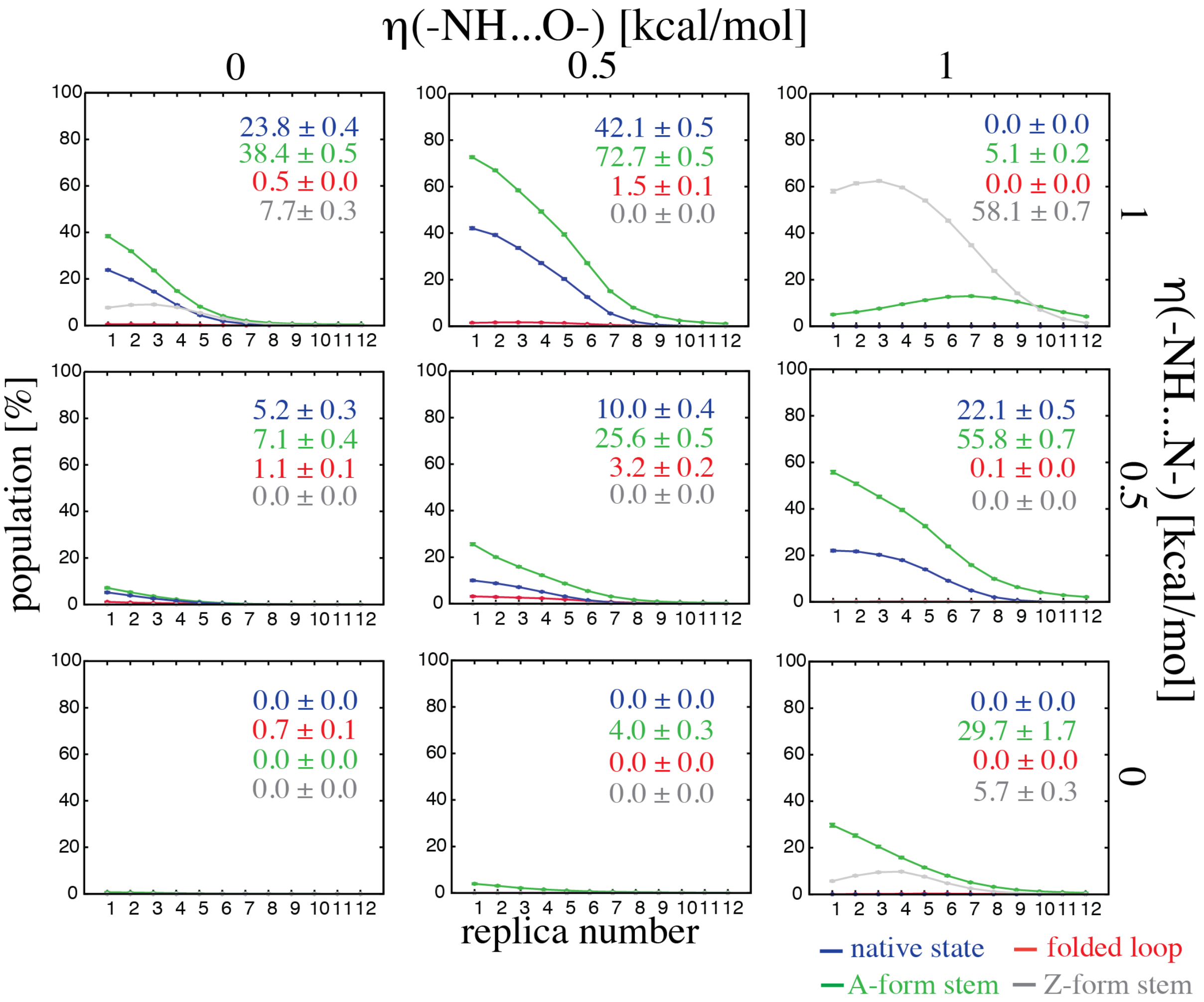

**Figure S2:** Same data as shown on Figure 4 in the main text but with errors estimated by bootstrapping in time-domain only (see Methods in the main text for details).

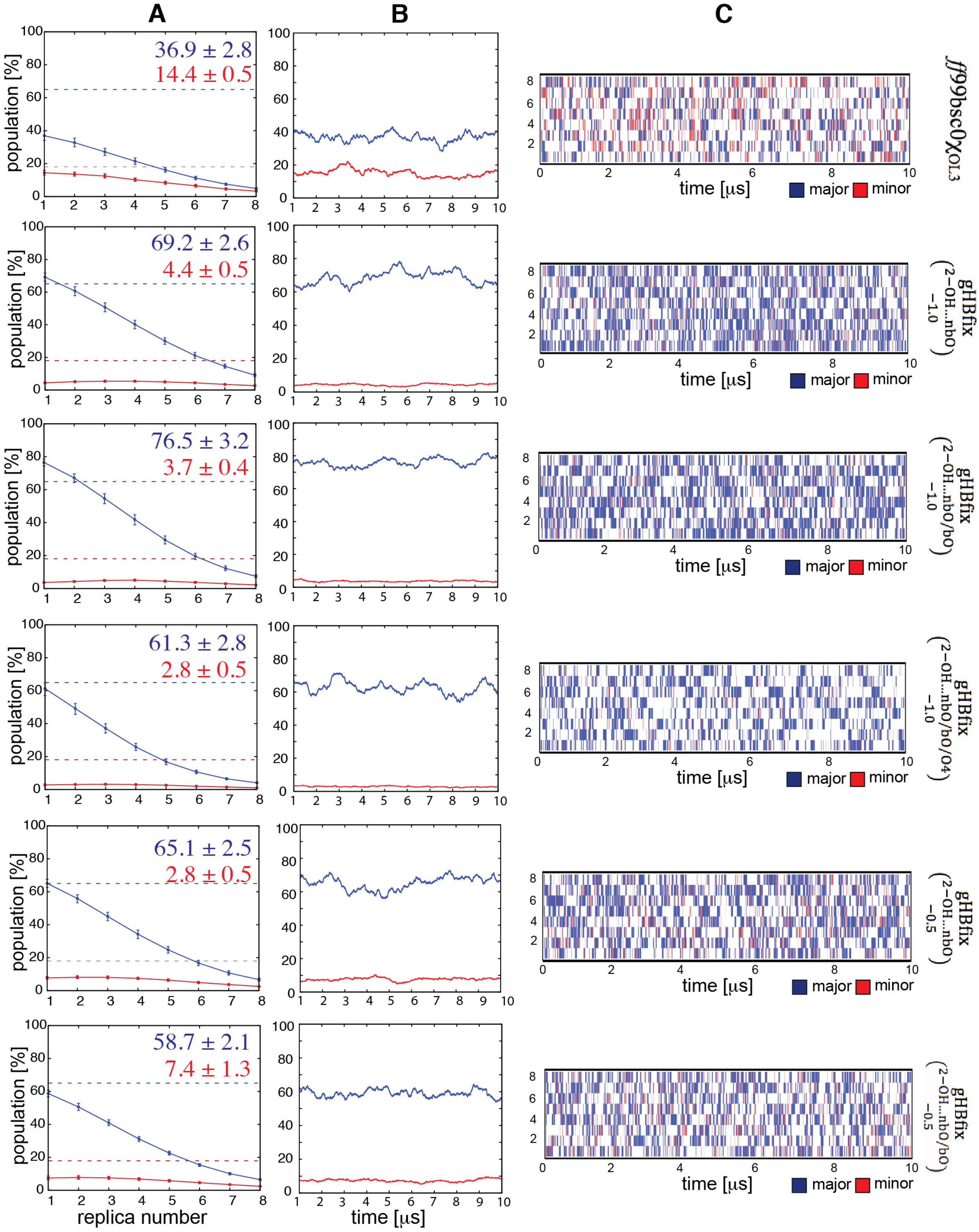

**Figure S3:** Conformational sampling and convergence of r(GACC) REST2 simulations with various gHBfix potentials penalizing formation of SPh contacts. (A) Total population of RNA A-major (blue) and RNA A-minor (red) conformers. Dashed lines indicate populations indirectly suggested by experiments.[70] The colored numbers with errors display final populations of respective conformers in the reference (unbiased) replica (T=298 K). (B) Fluctuations of A-major and A-minor states over the course of REST2 simulation obtained by time-averaging over the 1 μs window (so that the value at 1 μs corresponds to the average over states populated between time of 0 to 1 μs). (C) Time evolution of RNA A-major and A-minor conformers within all eight replicas analyzed from continuous trajectories.

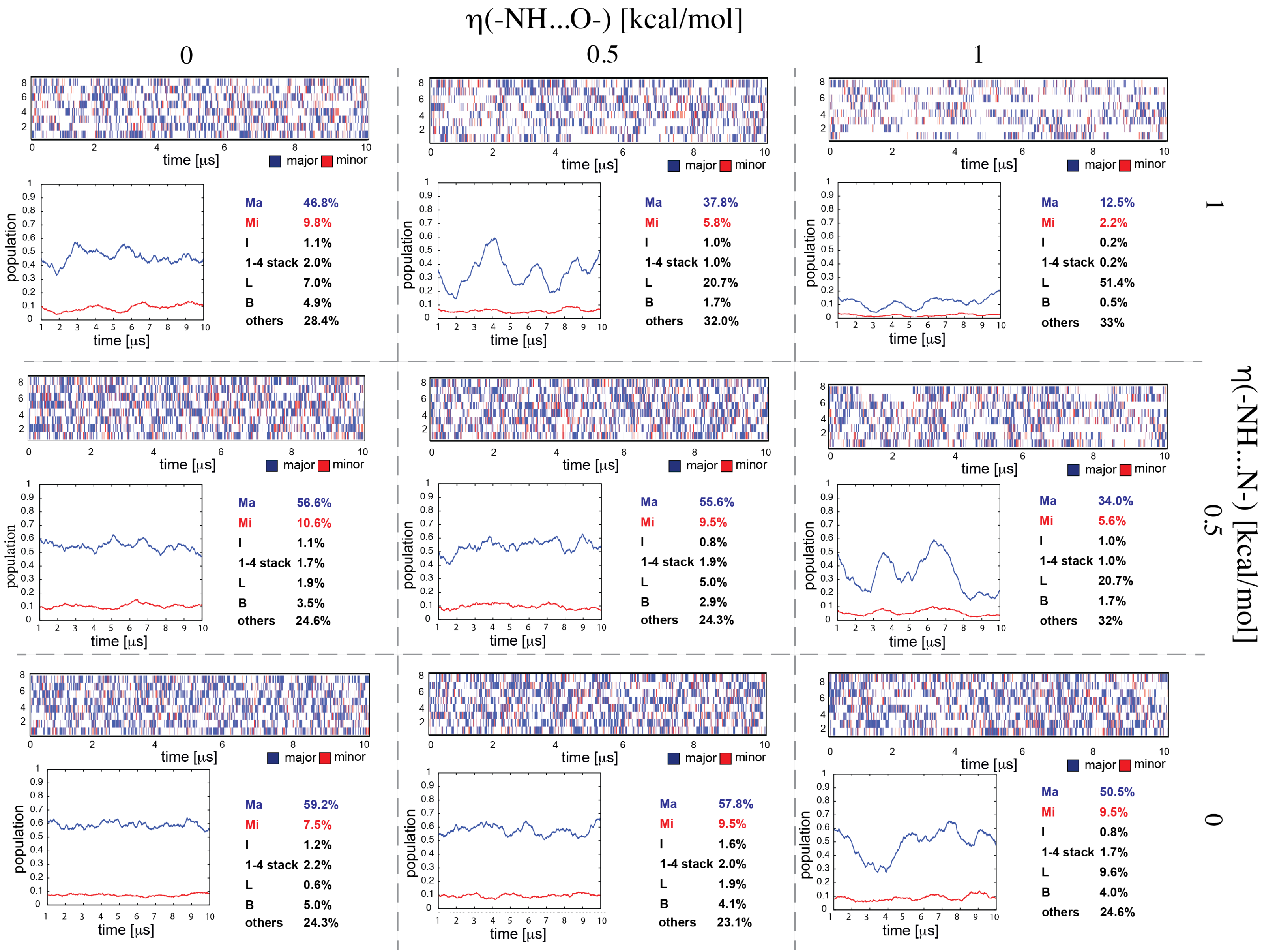

**Figure S4:** See the next page for the legend.

**Figure S4:** Conformational sampling and convergence of r(GACC) REST2 simulations with various gHBfix potentials. The gHBfix$\binom{2-\mathrm{OH}\ldots\mathrm{nbO/bO}}{-0.5}$ potential was applied in all simulations. Each panel shows: (i) time evolution of the major conformers, i.e., RNA A-major (in blue) and A-minor (in red), within all eight replicas analyzed from continuous trajectories, (ii) fluctuation of A-major and A-minor states over the course of REST2 simulation obtained by time-averaging over the 1 μs window for the reference (unbiased, T = 298 K) replica (so that the value at 1 μs corresponds to the average over states populated between time of 0 to 1 μs), and (iii) total population of all identified states for the reference replica, i.e., RNA A-major (Ma), RNA A-minor (Mi), intercalated (I), cluster with stacked terminal nucleotides (1-4 stack), loop-like structure with either 1-4 or 1-3 GC base pair formed (loop), and cluster with either second A or third C nucleotides bulged-out (B). See Methods in the main text for details about clustering procedure and Figure S6 for 3D structures of major clusters.

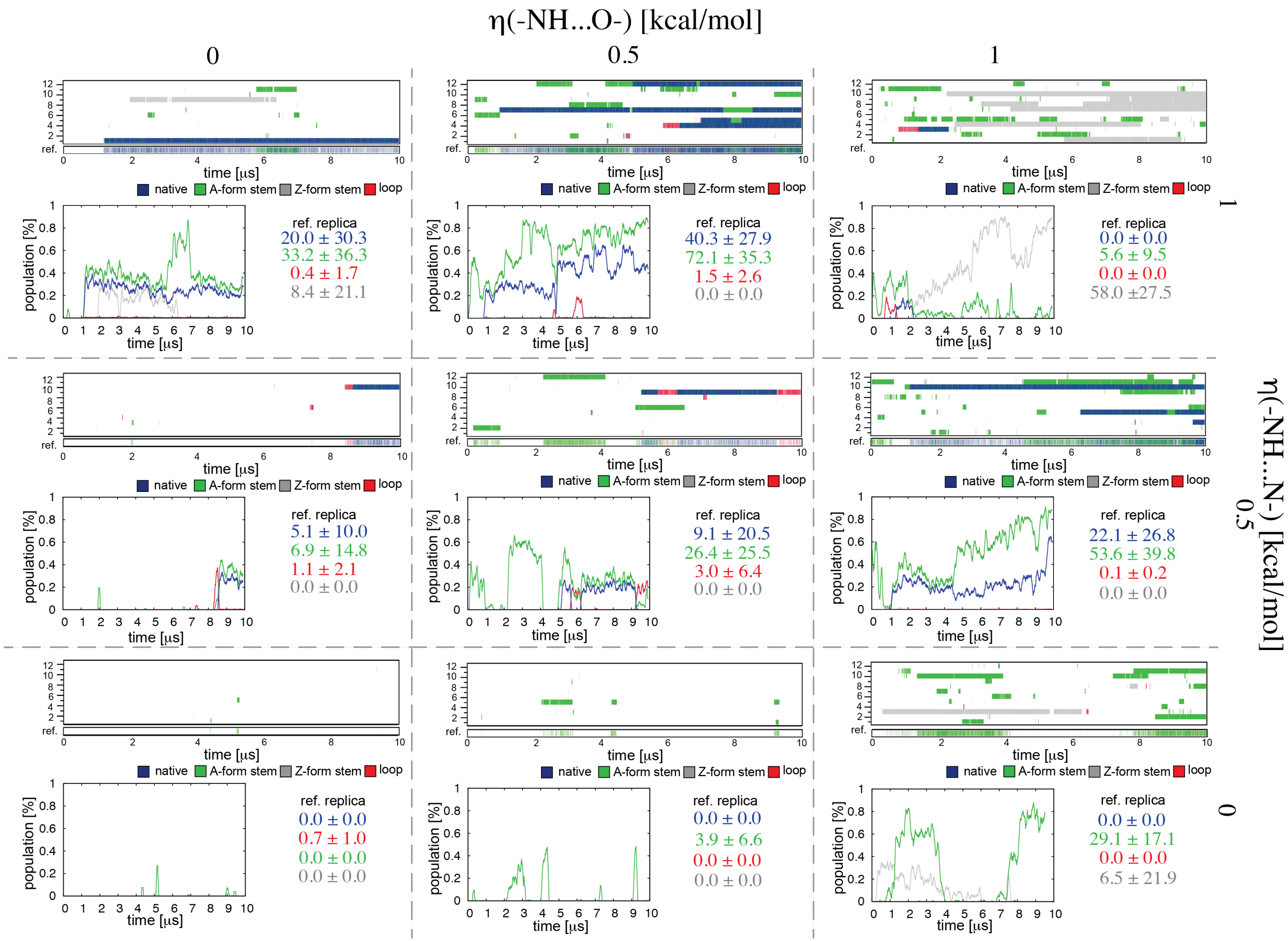

**Figure S5:** See the next page for the legend.

**Figure S5:** Conformational sampling and convergence of r(gcGAGAgc) REST2 simulations with various gHBfix potentials. The gHBfix$\binom{2-OH\ldots nbO/bO}{-0.5}$ potential was applied in all simulations. Upper plots in each panel shows the time evolution of major conformers, i.e., structures with (i) correctly folded A-form stem and loop (native states with all signature interactions formed, blue), (ii) folded A-form stem (independent of loop conformation, green), (iii) correctly folded loop (stem not in A-form, red), and (iv) left-handed Z-form stem (independent of loop conformation, grey), within all twelve continuous (demultiplexed) trajectories and the reference replica. Lower plots on each panel show fluctuations of major conformers in the reference replica over the course of REST2 simulation obtained by time-averaging over the 1 μs window (so that the value at 1 μs corresponds to the average over states populated between time of 0 to 1 μs). Displayed numbers indicate the final population of major conformers (in respective colors) in the unbiased (reference) replica (T = 298 K) calculated from the last 7 μs (see section Convergence of replica exchange simulations for details). See Methods in the main text for details about clustering procedure and Figure S7 for examples of 3D structures from selected simulations.

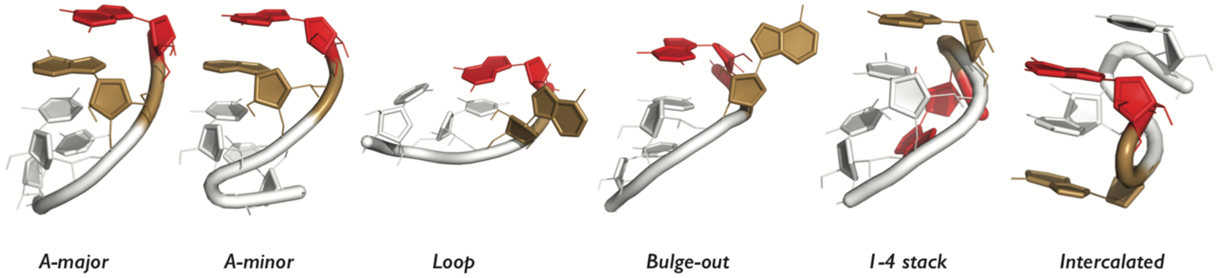

**Figure S6:** Tertiary structures of the most populated clusters from r(GACC) REST2 simulations. A, C, and G nucleotides are colored in sand, white, and red, respectively. H-atoms (except those from 2'-OH groups) are not shown for clarity.

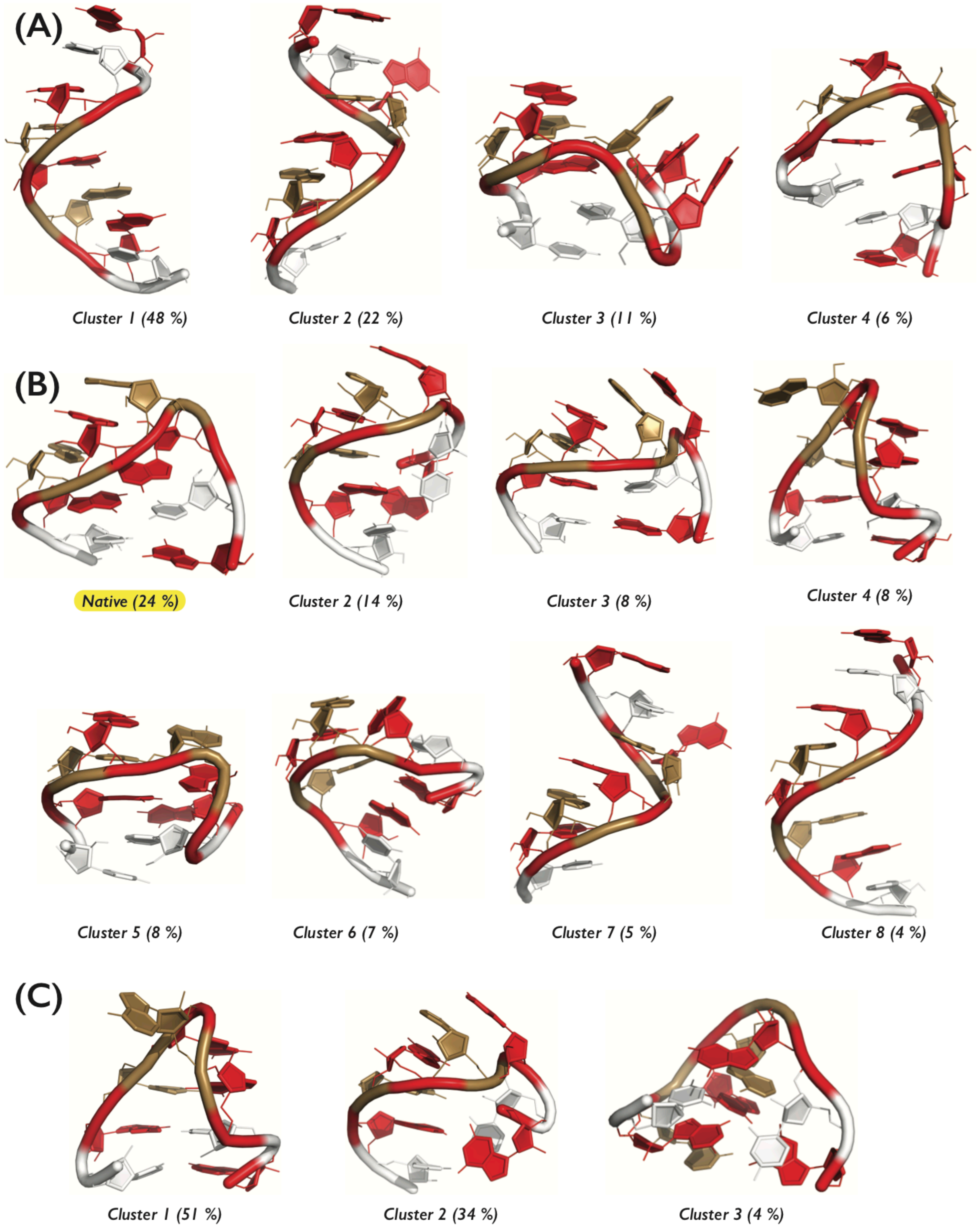

**Figure S7:** Tertiary structures of the most populated clusters from r(gcGAGAgc) REST2 simulations with (A) gHBfix$\binom{\text{2−OH…nbO/bO}}{-0.5}$, (B) gHBfix$\binom{\text{2−OH…nbO/bO}}{-0.5}\binom{\text{NH…N}}{+1.0}$, and (C) gHBfix$\binom{\text{2−OH…nbO/bO}}{-0.5}\binom{\text{NH…N}}{+1.0}\binom{\text{NH…O}}{+1.0}$. Only clusters with population higher than 4 % are shown. Nucleotides are colored as in Figure S6. H-atoms (except those from 2'-OH groups) are not shown for clarity. Native cluster that forms all signature interactions is highlighted in yellow and is sampled only in (B). Cluster 1 in (C) shows the structure with the left-handed Z-form helix conformation (stem guanines having *syn* orientation of the glycosidic bond).[12]

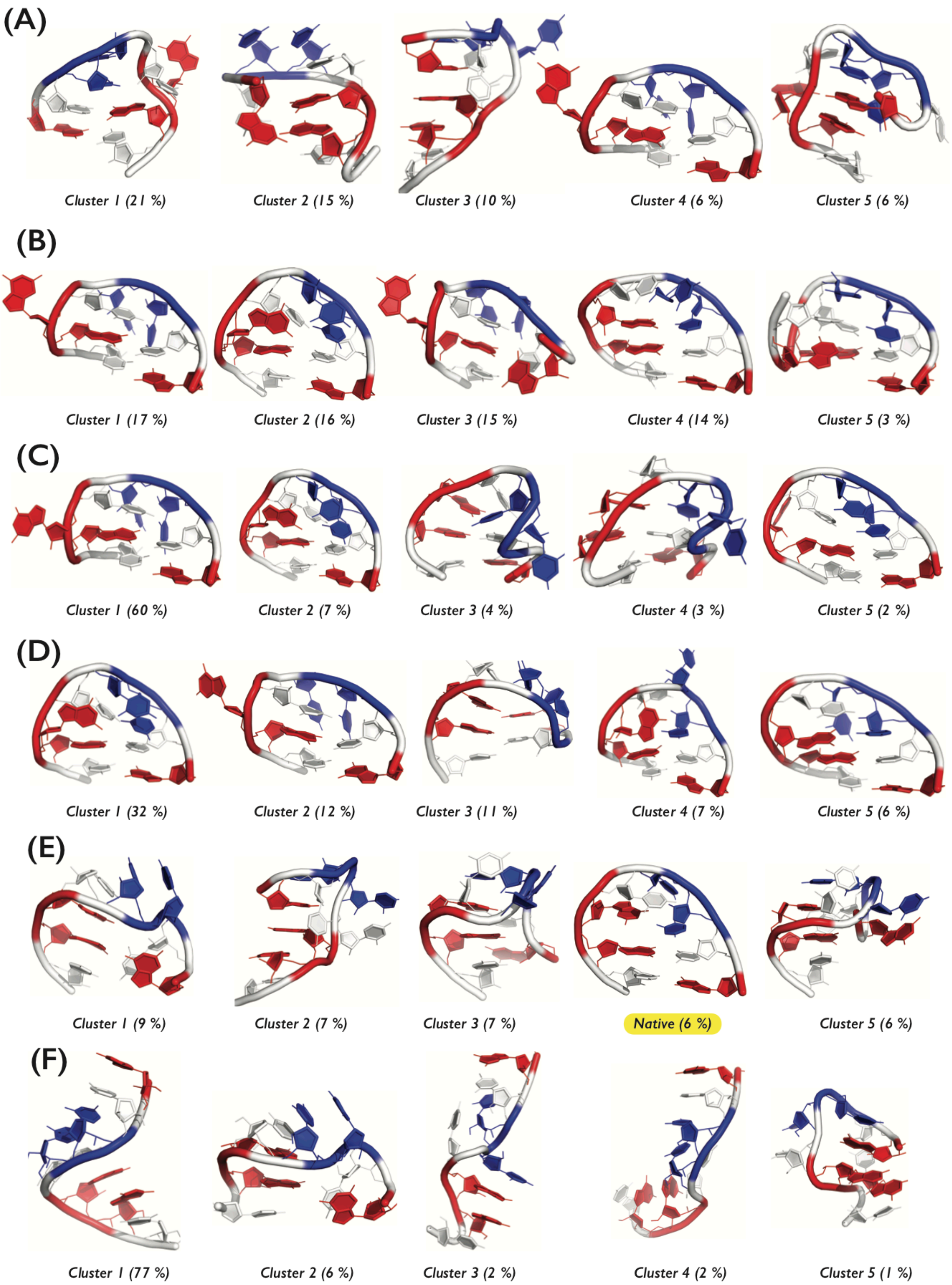

**Figure S8:** Tertiary structures of the most populated clusters from r(gcUUCGgc) REST2 simulations with (A) $\mathrm{gHBfix}\binom{2\text{–}\mathrm{OH...nbO/bO}}{-0.5}\binom{\mathrm{NH...N}}{+1.0}$, (B) $\mathrm{gHBfix}\binom{2\text{–}\mathrm{OH...nbO/bO}}{-0.5}\binom{\mathrm{NH...N}}{+1.0}\binom{\mathrm{NH...O}}{+1.0}$, (C) $\mathrm{gHBfix}\binom{2\text{–}\mathrm{OH...nbO/bO}}{-0.5}\binom{\mathrm{NH...N}}{+1.0}\binom{\mathrm{NH...O}}{+1.0}\binom{2\text{–}\mathrm{OH...N/O}}{+0.5}$, (D) $\mathrm{gHBfix}\binom{2\text{–}\mathrm{OH...nbO/bO}}{-0.5}\binom{\mathrm{NH...N}}{+1.0}\binom{\mathrm{NH...O}}{+1.0}\binom{\mathrm{NH...O2}}{-0.5}\binom{2\text{–}\mathrm{OH...O2/O4}}{-0.5}$, (E) $\mathrm{gHBfix}\binom{2\text{–}\mathrm{OH...nbO/bO}}{-0.5}\binom{\mathrm{NH...N}}{+0.5}\binom{\mathrm{NH...O}}{+0.5}\binom{2\text{–}\mathrm{OH...N/O}}{+0.5}\binom{\mathrm{NH...O2}}{-0.5}\binom{2\text{–}\mathrm{OH...O2/O4}}{-0.5}$, and (F) DESRES *ff*.

All REST2 simulation were started from unfolded states. Five highest populated clusters from each simulation are shown. A, C, G, and U nucleotides are colored in sand, white, red, and blue, respectively. H-atoms (except those from 2'-OH groups) are not shown for clarity. Native cluster that forms all signature interactions is highlighted in yellow and is sampled only in (E).

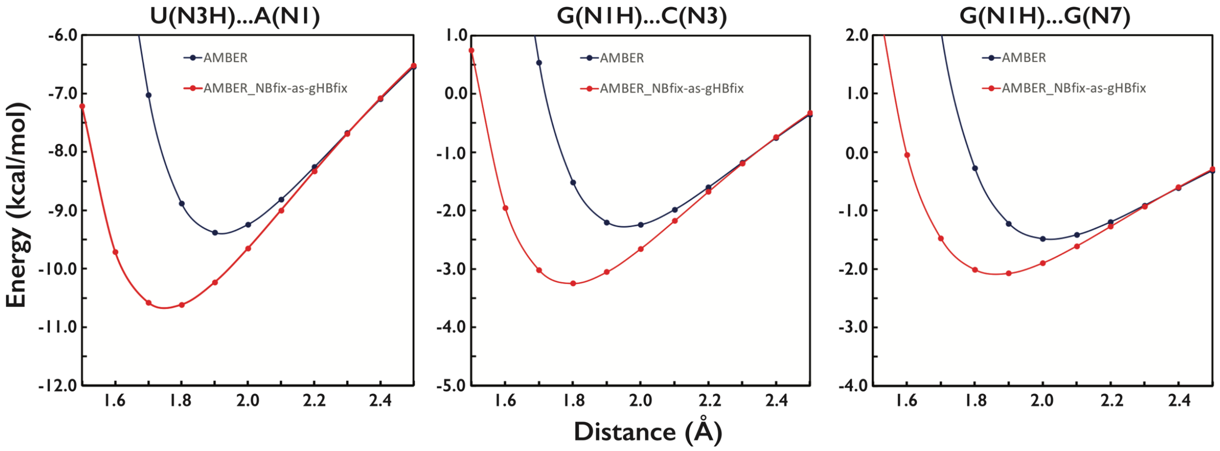

**Figure S9:** Interaction energies of H-bonds calculated by the standard $\chi_{OL3CP}$[6-9] AMBER *ff* (AMBER, blue) and the same *ff* modified by the NBfix approach (AMBER_NBfix-as-gHBfix, red). Plots show interaction energies of three different H-bonds, i.e., U(N3H)…A(N1), G(N1H)…C(N3), and G(N1H)…G(N7), which share the same NBfix parameters (for nitrogen and hydrogen atoms, Table S4) but are affected unequally due to different partial charges of those atoms. The NBfix parameters were prepared in a way to mimic the effect of the external gHBfix potential (Table S4).

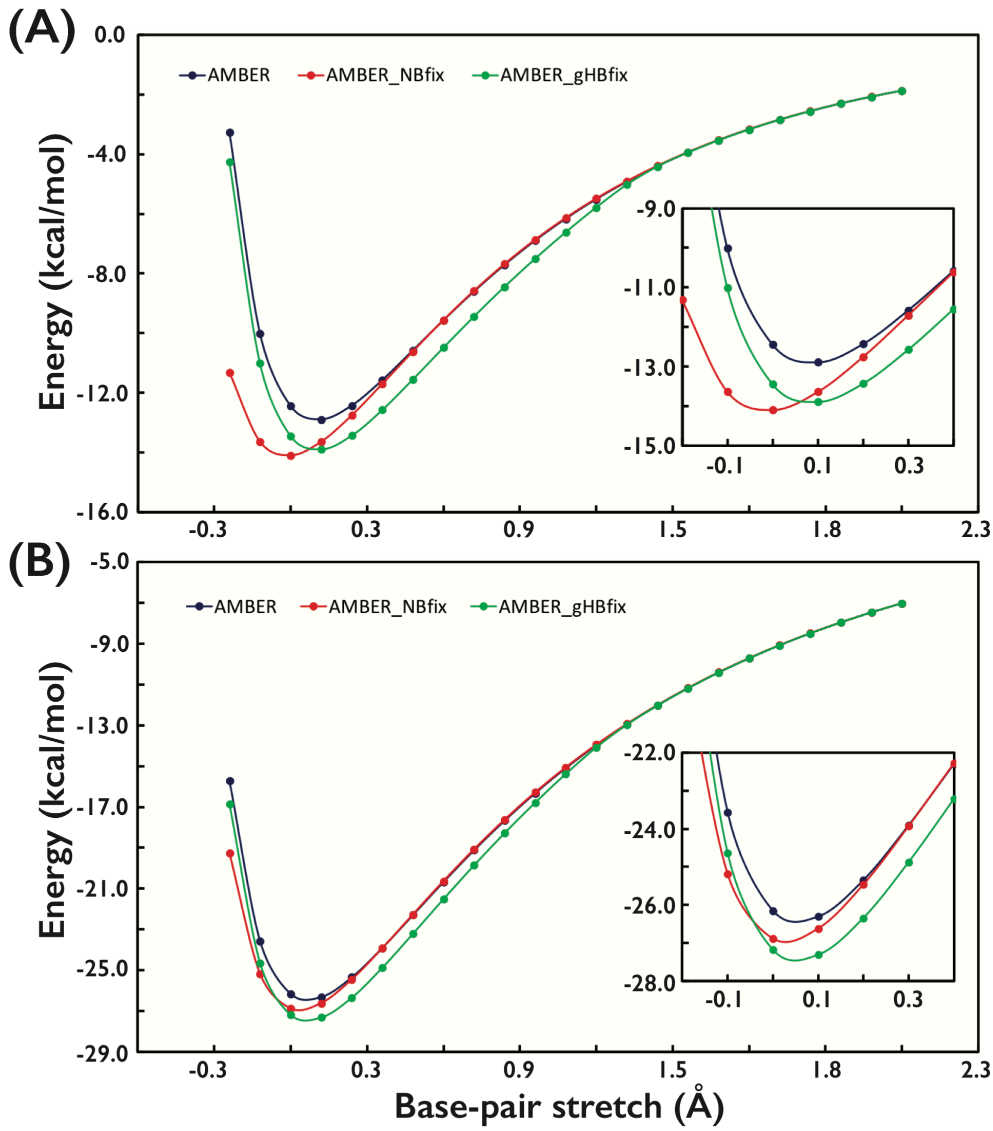

**Figure S10:** Interaction energies of adenine-thymine (A) and guanine-cytosine (B) Watson-Crick base pairs calculated as a function of the base-pair stretch (the region around the potential well is highlighted within the inset) in a 1-dimensional scan. The potential was calculated by three different *ff*s: (i) the standard $\chi_{OL3CP}$[6-9]AMBER *ff* (AMBER, blue), (ii) $\chi_{OL3CP}$ modified by the NBfix approach (AMBER_NBfix, red), and (iii) $\chi_{OL3CP}$ with the gHBfix potential (AMBER_gHBfix, green). The NBfix parameters were prepared in a way to mimic the effect of the external gHBfix potential (Table S4). The gHBfix approach was able to reproduce exactly the requested energy stabilization of 1.0 kcal/mol without affecting the position of stretch minima. The NBfix fit revealed non-uniform effect on the stabilization – see the main text for the explanation.

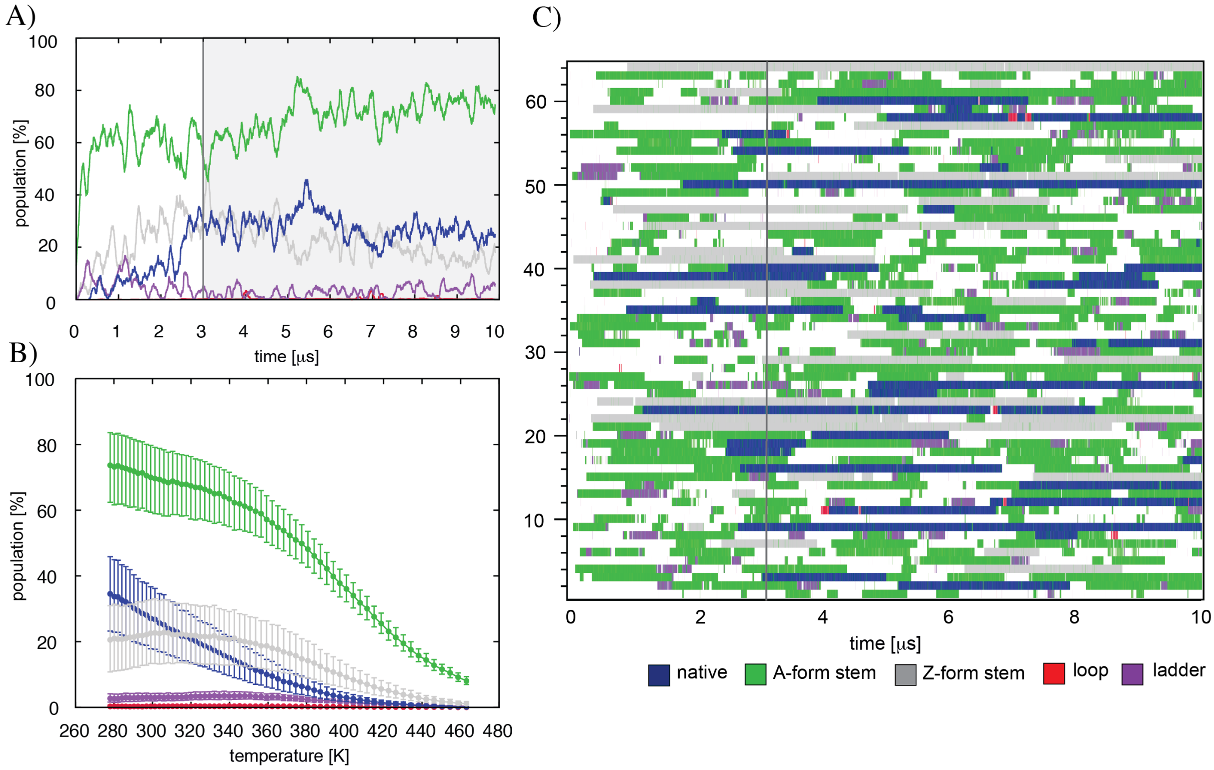

**Figure S11:** Conformational sampling and convergence of the r(gcUUCGgc) T-REMD folding simulation with structure-specific HBfix. (A) Population analysis (the 299 K replica) of the most important states, i.e., states with (i) correctly folded A-form stem irrespectively of the loop conformation (green), (ii) with correctly folded loop, where base pairs in the stem are not formed (red), (iii) native states with both stem and loop simultaneously folded with all signature interactions (blue), (iv) left-handed Z-form stem independent of loop conformation (grey), and (v) the spurious ladder-like structures[9, 27] characterized by the collective shift of the glycosidic torsions of all nucleotides from the *anti* to *high-anti* region (purple). The part in shadow starting with the grey line (3 μs) corresponds to the equilibrated part of T-REMD simulation. (B) Population of the most important conformers (calculated from the last 7 μs) as a function of the temperature. The error bars were estimated using bootstrapping with resampling both over time- and replica-domains (see Methods in the main text). (C) Time evolution of specific conformations within all 64 replicas analyzed from continuous trajectories. Note that the population of the ladder-like structure is entirely marginal, confirming again that the key $\chi_{OL3CP}$ dihedral reparameterization introduced in 2010 is entirely sufficient to suppress this spurious state. The coloring scheme is preserved within all panels; note that population of the stem state includes population of both stem with native loop (i.e., the true native state) as well as non-native loop conformations.

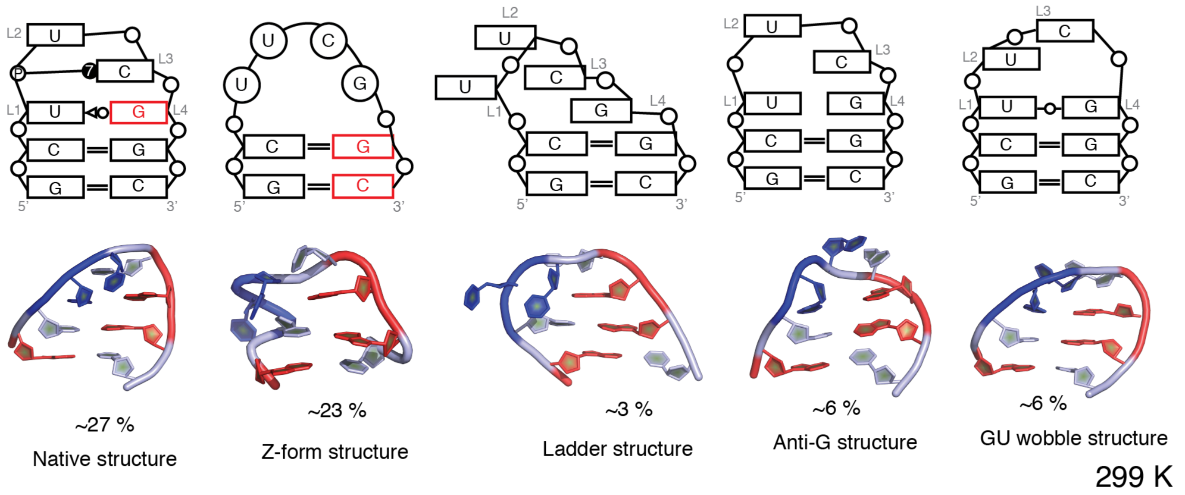

**Figure S12.** The secondary (top) and tertiary (bottom) structures of the most populated clusters (at 299 K) of the r(gcUUCGgc) TL taken from T-REMD folding simulation with structure-specific version of HBfix. The non-canonical base-pairing and BPh interactions are classified according to Leontis-Westhof-Zirbel nomenclature.[71-72] The bases in black correspond to *anti* orientation, while the bases in red correspond to *syn* orientation of the glycosidic bond. Tertiary structures have nucleotides colored as follows: G in red, U in blue and C in white. The used structure-specific bias is sufficient to achieve a substantial population of a fully correct native state.

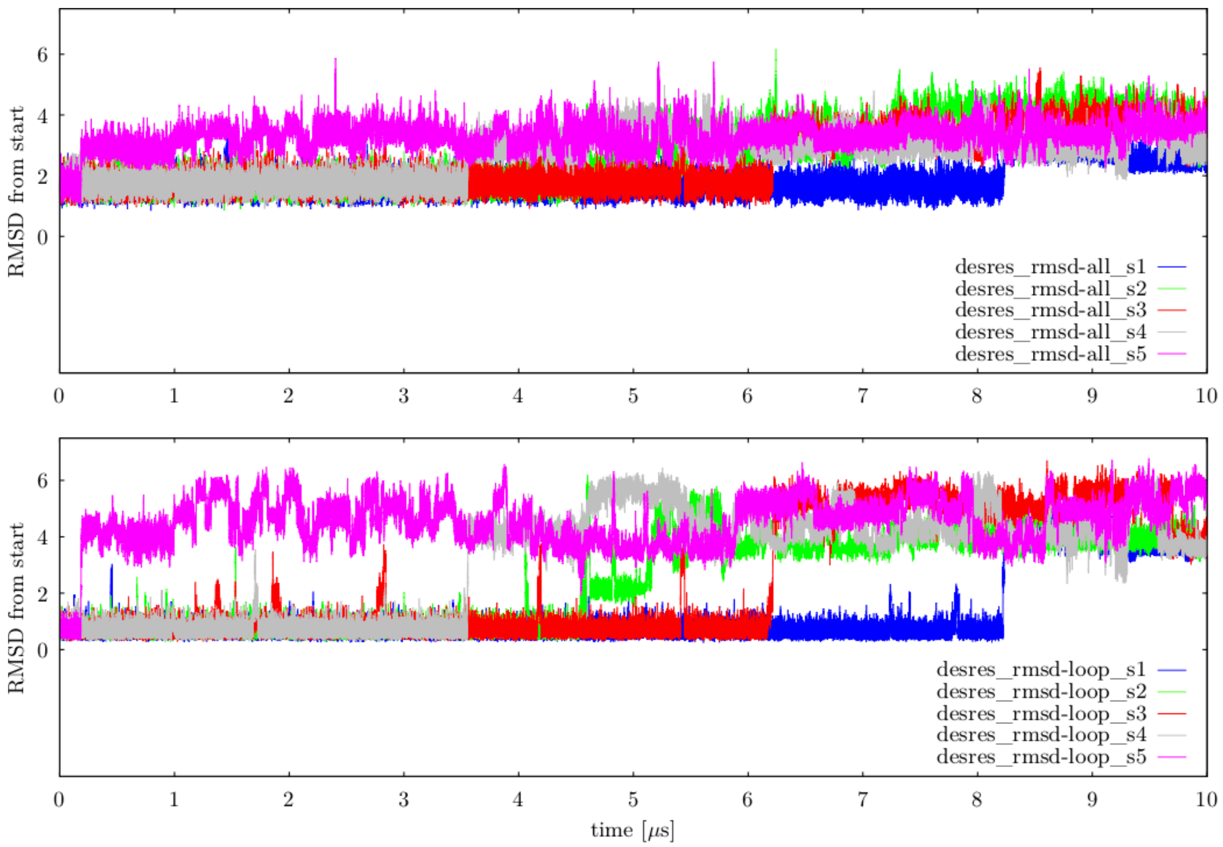

**Figure S13:** Evolution of RMSD in five independent unbiased MD simulations of 14-mer ggcacUUCGgugcc TL (PDB ID 2KOC) with DESRES potential.[13] The upper panel shows RMSD of heavy atoms of all nucleotides, while the lower panel displays corresponding RMSD of heavy atoms of only the four nucleotides forming the loop. The loss of the native TL arrangement in all five simulations is evident.

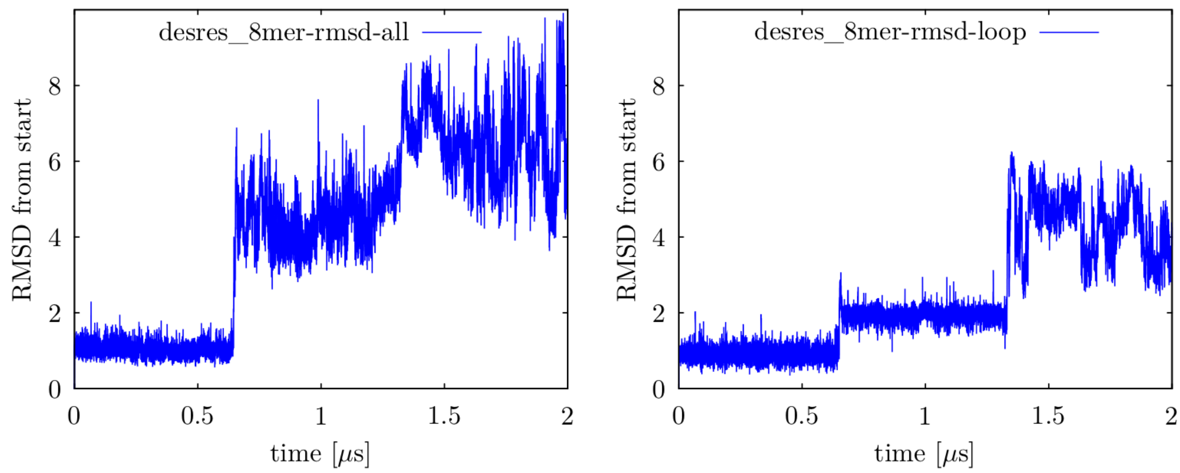

**Figure S14:** Evolution of RMSD in unbiased MD simulation of 8-mer r(gcUUCGgc) TL (PDB ID 3DVZ) using DESRES potential.[13] The left panel shows RMSD of heavy atoms of all nucleotides, while the panel on the right displays corresponding RMSD of heavy atoms of four nucleotides forming the loop. The unfolding of the TL motif is evident.

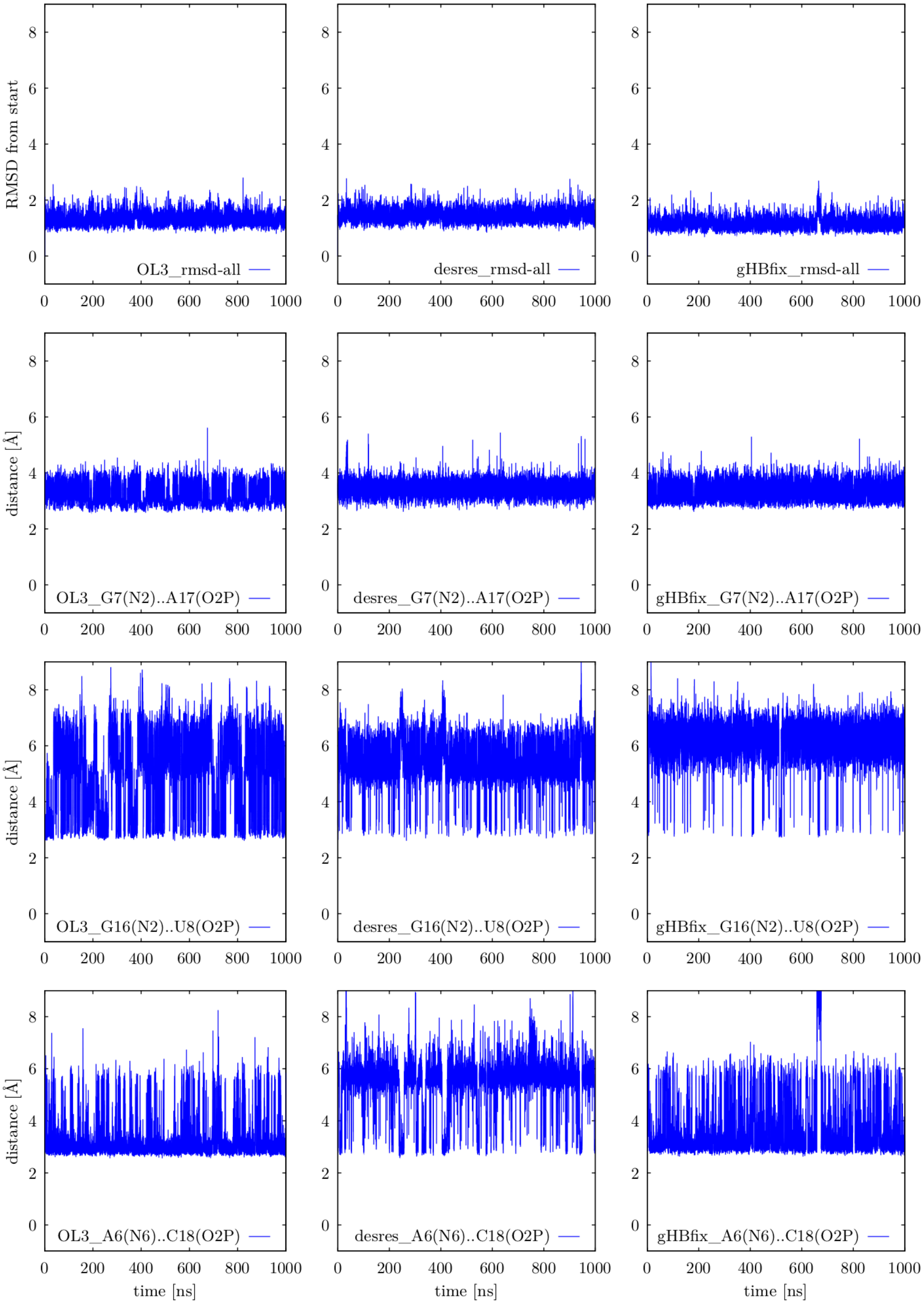

**Figure S15:** Structural analysis of three unbiased MD simulations of Sarcin-Ricin RNA loop (SRL) motif. See Figure 1 in Ref. [33] for the structure. Plots on the top are showing fluctuations of RMSD calculated over heavy atoms of all nucleotides during 1 µs-long unbiased MD simulations. Plots in the second, third and fourth row display fluctuations of distances indicating signature BPh contacts, i.e., G7(N2)…A17(*pro*-$R_P$), G16(N2)…U8(*pro*-$R_P$), and A6(N6)…C18(*pro*-$R_P$) distances, respectively. We compared the behavior in three different RNA *ff*s, i.e., the standard $\chi_{OL3CP}$[6-9] (OL3, plots on the left), DESRES potential[13] (desres, plots in the middle), and $\chi_{OL3CP}$ with the external gHBfix potential introduced in this paper (gHBfix, plots on the right, see Methods in the main text).

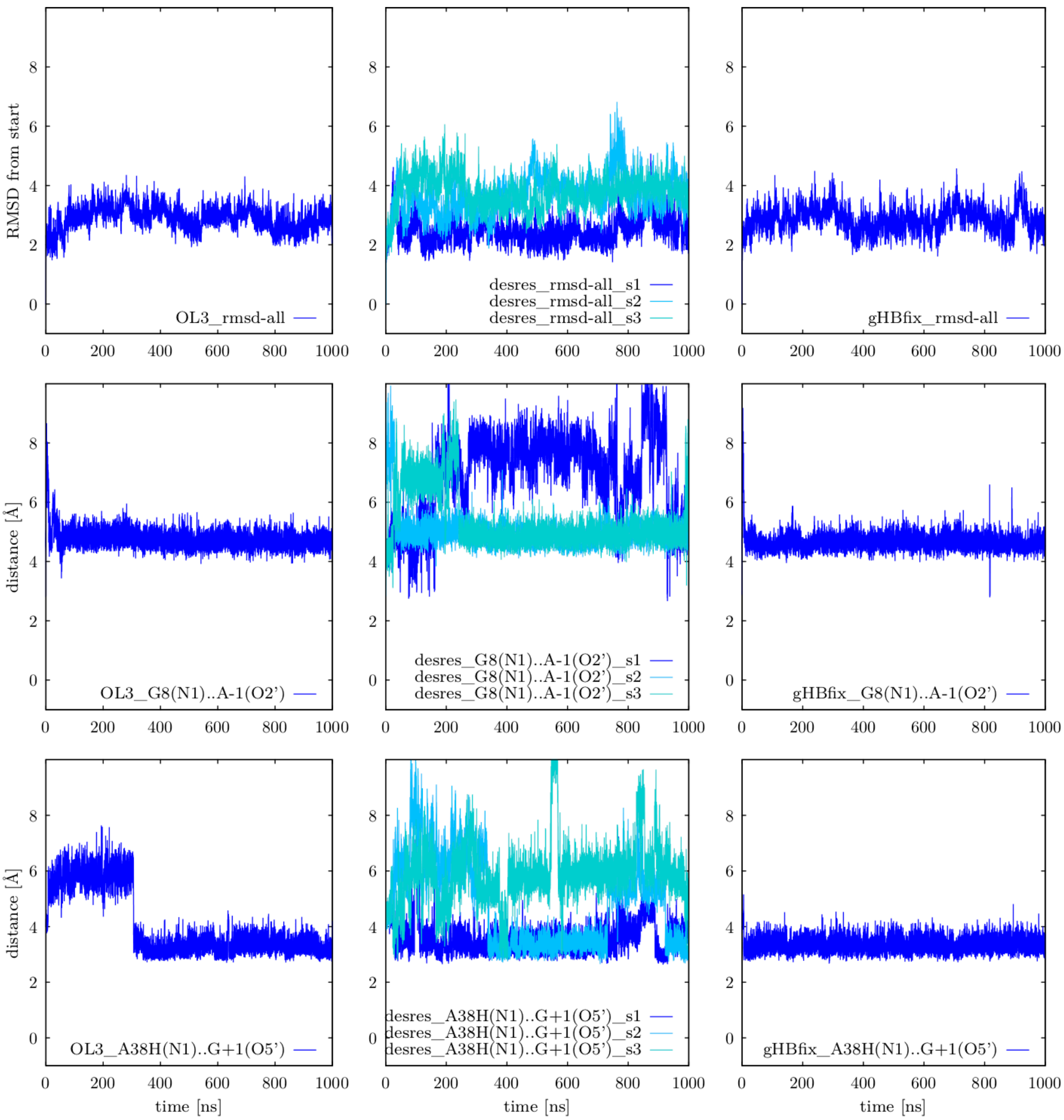

**Figure S16:** Structural analysis of unbiased MD simulations of the Hairpin Ribozyme (part I). Plots are showing fluctuations of (i) RMSD of heavy atoms calculated over all nucleotides, (ii) G8(N1)…A-1(O2'), and (iii) A38H$^+$(N1)…G+1(O5') catalytically important distances during 1 μs-long MD simulations. Data were taken from simulations in three different RNA *ff*s, i.e., $\chi_{OL3CP}$ (OL3, plots on the left), DESRES potential (desres, three independent simulations, plots in the middle), and $\chi_{OL3CP}$ with the gHBfix potential (gHBfix, plots on the right).

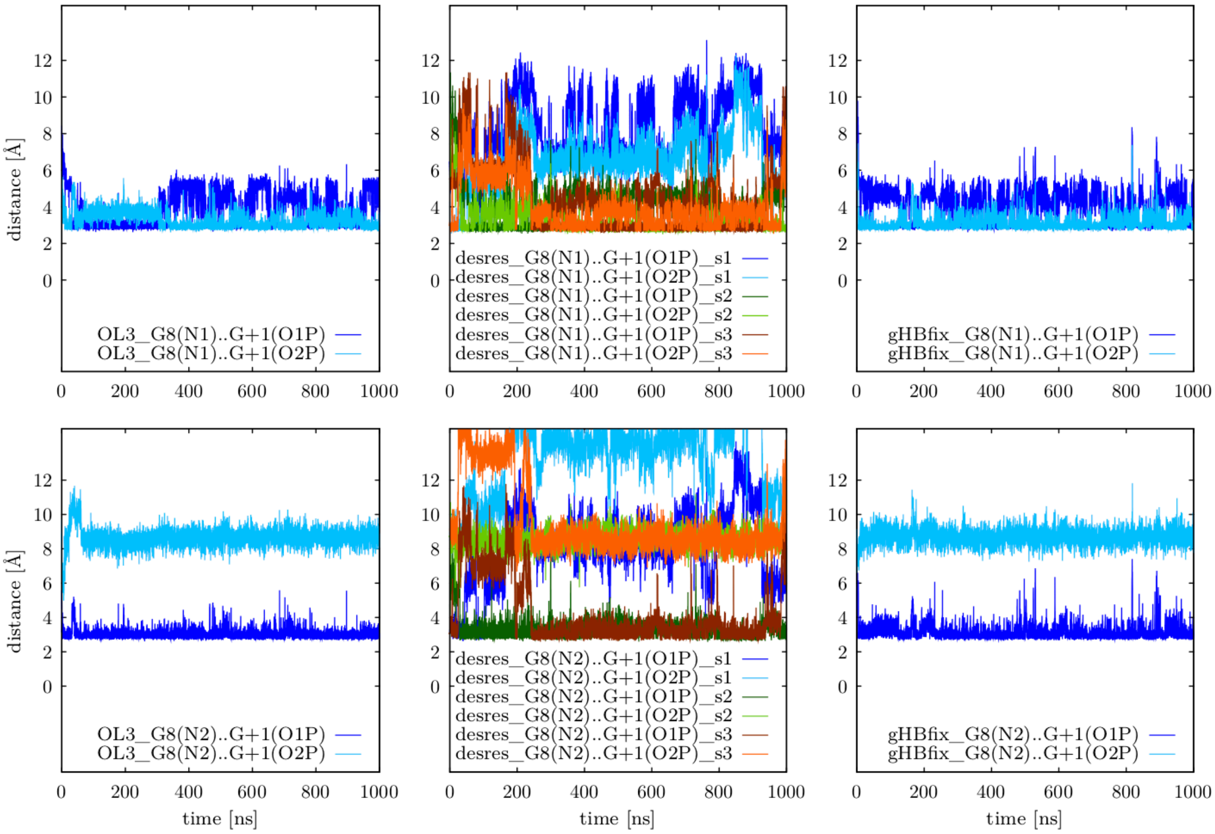

**Figure S17:** Structural analysis of three unbiased MD simulations of the Hairpin Ribozyme (part II), where we investigated the stability of the catalytically important BPh contact. Plots are showing fluctuations of distances between G8(N1/N2) atoms and both G+1(*pro*-$S_P$(O1P)/*pro*-$R_P$(O2P)) nbOs of the scissile phosphate. Data were taken from simulations in three different RNA *ff*s, i.e., $\chi_{OL3CP}$ (OL3, plots on the left), DESRES potential (desres, three independent simulations, plots in the middle), and $\chi_{OL3CP}$ with the gHBfix potential (gHBfix, plots on the right).

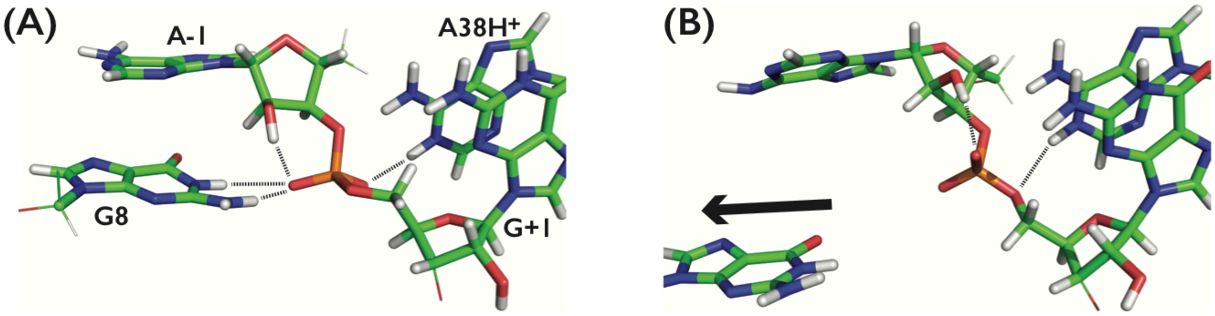

**Figure S18:** Close-up view into the active site of the Hairpin ribozyme from MD simulations with DESRES potential. (A) Snapshot from the beginning of the simulation, where both catalytically important canonical G8 and protonated $A38H^+$ residues established H-bonds (black dashed lines) with the scissile phosphate. (B) Illustrative snapshot, where G8 irreversibly lost its H-bonds with the scissile phosphate (G8(N1/N2)…G+1(*pro*-$S_P$/*pro*-$R_P$)) H-bonds were broken) and departed from the active site (black arrow).

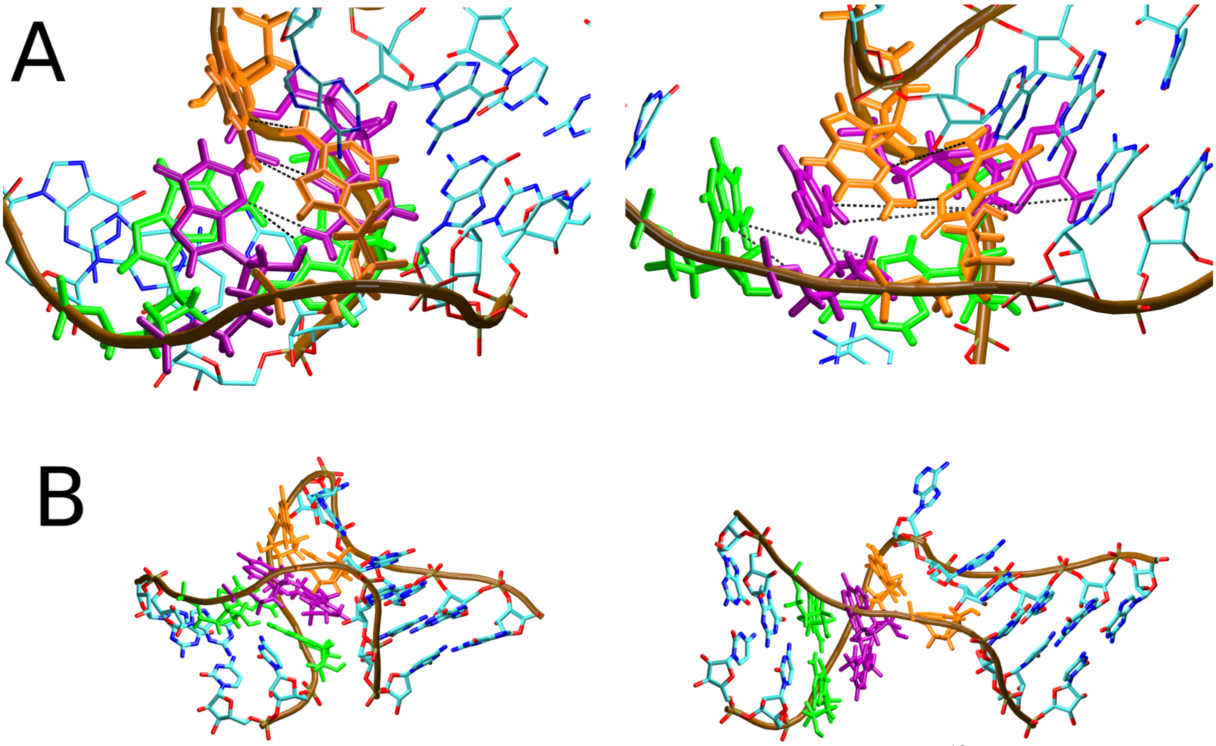

**Figure S19:** Snapshots from the standard MD simulation with DESRES[13] potential illustrate the inability of this *ff* to properly describe noncanonical interactions within the kink-turn Kt-7 RNA motif. (A) The sheared AG base pairs (left) revealed unstable behavior resulting in breakage of these pairs (right). The dashed black lines indicate the base pairing. The individual AG base pairs are highlighted in green, purple and orange, respectively. (B) The characteristic bent shape (left) of the kink-turn Kt-7 was lost on average timescale of ~200 ns in all attempted DESRES simulations (Table S2) and the structure became straightened (right). Similar behavior was observed in simulations with Pak-NBfix[66] RNA *ff*. Taken together, the structural changes observed in both these RNA *ff*s indicate a considerable overall bias in favor of a straight A-RNA.

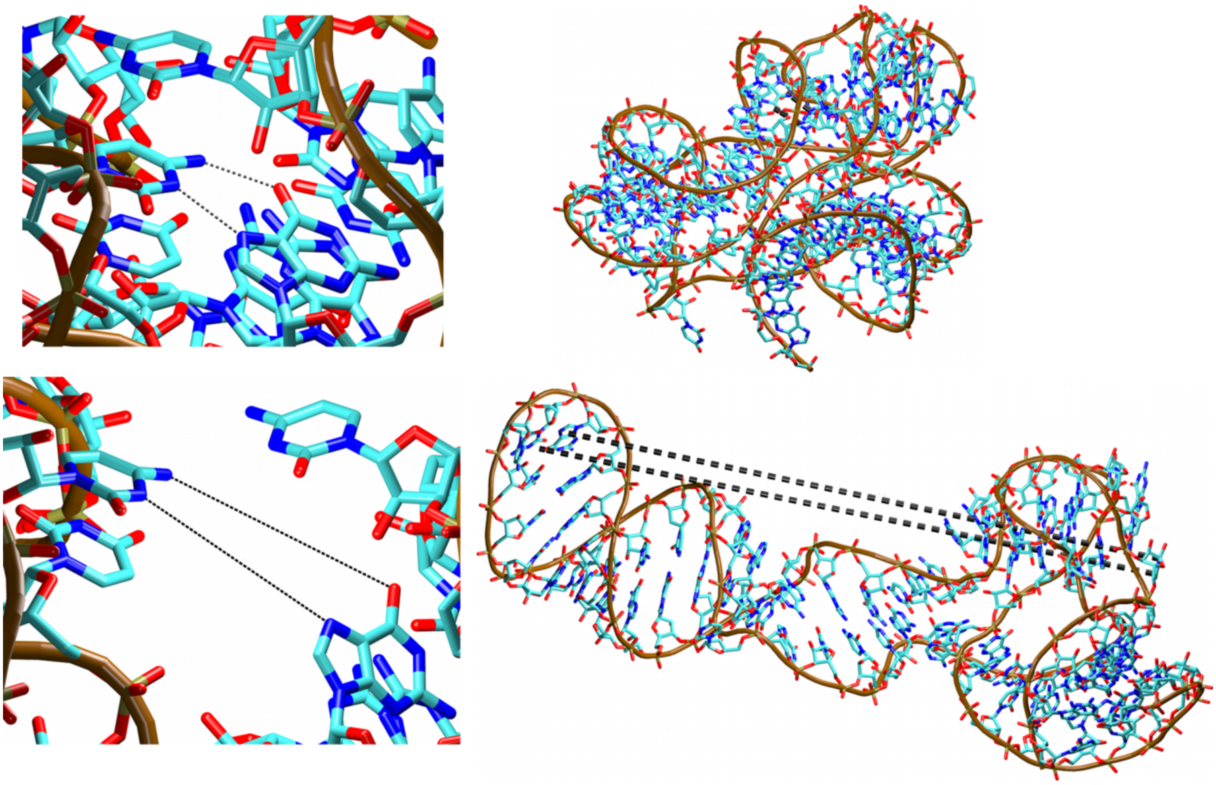

**Figure S20:** The loss of a key tertiary interaction between UNCG TL and an RNA internal loop in the simulations of the L1 stalk RNA with the DESRES potential (left) precipitates a large-scale degradation of the RNA fold (right). This degradation began immediately after the simulations started and fully progressed on timescale of ~500 ns. The black dashed lines mark the native tertiary interaction in the initial structure (top) and the loss of the interaction in the DESRES simulations (bottom). The same behavior has been seen in a number of independent simulations.
